# Residual dynamics resolves recurrent contributions to neural computation

**DOI:** 10.1101/2021.07.19.452951

**Authors:** Aniruddh R. Galgali, Maneesh Sahani, Valerio Mante

**Author notes:** Current Affiliation : Department of Experimental Psychology, University of Oxford, Oxford, United Kingdom (Aniruddh R. Galgali).

## Abstract

Relating neural activity to behavior requires an understanding of how neural computations arise from the coordinated dynamics of distributed, recurrently connected neural populations. However, inferring the nature of recurrent dynamics from partial recordings of a neural circuit presents significant challenges. Here, we show that some of these challenges can be overcome by a fine-grained analysis of the dynamics of neural residuals, i.e. trial-by-trial variability around the mean neural population trajectory for a given task condition. Residual dynamics in macaque pre-frontal cortex (PFC) in a saccade-based perceptual decision-making task reveals recurrent dynamics that is time-dependent, but consistently stable, and suggests that pronounced rotational structure in PFC trajectories during saccades is driven by inputs from upstream areas. The properties of residual dynamics restrict the possible contributions of PFC to decision-making and saccade generation, and suggest a path towards fully characterizing distributed neural computations with large-scale neural recordings and targeted causal perturbations.

## Introduction

Perception, decisions, and the resulting actions reflect neural computations implemented by large, interacting neuronal populations acting in concert^1,2^. Inferring the nature of these interactions from recordings of neural activity is a key step towards uncovering the neural computations underlying behavior^3–8^. One promising approach towards achieving this goal is based on the premise that neural computations are instantiated by a dynamical system^9,10^, emerging from an interaction between feed-forward inputs into a distributed neural population, and dynamics implemented through its recurrent connectivity^10–15^. The utility of such a “computation-through-dynamics” framework hinges on our ability to characterize the nature of this interaction and to disentangle the individual contributions of inputs and recurrent dynamics^6,16,17^.

Here, we investigate if and how the differential contributions of inputs and recurrent dynamics can be disentangled based on recordings of neural activity. We find that recordings from single areas are generally not sufficient to map these contributions onto specific anatomical structures. Nonetheless, key properties of inputs and recurrent dynamics can sometimes be revealed by analyzing the dynamical structure of neural population residuals, i.e. the trial-to-trial variability in neural population responses^18–23^. Our approach is based on the intuitive idea that the effect of recurrent computations can be revealed by observing how a perturbation of the state of the neural population evolves over time^24–27^. Unlike in causal perturbation experiments, where the perturbations are generated externally, we rely entirely on an analysis of recorded response residuals, which we interpret as naturally occurring perturbations within the repertoire of neural patterns produced by a recurrent neural network^28,29^. We term the time-varying dynamics of response residuals as “residual dynamics”, and show that it provides insights into the combined effects of the recurrent dynamics implemented locally, in the recorded area, and in upstream areas providing inputs to it. Obtaining a complete and quantitative description of residual dynamics is difficult, because the structured component of neural population residuals is typically dwarfed by unstructured noise that reflects variability in single-neuron spiking^18–20^. To obtain reliable and unbiased estimates of residual dynamics, we developed novel statistical methods based on subspace identification^30,31^ and instrumental variable regression^32^.

Our findings are organized in three sections. First, we illustrate the challenges in disentangling inputs and recurrent dynamics based on the simulations of simple dynamical system models (Fig. 1-2). These models are phenomenological, analogous to single-area, artificial recurrent neural networks (RNN) previously proposed for explaining the network-level mechanisms underlying sensory evidence integration^11,33^ and movement generation^12,34,35^. We use the simulations to establish what insights into recurrent dynamics can be obtained from different components of the neural responses, in particular conditioned-averaged responses and response residuals. Second, we study neural population recordings from pre-frontal cortex (PFC) of macaque monkeys during decision-making and saccadic choices (Fig. 3-5). While neural population trajectories in PFC are consistent with a number of previously proposed models of evidence integration and movement generation, we are able to rule out several candidate models based on the properties of the inferred residual dynamics. Third, to illustrate how inferred residual dynamics could be used to deduce circuit-level implementations of distributed recurrent computations, we consider simulations of more biologically realistic models, based on a previously proposed multi-area RNN of decision-making^36^ (Fig. 6-8).

**Figure 1.**
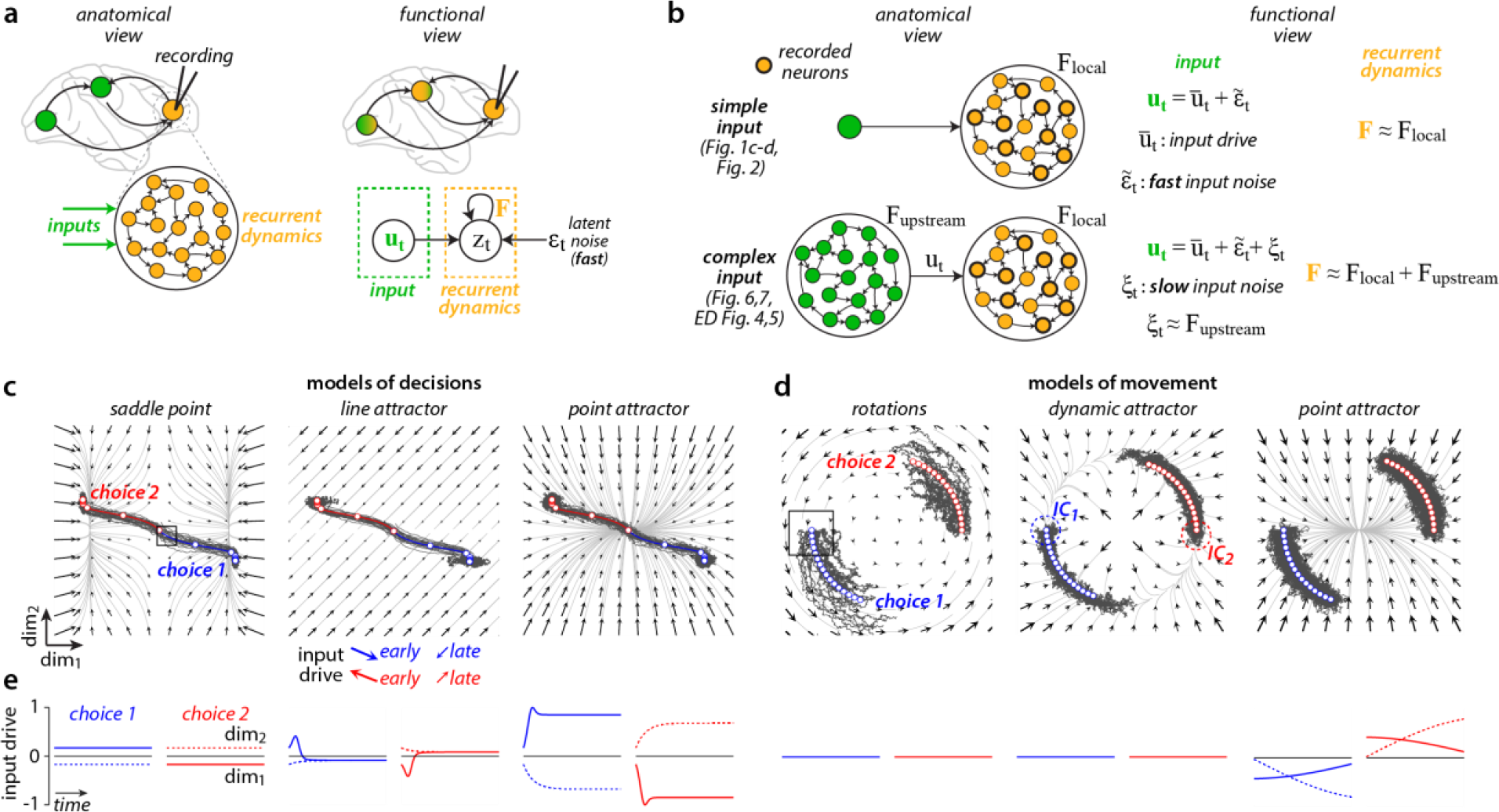
Challenges in disentangling contributions of inputs and recurrent dynamics from neural responses. **a**, Alternative views on computation through dynamics. Anatomical view (left): recurrent dynamics and inputs capture the influence of recurrent connectivity within the recorded area (orange) and from areas external to the recorded one (green) on the recorded neural responses. Functional view (right): recurrent dynamics and inputs reflect processes distributed across severa l areas (color gradient) and are defined based on their functional contributions to neural responses (graphical model, bottom). **b**, Relation of functional and anatomical viewpoints in two example scenarios (top & bottom row: simple vs. complex inputs). **c-d**, Models of decision-making (**c**) and movement generation (**d**) based on simple inputs as in **b** (top). Each panel shows simulated single trials (dark-gray trajectories) and condition-averaged trajectories (blue and red trajectories) for two task conditions (choice 1 and 2). Black arrows show the effect of the recurrent dynamics on the response at any location in state-space. The effect of the inputs is constant across state-space, but can change over time and across task conditions (middle panel in **c**, example input directions at bottom). **c**, Models of decision-making. Left: a model implementing a saddle point close to the initial conditions for both choice 1 and 2. Middle: a line attractor model. Right: a point-attractor model. The three models implement unstable (left), perfect (middle), and leaky integration (right) of an appropriately chosen input. **d**, Models of movement-generation. Left: purely rotational dynamics. Perturbations of the condition-averaged trajectory along both state-space dimensions are persistent; Middle: dynamic attractor model. Perturbations along the radial dimension decay, perturbations along the circular “channel” are persistent. Right: point attractor model. Responses are pushed away from the point attractor by strong inputs. IC: approximate extent of the initial conditions, shown as an example for the dynamic attractor model. **e**, Input drive (see **b**) for the models in **c** and **d**. Curves indicate the components of the input along the two state-space dimensions (solid vs dashed) as a function of time (horizontal axis) and condition (red vs blue). The input drives are chosen such that the different models of decision-making in **c**, and of movement-generation in **d**, cannot be distinguished based on the condition-averaged trajectories. Boxes in **c** and **d** (left sub-panels) show the regions of state-space analyzed in Fig. 2.

**Figure 2.**
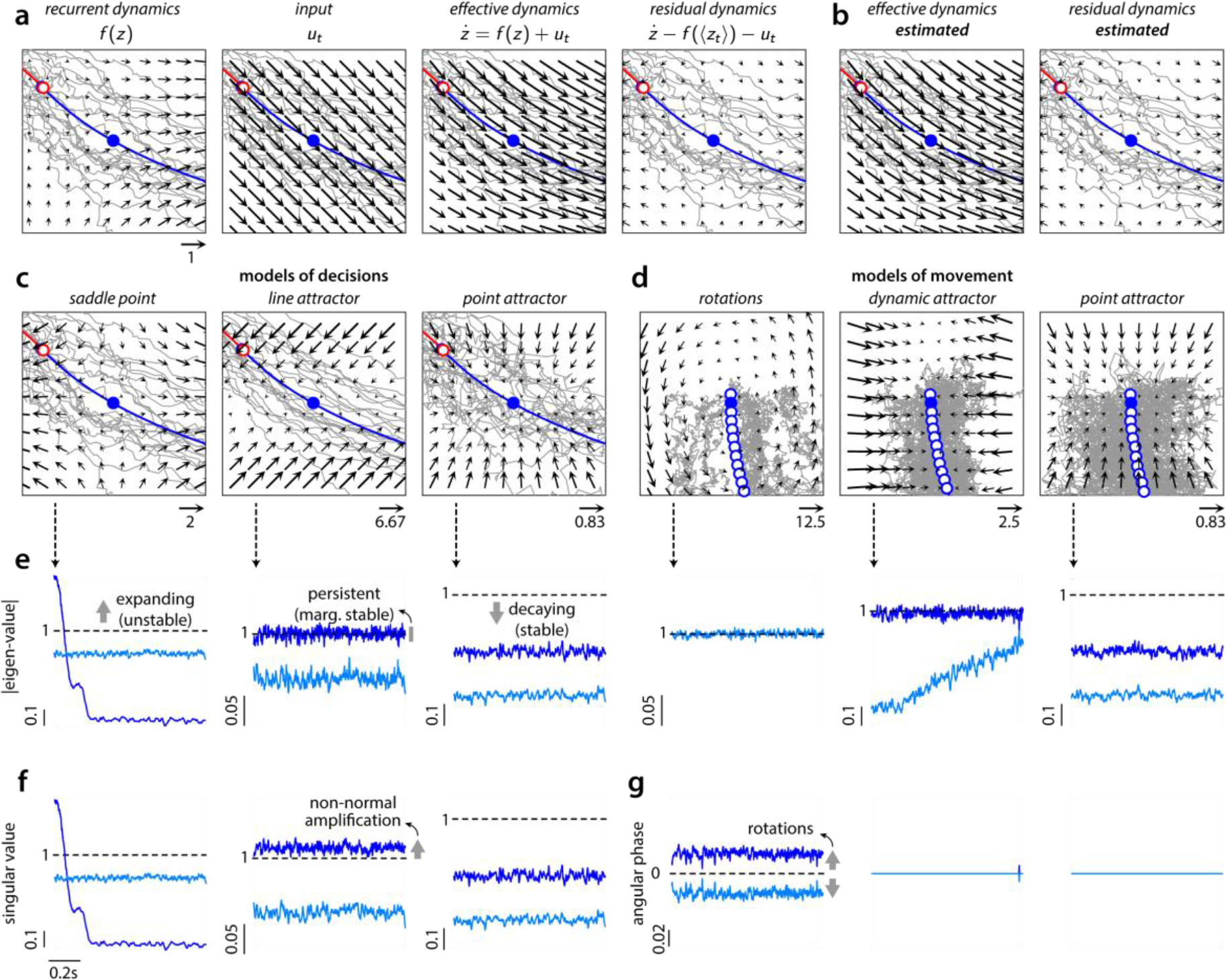
Residual dynamics reveals population-level computations. **a**, Different factors contributing to the dynamics of the saddle point model, shown in the state-space region marked in Fig. 1c for an early time in choice 1 trials (box). Same conventions as in Fig. 1c. Recurrent dynamics and input sum to generate the effective dynamics, determining the evolution of the response in the absence of noise. The residual dynamics is the component of the effective dynamics that explains the evolution of perturbations away from the condition-averaged trajectory (blue line; blue dot: reference time). **b**, Effective and residual dynamics estimated directly from simulated single-trial residuals match the ground-truth in **a**. **c**, Ground-truth residual dynamics for the models of decisions, same state-space region and reference time as in **a**. The residual dynamics reflects the key properties of the recurrent dynamics at the corresponding state-space region in Fig. 1c. The arrows in each flow field were scaled by a fixed factor that differed across models and with **a** (numbers close to arrows at the bottom). **d**, Analogous to **c**, but for the models of movement at an early time in choice 1 trials (box in Fig. 1d). **e-g**, Properties of the estimated residual dynamics for the models in Fig. 1c-d. Only residual dynamics for choice 1 is shown. The residual dynamics is described by a time and condition-dependent, autonomous, linear dynamical system. The corresponding time-varying dynamics matrices describe the residual dynamics at particular locations along one of the condition-averaged trajectories (Extended Data Fig. 1). **e**, Magnitude of the eigen-values (EV, y-axis) of the 2-dimensional dynamics matrix as a function of time (x-axis). **f**, Singular values (SV) of the dynamics matrix as a function of time for the models of decisions. The difference between EV and SV in the line-attractor model is a consequence of non-normal dynamics. **g**, Angular phase associated with complex-valued EV for models of movement. Larger angular phase implies faster rotational dynamics. EV, SV, and angular phase together distinguish between the different models (Extended Data Fig 10).

**Figure 3.**
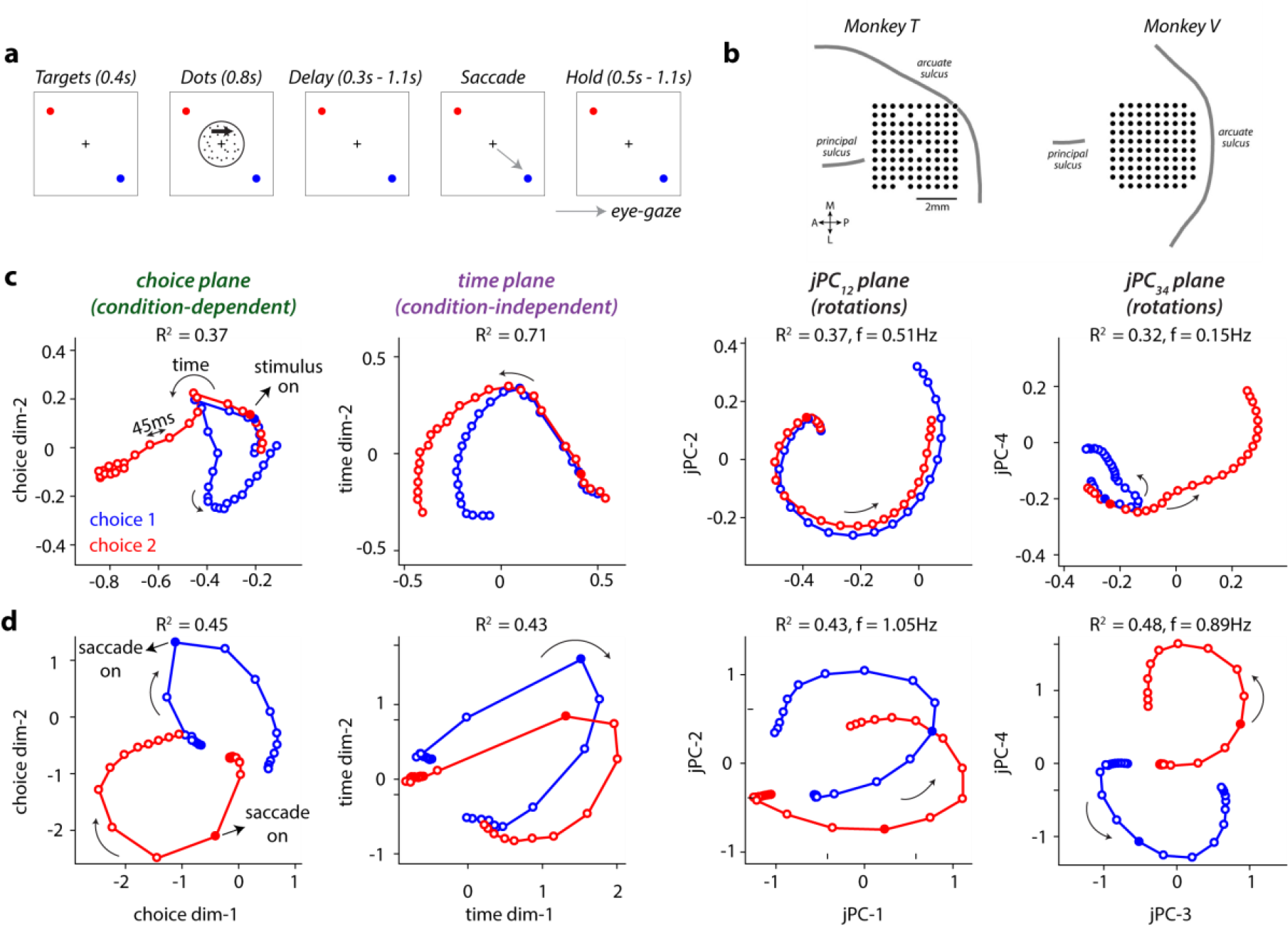
Average neural dynamics in prefrontal cortex during perceptual decisions and saccades. **a**, Behavioral task. Monkeys fixating at the center of a screen (fixation point, black cross) viewed a random dot stimulus for 800ms. After a delay period of random duration, they reported the perceived direction of motion with a saccade to one of two targets (red and blue circles; blue: choice 1; red: choice 2). Following the saccade, the monkeys had to fixate on the chosen target during a hold period of random duration. **b**, Position of the 10 × 10 electrode array in pre-arcuate cortex of the two monkeys. Black circles indicate the cortical locations of the 96 electrodes used for recordings. **c-d**, Neural trajectories in monkey T, averaged over trials of the same choice. Trajectories are obtained after aligning neural responses (see Extended Data Fig. 6) from experimental sessions with a similar configuration of saccade targets (config-3, Extended Data Fig. 6). Aligned responses are projected into four activity-subspaces: the choice, time, jPC_12_, and jPC_34_ planes, capturing variance due to choice, time, and rotations, respectively (R^2^: fraction of variance explained; f: rotation frequency associated with the jPC plane). **c**, Trajectories in the decision-epoch (−0.2 to 1s relative to stimulus onset, filled circle). **d**, Trajectories in the movement-epoch (−0.7 to 0.5s relative to saccade onset, filled circle).

## Results

In the framework of computation through dynamics, the temporal evolution of the state of a neural population (**z**_***t***_, *t* indicates time) can be described through a differential equation:

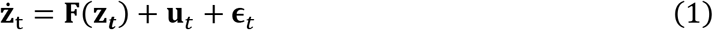

The momentary change in the population state (ż_t_) on each trial reflects the combined effect of four distinct factors, namely the *recurrent dynamics* (**F**(**z**_***t***_)), the *inputs* (**u**_*t*_), the *latent noise* (**ϵ**_*t*_), and the initial conditions (**z**_0_, the state at time zero). The first three factors are assumed to combine additively, as is approximately the case in many neural network models^11–14^.

A natural interpretation of Equation 1 would be to map these factors directly onto individual brain areas (Fig 1a, anatomical view). However, below we show that inferring such a mapping is typically not possible when using neural recordings from only one or few areas within a larger network^17,37^. We thus pursue a more abstract interpretation, whereby the state **z**_***t***_ amounts to a low-dimensional approximation of the activity of all recorded neurons^29^, and each factor can be distributed across many areas^38^ (Fig 1a, functional view). Under this interpretation, the factors in Equation 1 can sometimes be distinguished at a functional level, because they contribute distinctly to patterns of variability in neural responses. Specifically, **F**(**z**_***t***_) captures the functional consequences of distributed recurrent connectivity in driving the low-dimensional dynamics of the population and induces variability over slow time-scales (i.e. long autocorrelation over time), **ϵ**_*t*_ captures fast variability (no autocorrelation), and **u**_*t*_ can capture either fast or slow variability, depending on the complexity of processing in areas upstream of the recorded one (Fig 1b).

We illustrate the relation between the anatomical and functional interpretations by considering two idealized scenarios, which differ in the functional complexity of the inputs. We consider simulations based on “simple” inputs of a purely feed-forward nature (Fig 1b, top; Fig. 1c-d, 2) or more “complex” inputs resulting from recurrent processing that is upstream of the recorded area (Fig 1b, bottom; Extended Data Fig 4,5). These simulations illustrate the challenges in distinguishing the functional contributions of recurrent dynamics and inputs based on partial neural recordings, but also that response residuals are better suited to this challenge than the full neural population trajectories.

**Figure 4.**
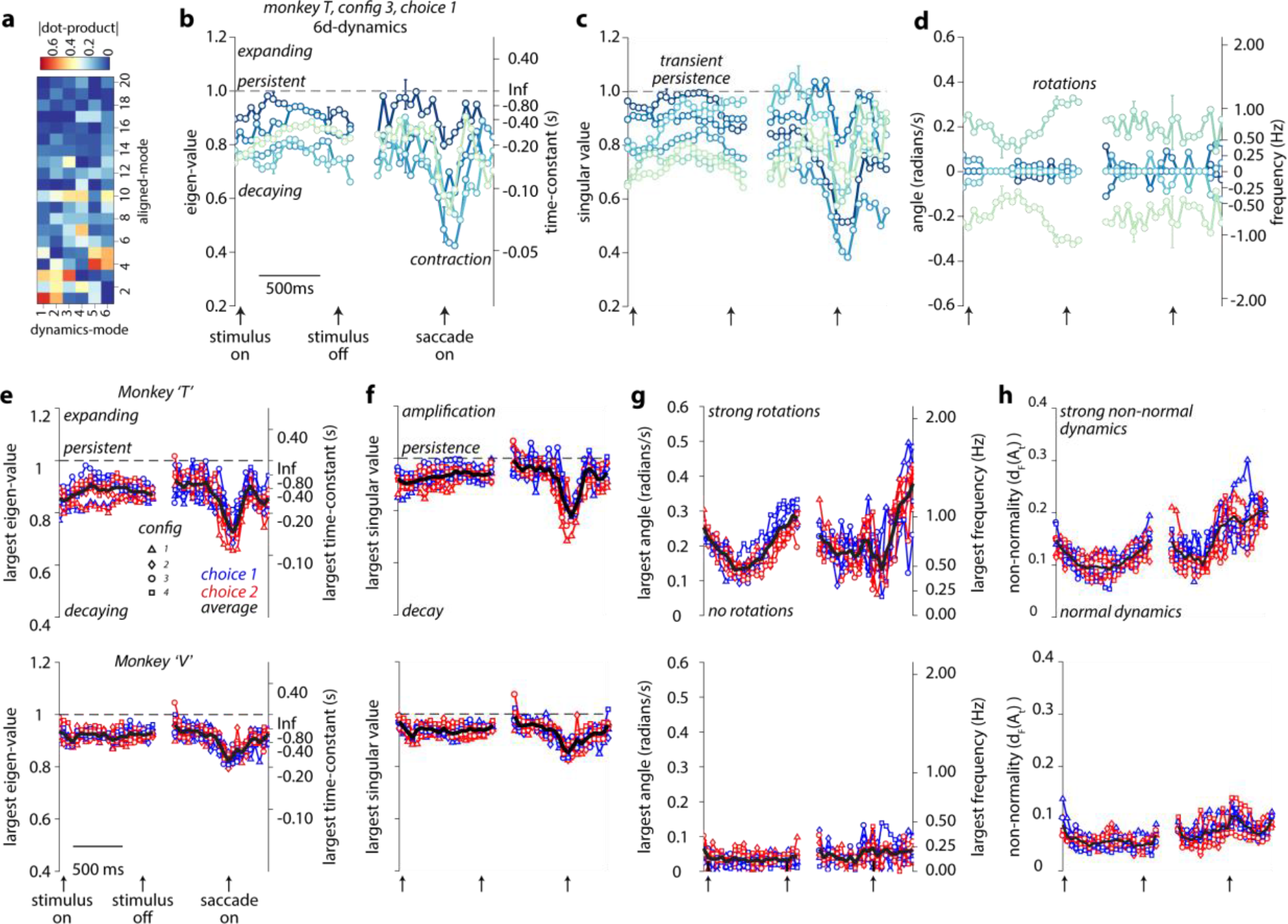
Residual dynamics in prefrontal cortex during perceptual decisions and saccades. **a-d**, Estimated residual dynamics in prefrontal cortex in monkey T, same task configuration as in Fig 3c,d. The residual dynamics was 8-dimensional for this example dataset. **a**, Relative alignment between the modes spanning the 8d-dynamics subspace and the modes spanning the 20d-aligned subspace (see Extended Data Figs. 6,7), measured as the absolute value of the corresponding dot-product. The dynamics modes project strongly onto the first few aligned modes, which capture most of the task-relevant variance in the responses. **b-d**, Properties of the residual dynamics for a single choice condition (choice 1). Error bars: 95% bootstrap confidence intervals (shown at selected times). **b**, Eigen-values (EV) of the dynamics (left axis), and associated time-constants of decay (right axis) as a function of time (x-axis). **c**, Singular values (SV) of the dynamics. The eigenvectors and singular vectors associated with the shown EV and SV can vary over time. **d**, Angular phase of the EV (left axis; angular phase = 0: real-valued EV) and associated rotation frequencies (right axis). Line colors reflect the magnitude of the EV or SV at the first time of the decision epoch. At later times, colors match those associated with the closest eigen-vector or right singular vector at the previous time. **e-h**, Properties of the residual dynamics across all animals (Monkey T, top; Monkey V, bottom), choices (blue: choice 1; red: choice 2), and task configurations (markers; see legend of Extended Data Fig. 6). Black curves: averages across all choices and configurations. **e**, Magnitude of the largest EV (left axis) and the associated decay time-constants (right axis). **f**, Largest singular value. **g**, Largest angular phase of the EV and the corresponding frequency of rotation. **h**, Time course of the index of non-normality.

**Figure 5.**
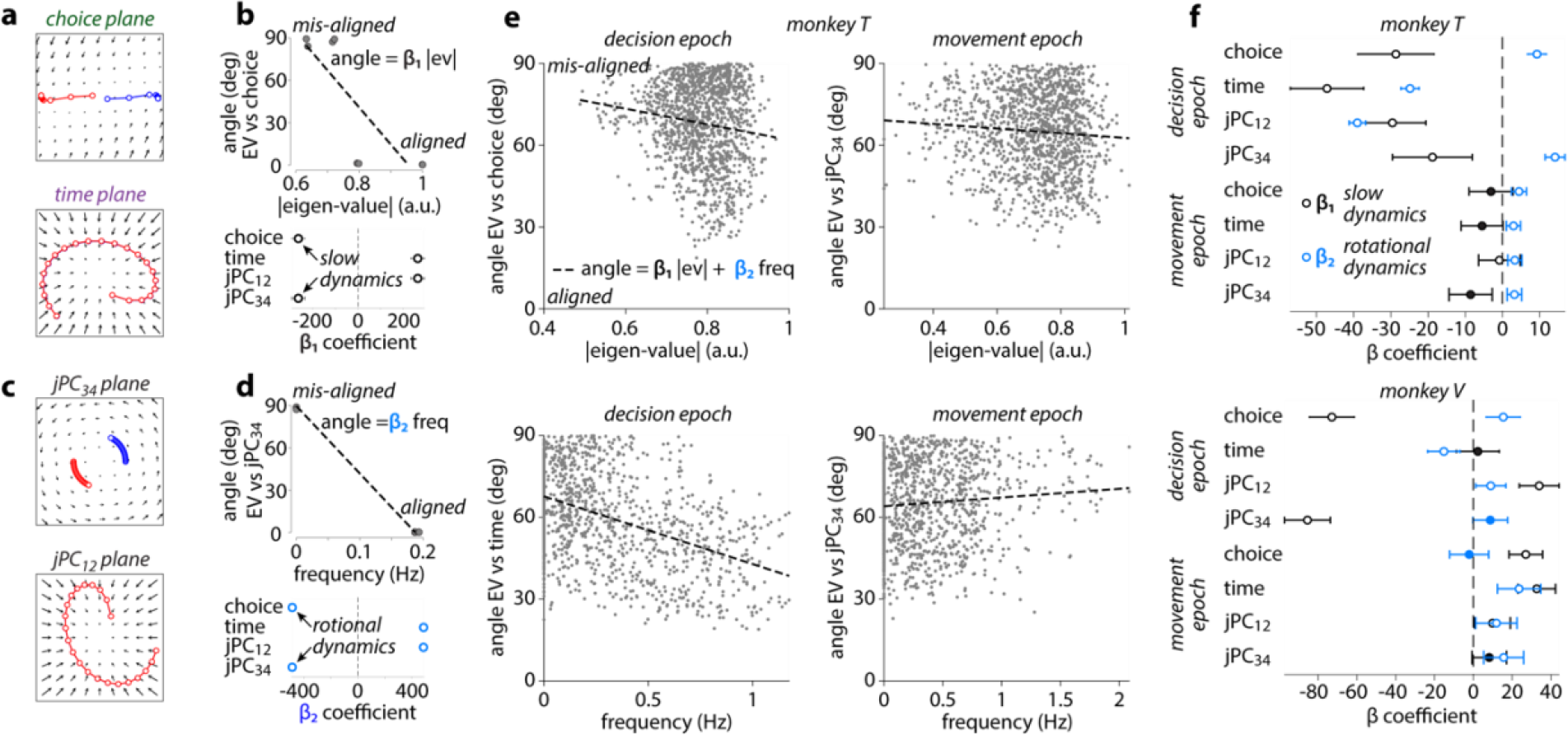
Alignment of residual dynamics and condition-averaged trajectories. **a**, Condition-averaged trajectories for the line attractor model (top; and Fig. 1c) augmented with an additional two-dimensional subspace with decaying dynamics and strong input drive (bottom, time plane; red and blue trajectories are overlayed). **b**, Alignment between the eigenvectors of the residual dynamics and the task-related subspaces, for the model in **a**. Top: Angle between the choice plane and the eigenvectors (gray points). Eigenvectors are indexed by EV magnitude. Bottom: regression coefficients, for linear regression as on top (line; angle vs. EV magnitude). Large negative coefficients identify task-subspaces aligned with slow residual dynamics. Task-subspaces are redundant (e.g. choice and jPC_34_) as residual dynamics is only 4-dimensional. **c**, Trajectories for the rotation model (top; and Fig. 1d) augmented as in **a** (bottom, jPC_12_ plane). **d**, like **b**, for the model in **c**. Eigenvectors are indexed by the associated rotation frequency. Large negative coefficients identify task-subspaces aligned with rotational residual dynamics. **e**, Example alignments for PFC activity in monkey T. Angles (gray points) are pooled across times within an epoch (titles), task configurations, and choices. Linear regression (dashed line) includes coefficients (β_1_ and β_2_) for EV magnitude (|ev|) and rotation frequency (freq). **f**, Regression coefficients for PFC activity (as in **e**), for all epochs, task-subspaces, and monkeys. Error bars indicate 95% normal confidence intervals; filled circles indicate non-significant regression coefficients (p>0.05).

**Figure 6.**
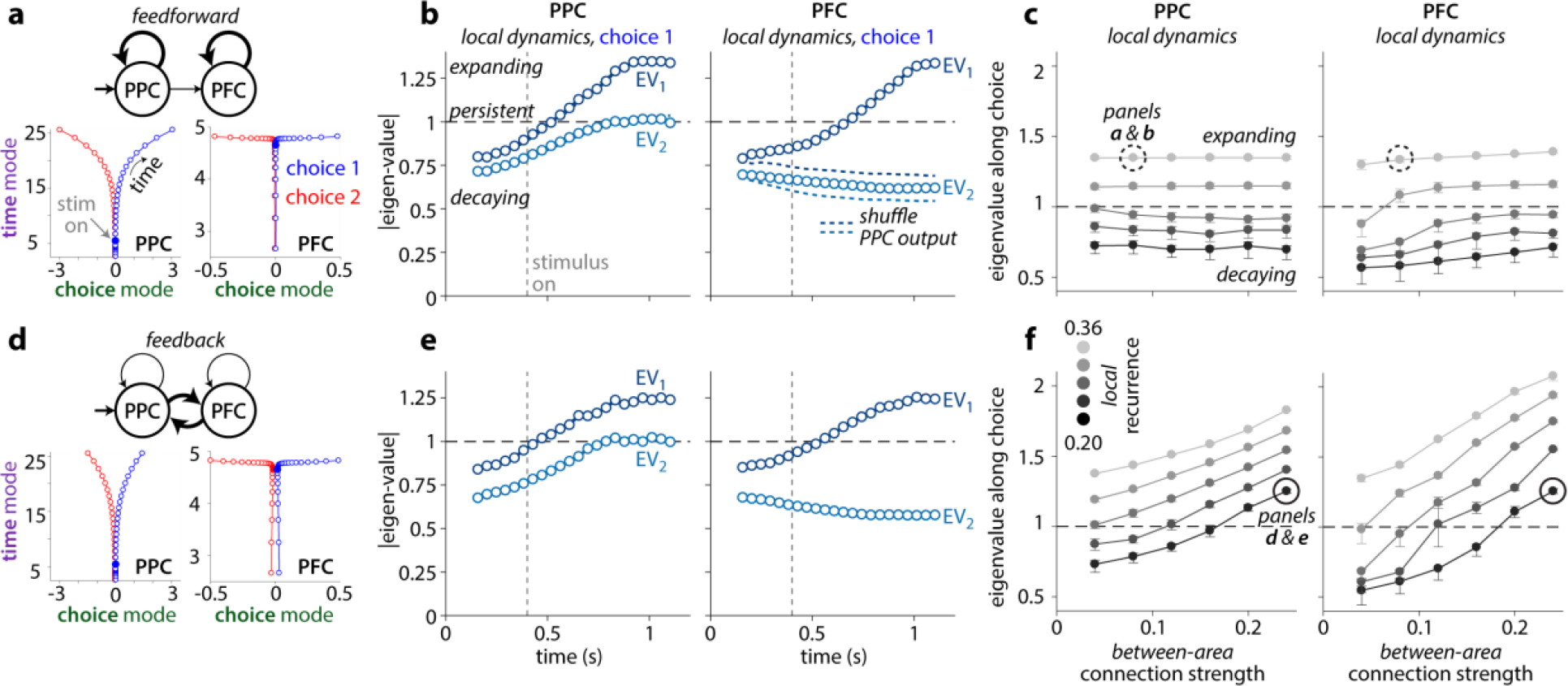
Local residual dynamics in multi-area networks of perceptual decision making. Each network consists of two interconnected modules (PPC and PFC), whereby a module mimics an RNN with a given level of local recurrence. PPC is driven by an external input, and feedback connections from PFC to PPC are either absent (**a-c**) or present (**d-f**). **a**, Connectivity (top) and average trajectories (bottom) for an example network with weak feedforward connectivity between areas (top, thin arrow) and strong local recurrence (thin arrows). Condition-averaged trajectories are shown separately for each area for two choices (blue: choice 1, red: choice 2). Trajectories are visualized in a subspace spanned by the choice mode, explaining variance due to choice, and a time mode, explaining condition-independent variance. **b**, Time-varying EV magnitude of the local residual dynamics estimated from residuals in PPC (left) or PFC (right) for choice 1, in the example network in **a**. The external input is turned on 400ms after the start of the trial (gray dashed line). EV magnitudes in PFC are strongly reduced upon shuffling the feedforward output of PPC across trials (blue dashed curves). **c**, Maximum EV magnitude (measured across time) for residuals projected onto the choice modes in PPC (left) or PFC (right), as a function of the strengths of local recurrence (black to gray: small to large recurrence) and between-area connections (x-axis). Errorbars indicate 95 percentile bootstrap confidence intervals. The dashed circle marks the example network shown in **a-b**. **d-f**, Same conventions as in **a-c**, but for networks with between-area feedback.

### Neural population trajectories poorly constrain recurrent computations

We simulated responses from a number of distinct models that explain neural population dynamics during perceptual decision-making and movement generation (Fig. 1c-d). While these hand-designed models are not meant to precisely reproduce neural recordings, they do capture the distinctive features of rather complex, non-linear RNNs trained to integrate sensory evidence towards a choice11,33 (Fig. 1c) or generate complex motor sequences^12,35^ (Fig. 1d). As is common in these RNNs, the input is defined as an additive combination of two components (Fig. 1b, functional view): the input drive ū_*t*_ which is deterministic, i.e. repeatable across trials of the same condition; and input noise 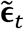 analogous to the latent noise (Fig. 1b, simple inputs).

We simulated single-trial responses for two task-conditions and represented them as trajectories in a 2-dimensional neural state-space (Fig. 1c,d, choice 1 & 2; dark-gray curves). The recurrent dynamics (**F**(**z**_***t***_)) can be represented as a flow field (Fig. 1c,d, black arrows and light-gray curves), which describes how the instantaneous neural state (**z**_***t***_) evolves from a given location in state-space in the absence of inputs and noise. The action of the input drive (ū_*t*_) corresponds to eliciting a consistent pattern of activity in the neural population, and therefore pushing the trajectory along a direction in state space that can vary both across time and task conditions (Fig. 1c; red and blue arrows; Fig. 1e).

Critically, the simulations show that very different combinations of these two factors can result in very similar trajectories. For example, the three models of decision-making differ in the nature of their inputs and recurrent dynamics, each mimicking a specific behavioral “strategy” for perceptual decision-making^39^, from unstable, impulsive decisions (Fig. 1c, saddle point), to optimal accumulation of evidence (Fig 1c, line attractor), and leaky, forgetful accumulation (Fig. 1c, point attractor). Yet, for the chosen input drive, which depending on the model is either constant (Fig. 1e saddle point) or transient (Fig. 1e, line and point attractor), all three models result in similar single-trial trajectories (Fig. 1c, gray curves) and essentially indistinguishable condition-averaged trajectories (Fig. 1c, blue and red curves).

Analogous observations hold for the models of movement generation (Fig. 1d). Two of the models have no input (Fig. 1e), and are driven entirely by recurrent dynamics starting from condition-dependent initial conditions—one model implements rotational dynamics^12,34^, implying that any variability in the initial condition on a given trial is reflected throughout the entire trajectory (Fig. 1d rotations; gray curves); the other implements what is referred to as a “dynamic attractor”^35^, whereby neural activity is pushed through a narrow channel in state space, and any variability along directions orthogonal to the channel is suppressed (Fig. 1d, dynamic attractor). In the third model, the recurrent dynamics implements a point attractor, and responses are mostly input driven^17^ (Fig. 1d, point attractor). The simulated condition-averages can neither distinguish between the models with or without inputs, nor between the different recurrent dynamics associated with models that lack an input.

Condition-averaged trajectories, which are often used to compare simulated neural responses to measured population activity^11,12,40^, thus appear inadequate to disentangle the functional effects of the recurrent dynamics and inputs, even in these simple models.

### Residual dynamics as a window onto recurrent dynamics

More insights into the computations implemented by the models can be obtained by considering the dynamics of response residuals, the component of single-trial responses that is not explained by the condition-averaged responses^19,41^. Residuals are defined as the difference between a given single-trial trajectory and the corresponding condition-averaged trajectory (Extended Data Fig. 1). We interpret residuals as perturbations away from the condition-averaged trajectory, and then describe how these perturbations evolve over time (Extended Data Fig. 1).

In the simulated models, the dynamics of residuals can be derived analytically (Fig. 2a, Extended Data Fig. 1). First, we define the *effective dynamics*, which describes how the population response would evolve from any given location in state-space and time in the absence of noise. The effective dynamics is obtained by summing the contributions of the recurrent dynamics and the input drive. The *residual dynamics* is then obtained by subtracting, from the effective dynamics, a component corresponding to the instantaneous direction of change along the condition-averaged trajectory (Fig. 2a, see labels over each panel).

The residual dynamics describes how a perturbation away from the condition-averaged neural state would evolve relative to the trajectory over the course of one time-step. In Fig. 2c,d, the blue dot indicates the unperturbed, “reference” neural state, which lies along the average trajectory. The tail of each arrow indicates the residual (the perturbed state), and the arrow-head shows how this residual evolves over one time-step. For the saddle point model (Fig. 2c, saddle point), perturbations along the horizontal direction, away from the trajectory, expand over time (arrows point away from the reference state), whereas perturbations along the vertical direction decay back to the trajectory (arrows point towards the reference state). These dynamics correctly reflect the influence of a saddle point in the vicinity of the examined region of state space (Fig. 1c, box). Likewise, the residual dynamics correctly reveals line attractor and point attractor dynamics in the other two models of decisions (Fig. 2c), as well as the main properties of the recurrent dynamics in the models of movement, i.e. rotational dynamics, decay towards the dynamic attractor, and point attractor dynamics (Fig. 2d). These differences in the underlying recurrent dynamics are less apparent in the effective dynamics, especially in cases where the input drive is strong (Extended Data Fig. 1).

A key property of residual dynamics simplifies the task of estimating it directly from neural responses, even when the underlying computations are non-linear and vary both in time and across state-space location. By definition, the residual dynamics always has a fixed point at the location of the reference neural state (Fig. 2c,d, blue dot; see methods) making it amenable to be estimated using easily interpretable, statistical models characterized by dynamics that is linear and autonomous (i.e. without inputs). Specifically, the residual dynamics can be approximated by a condition and time-dependent, locally linear system, whereby time parameterizes location in state-space along the condition-averaged trajectory (Extended Data Fig. 1). We estimate these linear systems from neural response residuals by combining methods from subspace identification^30,31^ and instrumental variable regression^32^ (Extended Data Fig. 2). These methods, unlike simpler linear regression approaches, can produce robust and unbiased estimates of residual dynamics in biologically realistic settings (Extended Data Fig. 3).

We summarize the residual dynamics through the main properties of the estimated local linear dynamical systems, specifically the magnitude of the eigen-values (EV), the singular values (SV), and the rotation frequency associated with the EV (Fig. 2e-g). Together, these properties distinguish between the various models in Fig. 1c-d. For locations close to the saddle point in the model of decision-making, one of the EV is larger than 1, implying that perturbations along the associated eigen-vector (the horizontal direction in Fig. 1c, left) *expand* over time; the other EV is smaller than one, corresponding to *decay* along the vertical direction (Fig. 1c, left; center of flow field; Fig. 2e, left-most panel; early times). A line attractor results in a single EV of 1 (Fig. 2e, second from left) as horizontal perturbations are *persistent*, i.e. neither expand nor decay, and a point attractor in all EV smaller than 1 (Fig. 2e, third from left; all directions decay). Rotational dynamics results in EV that are complex-valued and thus associated with a non-zero rotation frequency (Fig. 2g). Finally, differences between the magnitude of SV and EV reflect non-normal dynamics, a critical feature of a number of previous models of neural computation^42–44^. The SV larger than 1 in the line attractor model implies that small perturbations along the corresponding right singular vector transiently expand, even though they are persistent (EV=1) or decay (EV<1) over longer time-scales (Fig. 2e,f).

### Residual dynamics reflects local and upstream recurrent dynamics

The simulations illustrate one setting in which residual dynamics, unlike the condition-averaged trajectories, can reveal the key properties of the recurrent dynamics. Specifically, when variability in the inputs is uncorrelated over time, any slow correlations in the residuals are entirely due to (and can be used to infer) the recurrent dynamics (Fig. 1b, top; simple inputs). This constraint, however, is likely to be violated at the level of single areas in biological networks, where any input into a given area could itself be the result of distributed recurrent processing in upstream areas^36,38^. As a result, the input (**u**_*t*_) from Equation 1 can itself be expected to have a component of variability characterized by slow temporal correlations reflecting the upstream recurrent dynamics (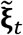 in Fig. 1b, bottom; complex input).

We illustrate the effect of such slow variability on estimates of residual dynamics in simulations of idealized two-area models (Extended Data Figs. 4-5). These simulations show that estimates of residual dynamics reflect not just the “local” recurrent dynamics (**F**_local_, Fig 1b), but rather the combined effects of the recurrent dynamics in the recorded area and in any upstream areas contributing an input to the recorded area^41^ (**F**_upstream_, Fig 1b). For example, residual dynamics with large EV or large rotation frequencies need not imply that the local recurrent dynamics in the recorded area is unstable or rotational, as such dynamics may be implemented also, or exclusively, in areas upstream of the recorded one.

Notably, direct or indirect connections from unrecorded to recorded neurons within the local, recurrently connected population need not result in a functional “input” in the sense of Equation. 1. If neural activity evolves within a low-dimensional manifold, recordings from a large enough subset of neurons within a network can be sufficient to estimate the population state **z**_***t***_ of the entire network^28,29^. The effect of unrecorded neurons in the local network is then fully captured by the recurrent dynamics **F** ^45^ (Fig. 1b, **F** ≈ **F**_local_).

### Neural population responses of decisions and movements in PFC

We compared the dynamics in the models of decision-making and movements generation to neural population dynamics recorded in the pre-frontal cortex (PFC; area 8Ar) of two macaque monkeys performing a saccade-based perceptual decision-making task^46^ (Fig 3a-b). To increase the statistical power of our analyses, we employed a dimensionality reduction technique to “align” the task-related subspaces of neural activity from different experiments with a similar task-configuration (Extended Data Fig. 6; 14-61 experiments per configuration; 150-200 units per experiment). This alignment yielded a shared, 20-dimensional neural state-space explaining >90% of task-related variance in the average neural responses measured across different experiments^29^ (Extended Data Fig. 6). All the analyses below are performed within this aligned subspace, although the main results can be reproduced from sufficiently long single experiments (Extended Data Fig. 7).

**Figure 7.**
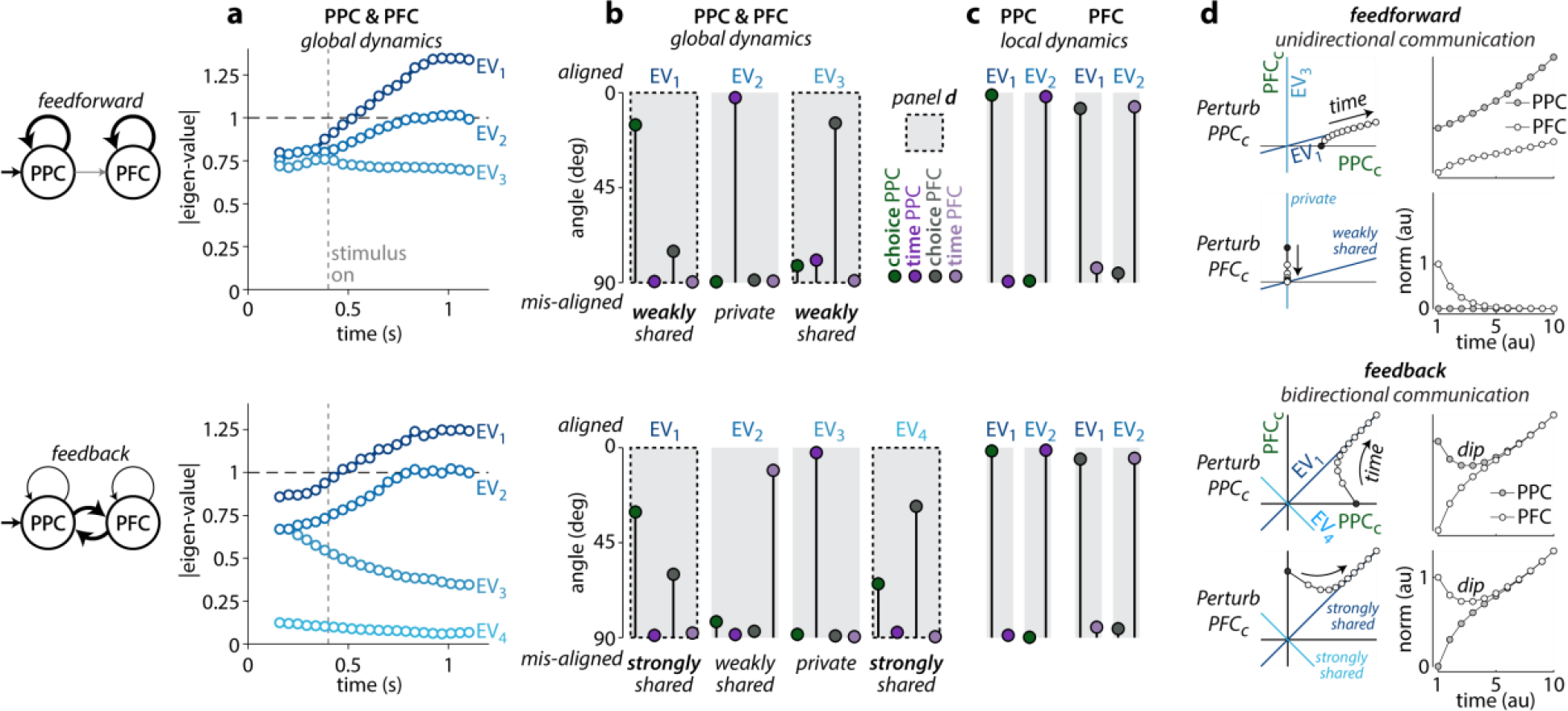
Global residual dynamics resolves local and between-area recurrent contributions. **a**, Time-varying EV magnitudes of the global residual dynamics for the example networks in Fig. 6a (top) and Fig. 6d (bottom). Global residuals are obtained by pooling observations from both areas for a single choice condition (here choice 1). **b**, Alignment (i.e. angle) between the eigenvectors of the global residual dynamics and the choice and time modes in PPC and PFC (see legend), for the feedforward (top) and feedback (bottom) networks. Eigenvectors are estimated 0.7s after stimulus onset (dashed line in **a**). Shared eigenvectors span an angle < 90deg with at least one mode in each area. **c**, Analogous to **b**, but for the eigenvectors of the local residual dynamics (see Fig 6b,e) estimated separately based on PPC or PFC responses. **d**, Effect of local perturbations in two simple models implementing linear dynamics that mimic key features of the estimated global dynamics in **a,b**. Two-dimensional dynamics evolve in a subspace spanned by the PPC and PFC choice modes (PPC_c_ and PFC_c_). Feedforward model: EV_1_ is unstable, the corresponding eigenvector (line) mostly aligns with PPC, but is weakly shared with PFC. EV_3_ is decaying, but unlike in **b** (top), the corresponding eigenvector is entirely private to PFC, to ensure unidirectional communication from PPC to PFC. Feedback model: same eigenvalues as for the feedforward model, but the two associated eigenvectors (EV_1_, unstable, EV_4_ stable; lines) are shared among PPC and PFC. Activity is perturbed along the PPC_c_ and PFC_c_ axes (black circles, left; see labels) and then evolves based on the dynamics determined by the respective EV (white circles, left). The right column shows the norm of activity within each area (i.e. projected onto PPC_c_ or PFC_c_) for the different perturbation types (perturb PPC_c_ or PFC_c_) and models.

The condition-averaged population trajectories in PFC shared important features with the average trajectories of the models in Fig. 1c-d. We visualized the population trajectories through projections onto four distinct, two-dimensional activity subspaces: a “choice” plane, emphasizing choice-related activity; a “time” plane, emphasizing time-varying activity common to both choice conditions; and two “jPC” planes^34^, emphasizing rotational dynamics (Fig. 3c,d; left to right). Only the two jPC planes are orthogonalized with respect to each other, meaning that some planes can capture shared components of the activity (e.g. Fig. 3c, time and jPC_12_ planes). We estimated the planes separately during a decision-epoch, which coincided with the presentation of a random-dots stimulus (Fig. 3c), and during a movement-epoch aligned to the execution of the saccade (Fig. 3d). As in the decision-models (Fig. 1c), PFC responses started in an undifferentiated state prior to stimulus onset (Fig 3c; choice plane; filled dots mark stimulus onset) and gradually diverged based on the upcoming choice of the animal (Fig. 3c, red vs. blue). PFC responses during the movement period showed pronounced rotational components (Fig. 3d, jPC planes; filled dots mark movement onset) partly similar to those in the movement models (Fig. 1d). Prior to saccade-onset, PFC responses fell into largely stationary, choice-dependent states and then transitioned into rotational dynamics following the presentation of the go cue (Fig. 3d, jPC planes).

The measured PFC responses also differed from the model responses in several ways. Consistent with past reports of population dynamics during decisions, working memory and movements, PFC responses reflected strong condition-independent components during both task-epochs (e.g. Fig. 3c,d, time plane)^24,26,40,47^. Such condition-independent components were not implemented in the models shown in Fig 1c-d. Unlike in the models, pronounced choice-related activity occurred along more than one state-space direction (Fig. 3c, choice plane) and rotational dynamics within more than one plane. Moreover, rotational dynamics was observed also during the decision-epoch (Fig. 3c, jPC planes). These shortcomings, however, are common to all models, and do not provide a basis to favor one model over the others as the best explanation for the observed responses.

### Residual dynamics in PFC

To better resolve the contributions of recurrent dynamics to the recorded responses, we characterized residual dynamics in PFC, by proceeding in two steps. First, we estimated a “dynamics subspace”, contained within the previously defined aligned subspace (Fig. 4a, Extended Data Figs. 2,6). The dynamics subspace was defined such that within it, but not outside of it, residuals at any given time are significantly correlated with residuals at previous or future times. Second, we exploited these correlations to estimate residual dynamics within the dynamics subspace, following the same approach as for responses simulated from the models above (Fig. 2e-g, Extended Data Fig. 2,8,9).

**Figure 8.**
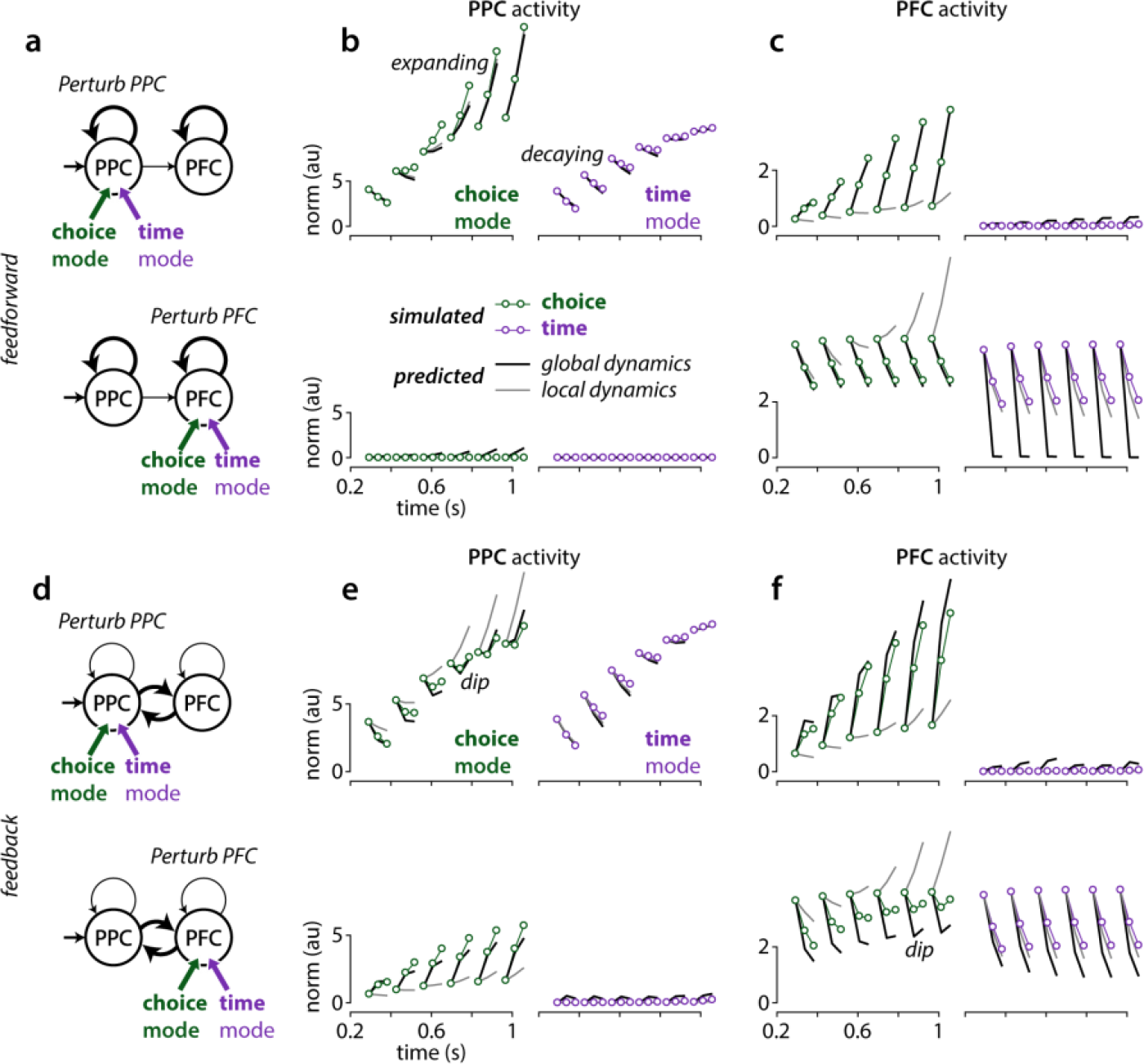
Residual dynamics explains the effects of targeted causal perturbations. Simulated responses to brief perturbations for the two example networks in Fig. 6,7 (small circles) are compared to predictions based on residual dynamics (**a-c** and **d-f**: network without and with feedback between areas). Perturbations are applied locally in each area, along the choice or time mode (green and purple circles) at one of six times in the trial (the first point of each curve in **b-c** and **e-f**). Predictions are based either on the local residual dynamics in the simulated area (gray curves; b,e: PPC; c,f: PFC) or on the global residual dynamics (black curves). **a**, Schematic of the location and type of perturbations shown in **b** and **c** for the network without feedback. **b**, Simulated impulse responses in PPC for perturbations in PPC (top) or PFC (bottom) along the respective choice (left) and time modes (right) compared to the corresponding predictions based on local PPC residual dynamics (gray) or global residual dynamics (black). The norm of the impulse response (y-axis) is shown against time in the trial (x-axis). The last two points on each curve correspond to responses for the two time-steps following the offset of each perturbation. **c**, Analogous to **b**, but for responses in PFC. **d**-**f**, Analogous to **a-c**, but for the network with feedback. Predictions based on the global, but not the local, residual dynamics capture the qualitative features of the simulated impulse responses, i.e. decay, expansion, or decay followed by expansion (e.g. **c**, top-left; **c**, bottom-left; **e**, top-left).

The dynamics subspace was close to 8-dimensional in all configurations (Fig 4a, Extended Data Fig. 8) and was best aligned with directions that explained most task-related variance within the aligned subspace (Fig. 4a, largest dot products at small values along y-axis; Extended Data Figs. 6, 7). Any directions lying outside the dynamics subspace can be thought of as being associated with an EV equal zero, meaning that perturbations along these directions completely decay within a single time step.

The EV magnitudes were strongly time-dependent, but consistently smaller than 1 (Fig. 4b, all EV; Fig. 4e, largest EV; monkey T: p<0.005 for all time points; Monkey V: p < 0.01 for 43 of 44, and p<0.001 for 41 of 44 time points; one-sample, single-tailed t-test on largest EV) implying stable, decaying dynamics. The largest EV were associated with decay time-constants in the range 187-745ms during the decision period (0s to +0.8s following stimulus onset) and 110-913ms during the delay period (−0.5s to +0.3s relative to saccade onset) for monkey T (95% CI, medians = 352ms and 293ms; Fig. 4e, top), and 309-1064ms and 192-3586ms for monkey V (95% CI, medians = 489ms and 491ms; Fig. 4e, bottom). Concurrently with the saccade onset, the largest EV consistently underwent a strong contraction (Fig. 4e; p<3·10^−5^ and p<3·10^−7^ in monkeys T and V; H_0_: largest EV equal at −275ms vs. −5ms relative to saccade onset; two-sample, single-tailed t-test)—the largest measured time constants at saccade onset fell to median values of 159ms in monkey T and 310ms in monkey V, implying that perturbations away from the average trajectory fall back to the trajectory more rapidly during movement.

The residual dynamics had rotational components along some dimensions in both monkeys. In monkey T, the largest rotation frequencies in the residuals (Fig. 4g top; ≈0.5-1 Hz) lay in the approximate range of frequencies for the prevalent rotations in the condition averages (Fig. 3c,d; values for f). In monkey V, even the largest rotation frequencies in the residuals (Fig. 4g bottom, ≈0.25-0.5 Hz) were smaller than those in the condition-averages (0.71-0.84Hz, decision epoch; 1.16-1.34Hz, movement epoch; range across all task configurations). The largest SV of the residual dynamics exceeded the magnitude of the largest EV in both monkeys (Fig. 4e,f; p<0.05 for 43 of 44 and 33 of 44 timepoints in monkeys T and V; two sample– t-test) implying that dynamics was weakly non-normal (Fig. 4h). The largest SV were mostly smaller than 1 in both monkeys (Fig. 4f; p<0.05 for 41 of 44 time points in both monkeys T and V; one-sample, one-tailed t-test). The non-normality is thus not sufficiently pronounced to amplify perturbations, but rather only transiently slows down their decay (Fig. 4c, “transient persistence”).

These empirical findings rule out several models of recurrent dynamics illustrated in Fig 1c-d. In the decision-epoch, the inferred EV are inconsistent with unstable dynamics (EV>1, Figs. 1c,2e; saddle point) and for the most part substantially smaller than what would be expected from persistent dynamics (EV≈1, Figs. 1c,2e; line attractor). In the movement-period, the small EV around the time of the saccade are inconsistent with purely rotational dynamics or a dynamic attractor, which would both result in directions with very slow decay (EV≈1, Figs. 1d,2e; rotations and dynamic attractor). Around the time of saccade onset (−200 to +200ms from onset) the largest inferred EV magnitude (0.80 and 0.88 in monkeys T and V; average) and the largest rotation frequency (0.74 and 0.33 Hz in monkeys T and V; average) imply that perturbations decay by at least 50% within every 1/10^th^ (monkey T) and 1/12^th^ (monkey V) of a rotational cycle. In comparison, during the same time window, the condition-averaged trajectories undergo about 1/4^th^ of a rotational cycle without obvious decay. The quickly decaying residual dynamics, and the mismatch between its properties and the condition-averaged trajectories, are consistent with strongly input driven activity (Figs. 1d, 2e; point attractor).

### Alignment of residual dynamics and condition-averaged trajectories

Additional insights into how recurrent dynamics and inputs contribute to the observed activity can be gained by analyzing the inferred eigenvectors of the residual dynamics. When inputs are weak, the trajectories mostly reflect the properties of the recurrent dynamics, which in turn results in distinct relations between trajectories and residual dynamics.

We illustrate such relations in two models, which we constructed from the line-attractor and rotation models by augmenting the original dynamics (Fig. 1c-d) with two new dimensions. Along these new dimensions, unlike the original ones, the recurrent dynamics was quickly decaying, while the input was strong and condition-independent. We then defined activity subspaces based on the simulated trajectories as in Fig. 3 (Fig. 5a,c) and asked how these subspaces align with the eigenvectors of the residual dynamics. For the augmented line-attractor model, the choice plane is preferentially aligned (angle close to 0) with eigenvectors associated with large EV magnitudes (Fig. 5b top), as slow dynamics along these eigenvectors underlies the observed choice-related activity. Likewise, for the augmented rotations model, the jPC_34_ plane is preferentially aligned with the eigenvectors associated with large rotational frequencies (Fig. 5d top), as these eigenvectors underlie the rotational activity in the jPC_34_ plane. On the other hand, the augmented subspaces are not preferentially aligned with the slow or rotational eigenvectors, as activity within them are mostly input driven. We summarize these relations with a linear regression analysis, whereby negative regression coefficients identify planes where slow or rotational recurrent dynamics may contribute to the observed trajectories (Fig. 5b,d bottom; regression with EV magnitude or rotational frequency, respectively). Notably, the augmented, input driven subspaces in the models are instead aligned with fast or weakly rotational eigenvectors, resulting in positive regression coefficients (Fig. 5b,d bottom). Such positive coefficients are a trivial consequence of the low dimensionality of these models, and thus need not occur in recorded dynamics.

We applied this regression analysis to the PFC responses separately for the decision and movement epochs (Fig. 5e,f). We found significant, negative coefficients primarily in the decision epoch, whereby planes containing choice-related activity were aligned with slow residual dynamics in monkeys T and V (Fig. 5f; choice and jPC_34_ planes; Fig. 5e top) and rotational residual dynamics was aligned with planes containing condition-independent activity in monkey T (Fig. 5f top, time and jPC_12_ planes; Fig. 5e bottom). Coefficients in the movement epoch, on the other hand, were mostly very small or not significant (Fig. 5f). These relations suggest that recurrent dynamics contributes to observed choice-related activity (in both monkeys) and condition-independent activity (in monkey T), but only during the decision-period. Activity at the time of the saccade instead appears more consistent with the influence of a strong input drive^17^, as we also concluded above based on the quickly decaying residual dynamics in this epoch (Fig. 4e, Extended Data Fig. 10).

### Mapping residual dynamics onto multi-area networks

The above analyses illustrate that residual dynamics estimated from single-area recordings can reveal the key functional properties of the recurrent dynamics contributing to neural population activity. However, explicitly mapping these functional properties onto the recorded area is not possible based on single-area recordings (Fig. 1b, right: complex input; Extended Data Fig. 4,5). Below, we use simulations to show how such a mapping could be achieved, either from “global” recordings including all the relevant areas (Fig. 7) or with a combination of local recordings and causal perturbations (Fig. 8).

We consider simulations of a two-area, non-linear, recurrent neural network, which was previously proposed to explain the interplay of posterior parietal cortex (PPC) and PFC during decision-making and working-memory^36^. The network implements both local recurrence within each area (PPC and PFC), as well as long-range connections between the two areas. PPC is assumed to be upstream of PFC, as it alone receives an input with temporally uncorrelated variability (Fig. 1b, simple input) that directly encodes the external stimuli. Here we consider only a limited set among all possible network configurations. First, the strength of local recurrent connectivity is set to be equal in both areas. Second, when feedback connections from PFC to PPC are present, their strength equals those of the feedforward connections from PPC to PFC.

Simulated responses of a random-dots task show choice-dependent and condition-independent components, both in PPC and PFC (Fig. 6a,d; choice and time modes). Both components are very similar between the two networks (Fig. 6a,d), despite substantial differences in their connectivity. The EV of the residual dynamics, estimated *locally* in PPC or PFC, are typically time-dependent (Fig. 6b,e). In particular, the dynamics can change from stable (EV<1) to unstable (EV>1) after the input is turned on, reflecting the non-linear nature of these networks.

To assess the interaction of local recurrence and long-range connections, we focus on residual dynamics estimated along the choice mode in each area (Fig. 6c,f). By design, the choice modes define the “communication subspace” between PPC and PFC in these networks^36,41^—the feedforward and feedback connections between areas are constructed such that activity along the choice mode in one area drives activity along the choice mode in the other area. We summarize the residual dynamics in each network with the peak magnitude of the EV attained within a trial along the choice mode (Fig. 6c,f)

In networks lacking feedback from PFC to PPC, the residual dynamics in PPC naturally only reflects the local recurrence (Fig 1b, simple input), whereby the largest EV gradually increases with stronger local recurrent connectivity. (Fig. 6c, PPC). The residual dynamics in PFC closely resembles that in PPC (Fig. 6c, PFC), but this resemblance conceals a critical difference between the two areas (Fig. 1b, simple vs. complex inputs). In PPC, the residual dynamics reflects the properties of the local recurrent dynamics. The same is not true in PFC, where any EV>1 mostly reflects recurrent dynamics implemented upstream, in PPC. Indeed, if the output of PPC is “shuffled” to remove any temporal correlations (analogous to setting 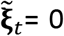 in Fig 1b, complex input), while retaining its time-varying mean, the EV estimated in PFC fall below 1, indicating that local recurrent dynamics (**F**_local_, Fig 1b) in PFC is actually decaying in these networks (Fig. 6b, PFC; dashed). We refer to this effect as an “inflation” of the EV in PFC, due to the correlated input from PPC (Extended Data Figs 4,5).

In networks with long-range feedback, the residual dynamics in PPC and in PFC reflects both the strength of local recurrent connectivity and of long-range connections, whereby reduced local recurrence can be entirely compensated by increased long-range coupling between areas (Fig. 6f). Unlike in the feedforward networks, where the choice results entirely from dynamics unfolding locally within PPC (and simply inherited by PFC), here the choice dynamics reflects a process distributed across both areas.

Overall, these simulations reiterate the conclusion that residual dynamics estimated from recordings in a single area reflect the combined effects of the recurrent computations within the recorded area and the recurrent dynamics unfolding within the output space of any upstream area providing an input to the recorded area (Fig 1b; Extended Data Fig. 4,5). As a consequence, very different combinations of local and long-range connectivity can result in virtually indistinguishable residual dynamics at the level of single areas (Fig. 6a,b vs. d,e).

### Global residual dynamics resolves local and global recurrent computations

If recordings from a single area cannot distinguish between local and long-range recurrent contributions, could recordings from all areas in the network? To address this question, we characterized also the *global* residual dynamics, estimated from the concurrent, pooled responses from PPC and PFC for the two example two-area networks (Fig. 6a,d).

We find that EV magnitudes, even when computed on the global residual dynamics, cannot distinguish between the two networks, with one EV unstable (EV>1), one persistent (EV≈1), and the others decaying (EV<1; Fig. 7a) in both networks. The number of global EV does not robustly distinguish between networks, as it reflects a somewhat arbitrary cutoff in the dimensions to include in the dynamics subspace (excluded dimensions effectively have EV=0).

The *eigenvectors* of the global residual dynamics, however, can distinguish between the two networks. Specifically, we examined the alignment between the global eigenvectors and the local choice and time modes in PPC and PFC (Fig. 7b). Eigenvectors can be qualitatively categorized as being “shared” across areas, or “private” to an area, depending on whether they have substantial projections (i.e. angle<90) onto modes in both areas (shared) or only a single area (private). Both networks result in two eigenvectors that are at least partially shared with the choice modes in the two areas, but the relative projections onto each area varies across networks—the two eigenvectors are only “weakly” shared across areas in the feedforward network, whereas they are more “strongly” shared in the feedback network (Fig. 7b, dashed squares; top vs bottom). Notably, these differences are not reflected in the eigenvectors of the local residual dynamics (Fig. 7c, top vs bottom).

To better understand the functional implications of these differences, we considered the effect of perturbations in two simple models mimicking key properties of the inferred global network dynamics. Both models implement time-independent, two-dimensional, linear dynamics, whereby the two cardinal dimensions (Fig. 7d, left) represent the choice modes in PPC and PFC (Fig. 6a,d). The time modes in each area (Fig. 6a,d) are ignored here. In the “feedforward” model, an unstable eigenvector projects mostly onto the PPC choice mode, while a stable eigenvector is aligned with the PFC choice mode, similarly to EV_1_ and EV_3_ of the feedforward network (Fig. 7b top). In the “feedback” model, instead, both the unstable and stable eigenvectors have large projections onto the PPC and PFC choice modes, like EV_1_ and EV_4_ in the feedback network (Fig. 7b bottom). We simulated a local perturbation either in PPC or PFC by initializing activity along the corresponding choice mode (Fig. 7d left; black points), and then letting activity evolve based on the linear dynamics determined by the respective EV (Fig 7d left; white points).

These simple models exemplify how the arrangement of global eigenvectors determines the directionality of the communication between areas. In the feedforward model, a PPC perturbation causes expanding activity in PPC that propagates to PFC, whereas a PFC perturbation decays in PFC, and does not propagate to PPC (Fig. 7d, top, right column). This unidirectional communication results from non-normal dynamics, as EV_1_ is shared, while EV_3_ is private to PFC (Fig. 7d, top; EV_1_ not orthogonal to EV_3_). In the feedback model, perturbations in either PPC and PFC propagate to the other area (Fig. 7d, bottom, right column). Such bidirectional communication results from normal dynamics, and the fact that both EV_1_ and EV_4_ are shared equally between PPC and PFC. Notably, the estimated residual dynamics does not exactly match all these properties of the simple models. Specifically, EV_3_ is shared, albeit weakly, also for the feedforward network (Fig. 7b top). This structure reflects imperfections in our estimates of residual dynamics, as it erroneously implies the existence of a weak feedback connection from PFC to PPC also in the feedforward network (see also the simulation in Fig. 8).

Interestingly, in the simple models the existence of bidirectional communication is also reflected in the activity of the perturbed area. Somewhat counter-intuitively, activity in the area that was perturbed initially decays, and expands only later; activity in the unperturbed area does not show this dip (Fig. 7d, feedback; PPC and PFC activity in right panels). This dip in activity occurs because any local perturbation is only partially aligned with the shared, unstable direction (EV_1_). Initially, activity in the perturbed area then mostly reflects the rapidly decaying component of activity along the second, global eigenvector (EV_4_).

### Inferring global dynamics with local causal perturbations

We directly verified the insights from these simple linear models by simulating the effect of causal perturbations in the example two-area networks (Fig. 8). We applied *local* perturbations, either in PPC or PFC, by “injecting” an activity pattern corresponding either to the choice mode or the time mode in each area. For each trial, we applied a brief perturbation at one of six different times after stimulus onset, and then let the activity evolve under the influence of the recurrent dynamics and the input. We visualize the effect of a given perturbation as the time-varying norm of the population activity in PPC and PFC for a brief time-window following the onset of the perturbation, averaged over many trials (Fig. 8b-c,e-f; a group of three connected points; analogous to Fig. 7d). The effects of a perturbation depend on the time at which it was applied (Fig. 8b-c,e-f, compare time-courses within each panel), reflecting the time-varying dynamics in these networks (Fig. 7a).

For perturbations applied late in the trial, when dynamics is unstable (Fig. 7a, EV>1), perturbations of the choice modes result in activity that largely matches the dynamics of the simple models above (Fig. 7d). In the feedforward network, PPC perturbations lead to expanding activity in PPC and PFC (Fig. 8b,c; top-left, green), whereas PFC perturbations lead to decaying activity in PFC (Fig. 8c, bottom-left) and no activity in PPC (Fig. 8b, bottom-left). In the feedback network, PPC and PFC perturbations lead to a dip in activity in the perturbed area (Fig. 8e, top-left and Fig. 8f, bottom-left) and to expanding activity in the non-perturbed area (Fig. 8f, top-left and Fig. 8e, bottom-left), as in the corresponding simple model (Fig. 7d, feedback). All these effects are specific to perturbation along the choice modes—perturbations along the time-mode, in either area, result in very different, consistently decaying dynamics (Fig. 8b-c,e-f; purple color).

These varied effects of causal perturbations can be predicted quite accurately based entirely on our estimates of the global residual dynamics (Fig. 7a-b). The predicted time-course of activity following a perturbation at least qualitatively matches the simulated one for most types of perturbations (Fig. 8b-c,e-f, black, R^2^ = 0.97). The predictions do make some qualitative mistakes, specifically for components of the activity that are very small. In particular, because EV_3_ is estimated to be weakly shared across both areas (see Fig. 7b), PFC perturbations are erroneously predicted to somewhat propagate to PPC also in the feedforward network (Fig. 8b, bottom-left). Nonetheless, predictions based on local estimates of residual dynamics fare worse overall (Fig. 8b-c,e-f, black, R^2^ = 0.93). Even the failures of the local predictions can be informative about the properties of the underlying networks (Fig. 8b-c,e-f, gray). For example, the inflation of local PFC residual dynamics in the feedforward network (Fig. 6b) leads to the erroneous prediction that PFC perturbations result in expanding, rather than decaying, PFC activity (Fig. 8c, bottom-left, gray). In the feedback model, predictions based on local residual dynamics instead fail to account for the dip in activity in the perturbed area (Fig. 8e, top-left, Fig. 8f, bottom-left) and underestimate the increase in activity in the unperturbed area (gray; Fig. 8f, top-left, Fig. 8e, bottom-left). Both failures reflect the existence of a global, shared unstable direction, which local residual dynamics cannot adequately capture.

## Discussion

Disentangling the functional contributions of recurrent dynamics and inputs to neural population responses is a key step in understanding neural computations. We demonstrated that disentangling these contributions is challenging, but also that one component of the recorded neural response, the response residuals, is better suited to this challenge. Our analysis of residual dynamics extends previous work that leveraged trial-by-trial variability to understand neural computations^19,20,22,23,41^, by providing a full, quantitative description of the time-varying dynamics of population-level trial-by-trial variability. Our approach can capture dynamics that are globally non-linear^8^, through a series of local approximations capable of resolving fine differences in dynamics across state-space locations and time.

Response residuals are computed by discounting the component of neural responses that is *repeatable* across trials of a given task condition. As a result, residual dynamics can be captured with more easily interpretable models than previous attempts to capture the dynamics of the full, single-trial neural response^5,6^. However, discounting this component does not necessarily amount to removing all the sources of external inputs into the recorded area (Fig 1a), implying that residual dynamics estimated using single-area recordings should not be interpreted as reflecting only the local recurrent dynamics of the recorded area. Instead, residual dynamics reflects the combined effects of local recurrent dynamics and recurrent dynamics unfolding within the output space of any upstream areas that provide an input to the recorded area (Fig 1b, Fig 6). Similar caveats would apply for recordings that include only a subset of areas from the full network.

Resolving the contributions of local (within-area) recurrence and recurrence arising through long-range (across-area) connections is possible with estimates of the global residual dynamics, based on recordings made across the entire network of inter-connected areas (Fig. 7). The resulting description of dynamics in terms of modes (i.e., eigenvectors) that are shared across areas^38^, or private to a single area, appears plausible based on past identification of communication- and null-subspaces between areas^41,48^. However, global residual dynamics, goes beyond a static description based on such subspaces, as it can also capture the dynamics of the responses (Fig. 8) resulting from unidirectional or bidirectional communication between areas (Fig. 7d, top vs. bottom).

Nonetheless, even our local estimates of PFC residual dynamics provide some constraints on the properties of recurrent dynamics implemented by the recorded PFC population, and on the nature of the computations underlying decision-making and movement generation. For one, the largest estimated time constants provide an upper bound on the time-constants of the local recurrent dynamics in PFC (Fig. 4e; 324ms and 510ms in monkeys T and V; medians), as any upstream contribution to PFC responses would have inflated these estimates (Fig. 6b; Extended Data Fig. 4,5). Recurrent dynamics in PFC is thus slow^49,50^, but stable throughout the decision and movement epochs.

This finding does not rule out that the decision-process leading to the monkeys’ choices involves unstable or line-attractor dynamics (Fig. 1c), but those dynamics would have to unfold in areas upstream of PFC^51^, and at least partly outside their communication subspace with PFC. The estimated time-constants would reflect the dynamics of the decision-process if that process unfolded either in PFC alone, or within its communication subspace with other areas (as for all networks in Fig. 6). In such scenarios, our estimates would imply a leaky decision-process, whereby late evidence affects choice more strongly than early evidence. In practice though, monkeys are thought to terminate the accumulation of evidence early in the trial, when a decision-threshold is reached^52^, which would reduce the behavioral effects of any leaks in the accumulation. Notably, a recent study hypothesized that the termination of evidence accumulation coincides with the onset of rotational dynamics in PFC^53^. In our study, condition-independent, rotational dynamics during the decision-epoch also stands out, as in monkey T it is the component of the recorded activity that can be best explained as resulting from recurrent computations (Fig. 5). Irrespective of the possible contributions of PFC to the process underlying the monkeys’ choices, this finding may be indicative of a broader role for PFC in governing transitions between cognitive states^53,54^, e.g. the transition from an uncommitted to a committed state.

Around the time of the saccade, PFC residual dynamics is quickly decaying, largely non-rotational, and only weakly non-normal, implying that PFC does not implement rotational dynamics^12,34^, dynamic attractors^35^, or strongly non-normal^55^ recurrent dynamics of the kind previously proposed to explain movement activity in motor cortex. Rotational dynamics and dynamic attractors are also unlikely to be implemented in an upstream area driving PFC movement responses through a communication subspace, since the signatures of those dynamics would then also appear in PFC residuals (Fig 6, Extended Data Fig. 4). Strong non-normal dynamics in an upstream area, however, could possibly explain the residual dynamics and condition-averages observed in PFC. Non-normal systems can generate large activity transients along directions with only a small projection onto the activity subspace containing the slowest dynamics. If the output from such an upstream area was partially aligned with the activity transients, but orthogonal to the slow dynamics, it could possibly drive strong “input-driven” movement-related activity in PFC without revealing the signatures of the strongly non-normal dynamics that created it.

Alternatively, the mismatch between average trajectories and residuals in the movement epoch could reflect a failure in our estimation procedure. For one, estimates of residual dynamics become biased when trial-by-trial variability is too small, which however does not seem to be the case in our data (Extended Data Fig. 9). For another, dynamics during movement may be strongly non-linear, and thus not well approximated by our local linear description (Extended Data Fig. 1). In both scenarios, our estimated dynamics would not provide a good description of the true dynamics.

A complementary approach to distinguishing between the above interpretations of PFC function, beyond characterizing global residual dynamics, would involve combining local estimates of residual dynamics with targeted causal perturbations^17,24–27^. Residual dynamics naturally leads to predictions of the consequences of such perturbations, and failures of the predictions can be diagnostic of the underlying long-range connectivity (Fig. 8). Most useful in this respect are small perturbations that probe the intrinsic manifold explored by the neural variability^27,28^.

Residual dynamics and the structure of variability may also speak to specific biological constraints at play in neural circuits. The observation of eigenvalues that are smaller, but close to 1 during the decision-epoch is consistent with the underlying neural circuit operating near a critical regime, resulting in large variability and sensitivity to inputs^56–58^. Variability at the level of single neurons is transiently reduced at the time of stimulus and movement onset (Extended Data Fig. 7), potentially reflecting the widespread quenching of variability across cortex in response to task events^20,59^. Near-critical dynamics, non-normality, and variability quenching are thought to emerge naturally in balanced excitation-inhibition (E-I) networks^60,61^. A disruption of E-I balance by the onset of an input could potentially lead to contracting dynamics, and thus reduced variability. Notably, the observed reduction in variability in PFC coincides with contracting dynamics at movement onset, but not at stimulus onset (Extended Data Fig. 7), suggesting that such E-I networks may have to be adapted to fully capture the interactions of internal dynamics, inputs, and variability we observed in PFC.

## Author Contributions

A.R.G and V.M conceived and designed the study. A.R.G developed the methods and performed the analyses, with input from M.S. and V.M. A.R.G and V.M wrote the manuscript. All authors were involved in discussing the results and the manuscript.

## Acknowledgements

We thank John Reppas and William Newsome for the data collection. We thank Kevan Martin and all members of the Mante Lab for their valuable feedback, as well as Nicolas Meirhaeghe, Lea Duncker and Mehrdad Jazayeri for discussions and comments on the manuscript.

## Extended Data Figures

**Extended Data Fig. 1:**
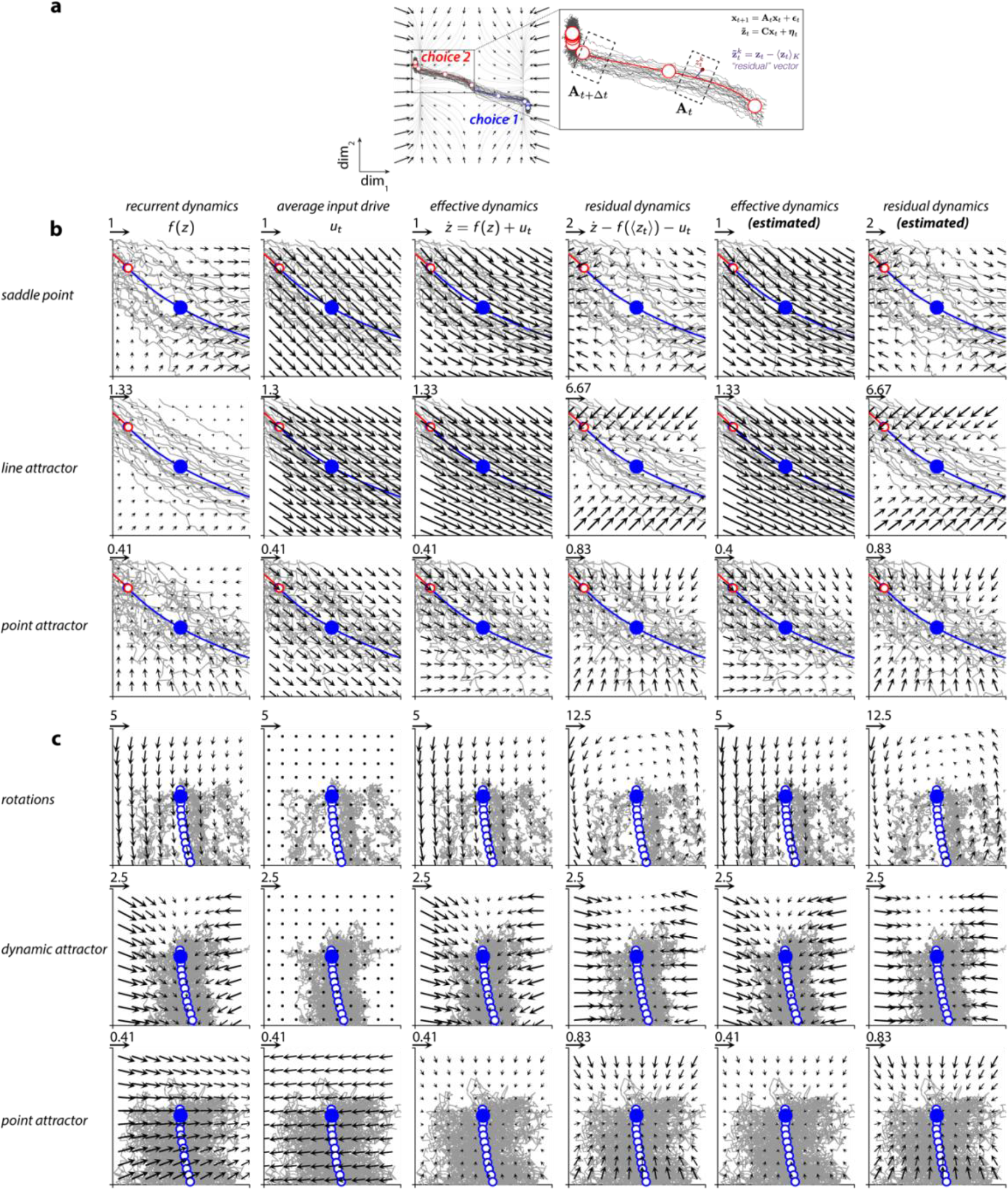
Residual and effective dynamics in models of decisions and movement. **a**, Variability in responses across trials from the same task condition are interpreted as perturbations away from the condition-averaged trajectory. The evolution of these perturbations reflects the properties of the underlying recurrent dynamics (flow field, same conventions as in Fig 1c). Inset on right shows a magnified view of the condition-averaged trajectory (red, choice 2) and corresponding single trials (dark gray) simulated from the saddle point model. Residual vectors at each time (shown in purple for a single trial and time) are computed by subtracting the condition-averaged response at that time from the corresponding single-trial response (purple equation). Time-varying dynamics matrices (**A**_t_) of a linear time-varying, autonomous state-space model (black equations, top-right) are fit to the residuals. These matrices approximate the dynamics in distinct ‘local’ regions of state space (e.g. dashed boxes) and are indexed according to time and condition. **b-c**, Components of the dynamics for the models of decisions (**b**) and movement (**c**) for an example reference time (blue dot) along the condition-averaged trajectory for choice 1. Same conventions as in Fig 2a. Dynamics are shown for a local state-space region close to the corresponding initial condition (boxes in Fig. 1c, d; left). For all models, the estimated effective and residual dynamics (columns 5 and 6) closely match the true effective and residual dynamics (columns 3 and 4). In these models, the residual dynamics (column 4) reflects only the recurrent dynamics (column 1), but is not identical to it. For one, the fixed point of the residual dynamics by definition is located at the location of the reference state (the blue dot), which in general does not match the position of fixed points of the recurrent dynamics (e.g. the red circle in the first row and first column, corresponding to the position of the unstable fixed point in the saddle point model). The position of fixed points of the recurrent dynamics can only be inferred if the external inputs are known, a requirement that is not fulfilled in many experimental settings. For another, consistent drifts resulting from the recurrent dynamics (e.g. the drift along the limit cycle in the dynamic attractor model) are not reflected in the residual dynamics. Such drifts are “subtracted” from the variability in the computation of residuals. Differences in the underlying recurrent dynamics are more apparent in the residual compared to the effective dynamics in cases where the input drive is strong. For example, the average cosine similarity between flow fields is 0.27/0.99 (saddle vs. line-attractor), 0.02/0.94 (saddle vs point-attractor) and 0.58/0.95 (line-attractor vs point-attractor) for the residual/effective dynamics.

**Extended Data Fig. 2:**
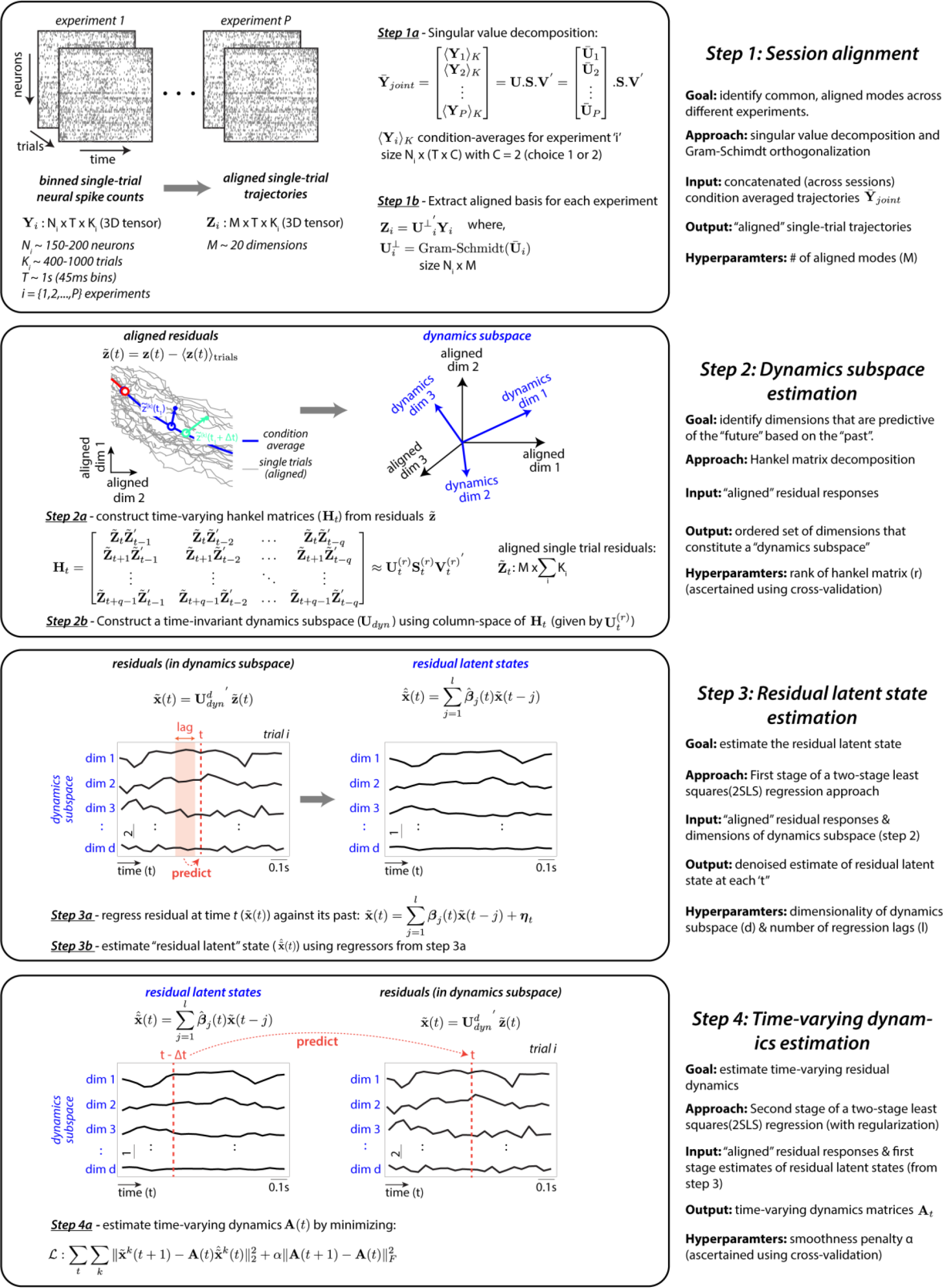
Schematic of analysis pipeline. Schematic depicting the complete data analysis pipeline for inferring residual dynamics from noisy neural population recordings. The pipeline involves four key sequential steps. Step 1: session alignment; involves pooling single trials from different recording sessions in order to increase the statistical power of the analyses Step 2: dynamics subspace estimation; involves using ‘aligned’ single-trial neural residuals to obtain estimates of a dynamics subspace (**U**_dyn_) that effectively contains the residual dynamics; Step 3: residual latent state estimation; involves using the first stage of a two stage least squares (2SLS) approach to estimate a ‘denoised’ latent residual state; and Step 4: dynamics estimation; uses the denoised residual latent states (obtained in step 3) for the second stage of the 2SLS, in order to estimate the time-varying residual dynamics matrices (**A**_t_).

**Extended Data Fig. 3:**
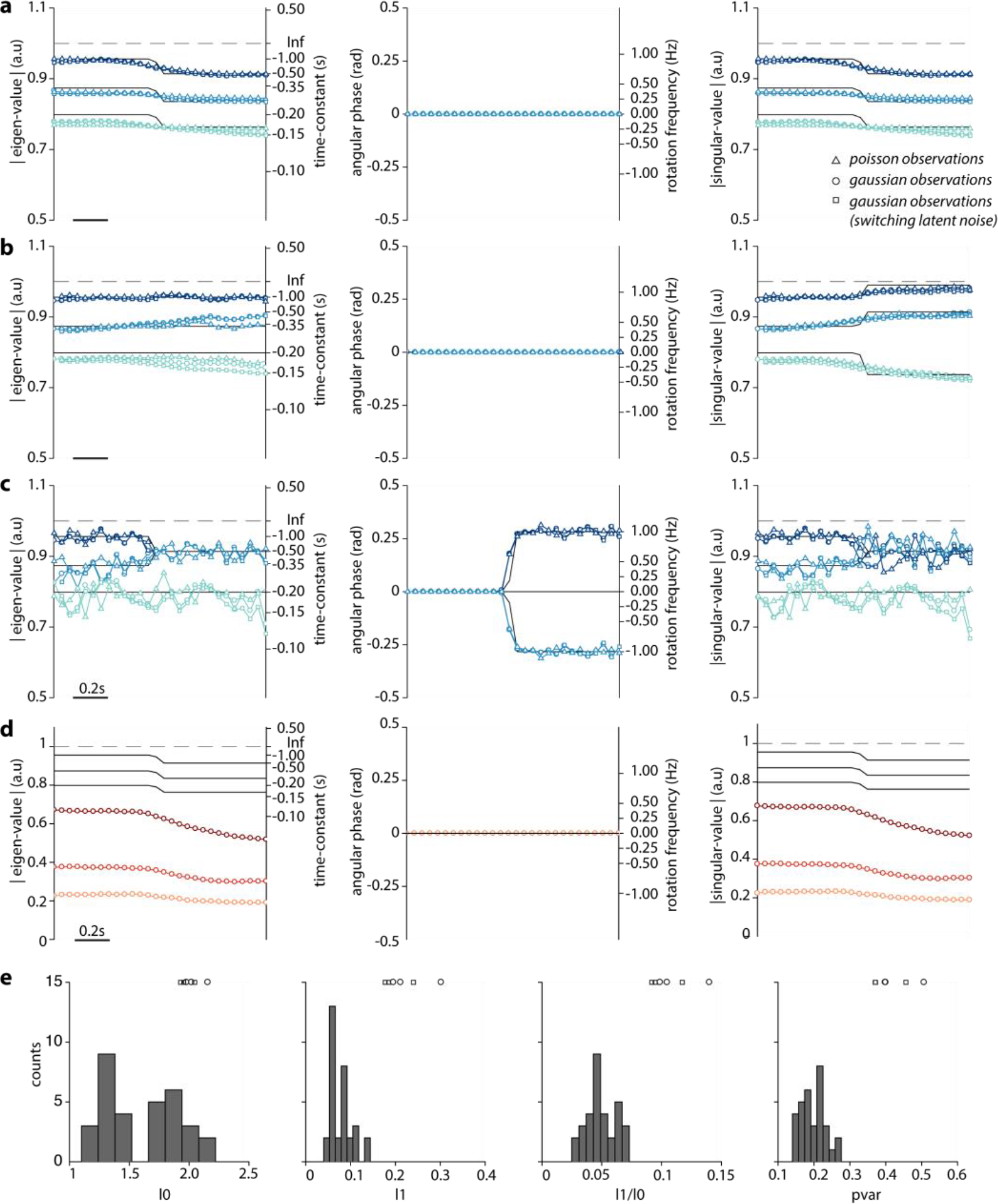
Residual dynamics of simulated, time-varying, linear dynamical systems. **a-c**, Validation of the estimation procedure on simulations of time-varying, linear dynamical systems. Simulations are based on a latent variable dynamical system with 3 latent dimensions and 20 observed dimensions. Residual responses are generated using a gaussian (circle markers: fixed latent noise variance; square markers: latent noise variance switches mid-way through the trial) or poisson (triangle markers) observation process. In all simulations, the properties of the dynamics switch midway through the simulated time window, from slowly decaying to quickly decaying (**a**); from normal to non-normal (**b**); or from non-rotational to rotational (**c**). As in Fig. 4b-d, we characterize dynamics with the magnitude of the eigenvalues (left), the rotational frequency (middle), and the singular values (right). Markers correspond to the estimated residual dynamics, black curves to the ground-truth values. The estimated residual dynamics accurately matches the ground-truth for all types of dynamics and observation models, before and after the switch, and also reveals the time of the switch. We observed this match even when the latent noise variance of gaussian observations was switched at the same time as the eigenvalues/eigenvectors of the dynamics (square markers), demonstrating that estimates of residual dynamics are robust to changes in latent noise variance (see also Extended Data Fig. 5a-b vs e-f). **d**, Analogous to **c**, but for residual dynamics (circles) estimated using ordinary least squares (OLS) instead of two-stage least squares (2SLS) as in **c**. Results are only shown for data simulated using a gaussian observation process. Unlike the 2SLS estimates, the OLS estimates are strongly biased, i.e. the magnitude of the eigenvalues and the singular values are consistently underestimated. These biases are expected—they arise because both the regressors and the dependent variables are corrupted by observation noise (see Methods). The 2SLS instead produces unbiased estimates, as the first stage of 2SLS results in a denoising of the regressors (see also Extended Data Fig. 9). **e**, Parameters of the latent noise and observation noise for the simulations in **a-d** were chosen to approximately match the variability in the measured PFC responses. The variability in the measured responses were quantified in terms of four statistics (l0, l1, l1/l0 and pvar, x-axis; see Methods). Histograms indicate the respective values of these statistics in the neural data (one data point per task configuration, choice condition and monkey; see legend in Extended Data Fig. 6a). The open markers (top, same conventions as **a-c**) indicate the values of the statistics in the simulations for each of the three models.

**Extended Data Fig. 4:**
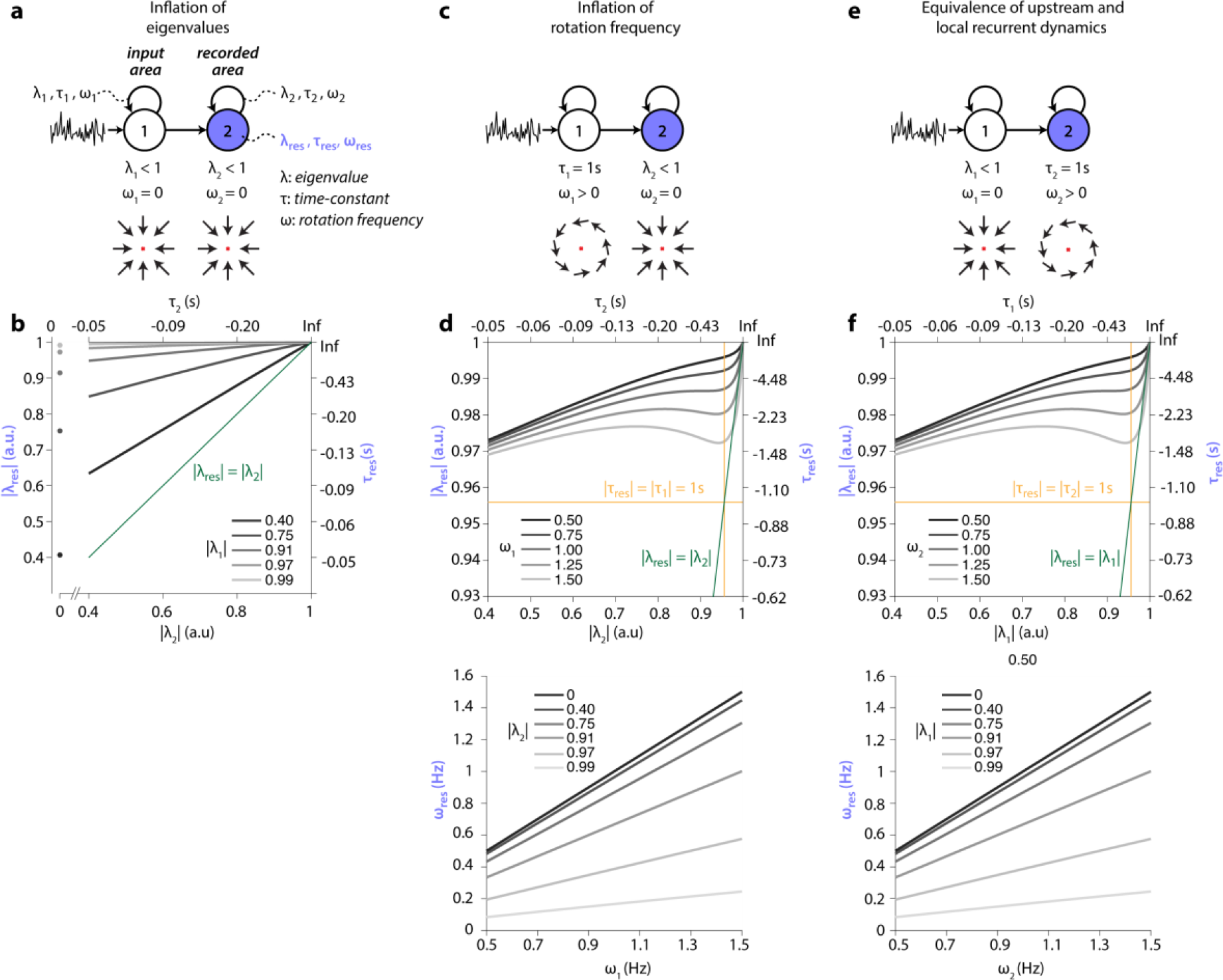
Inflation of local residual dynamics in a linear two-area dynamical system. We systematically explored the effect of correlated input variability on estimates of residual dynamics in a two-area, linear dynamical system. The input area implements 2D isotropic recurrent dynamics characterized by parameters **λ**_**1**_, **τ**_**1**_, and **ω**_**1**_ (eigenvalue, time-constant, rotation frequency). Activity in the input area is externally driven by uncorrelated noise. Values of **λ**_**1**_ closer to 1 result in longer auto-correlation times in the variability of activity in the input area. This activity provides the input into the recorded area, which implements 2d isotropic recurrent dynamics with parameters **λ**_**2**_, **τ**_**2**_, **ω**_**2**_. Residual dynamics at steady-state is estimated from activity of the recorded area. At steady state, estimates can be derived analytically (see Supplementary Math Note B). Because of temporally correlated input variability, the properties of the residual dynamics (**λ**_**res**_, **τ**_**res**_, **ω**_**res**_) in general do not match those of the recurrent dynamics in the recorded area. **a-b**, Inflation of eigenvalues. **a**, Schematic of the model (top) and recurrent dynamics in each area (bottom, flow fields). Recurrent dynamics is stable and non-rotational in both areas. **b**, Residual dynamics (**λ**_**res**_) in the recorded area as a function of recurrent dynamics in the recorded area (**λ**_**2**_, x-axis) and in the input area (**λ**_**1**_, gray lines). The eigenvalues of the residual dynamics are inflated, i.e. **λ**_**res**_ is larger than **λ**_**2**_ (for all gray lines above the green line). Larger **λ**_**1**_ (longer input auto-correlations) lead to stronger inflation. For **λ**_**2**_ = 0 (no recurrent dynamics in the recorded area) **λ**_**res**_ = **λ**_**1**_ (gray circles). **c-d**, Inflation of rotation frequency. **c**, Recurrent dynamics is rotational in the input area, but stable and non-rotational in the recorded area. **d**, Residual dynamics in the recorded area, expressed as the magnitude of the eigenvalue (**λ**_**res**_, top) and the rotation frequency (**ω**_**res**_, bottom). The eigenvalues of the residual dynamics are generally inflated (top), but the relation with **λ**_**2**_ is non-monotonic and depends on **ω**_**1**_. The residual dynamics is rotational (bottom, **ω**_**res**_ > **0**) even though the recurrent dynamics in the recorded area is not (**ω**_**2**_= 0). The inflation of rotation frequency is reduced for increasing **λ**_**2**_. **e-f**, Equivalence of upstream and local recurrent dynamics. **e**, Analogous to **c**, but dynamics is switched between input and recorded area. **f**, Analogous to **d**, but for the dynamics in e. The residual dynamics is identical to that in **d**. In general, residual dynamics in the recorded area reflects the combined effect of local and upstream recurrent dynamics.

**Extended Data Fig. 5:**
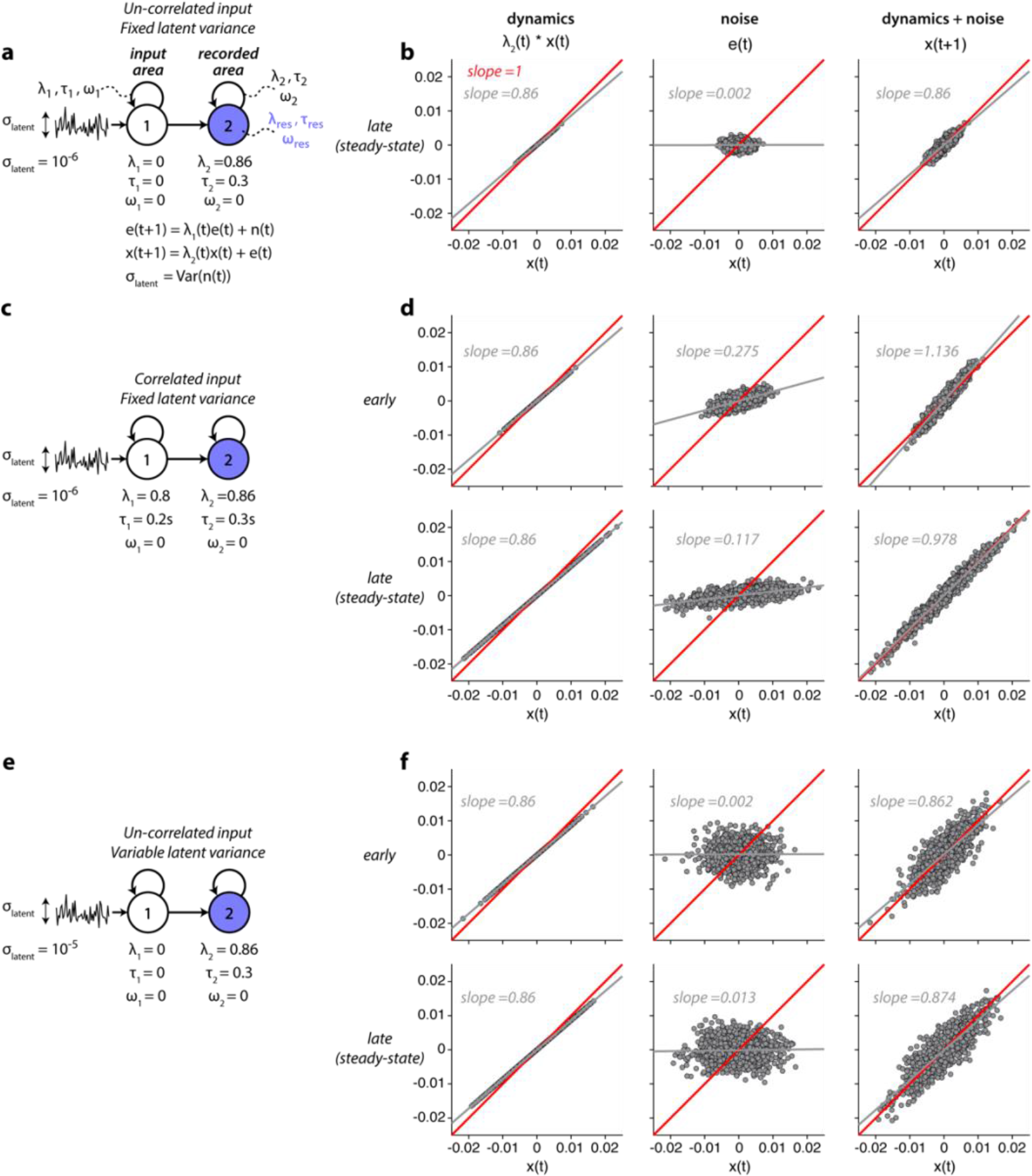
Explanation of input driven inflation in residual dynamics. To gain an intuitive understanding of inflation of eigenvalue magnitude, we consider simulations of two-area linear dynamical systems similar to those in Extended Data Fig. 4a. For simplicity, here we simulate stable 1d-dynamics in each area, whereby variability of the input into the recorded area is either correlated (**c-d**) or uncorrelated (**a-b**, **e-f**), and has fixed (**a-b**, **c-d**) or time-dependent latent noise variance (**e-f**). The variability injected into the input area is always uncorrelated. Recurrent dynamics in the recorded area is identical in all simulations. **a**, Model parameters for the case of uncorrelated input (λ_1_ = 0). **b**, Contributions to activity x in the recorded area at steady-state. Activity x(t) (x-axis) is propagated through the recurrent dynamics (left, y-axis) and added to the noise e(t) (middle, y-axis) to obtain activity x(t+1) at time t+1 (right, y-axis). The noise e(t) corresponds to activity/output of the input area, and is shaped by dynamics determined by λ_1_. Points in the scatter plots correspond to different simulated trials. Estimating the eigenvalue of the residual dynamics in the absence of observation noise amounts to measuring the slope of the regression line relating x(t) to x(t+1) (right, gray line). In this case, this slope is identical to that obtained if the latent noise had not been added to the activity (left, gray line), meaning that residual dynamics correctly reflects the effect of the recurrent dynamics in the recorded area (slope < 0, reflecting λ_2_ < 0; left). **c**, Model parameters for the case of correlated input (λ_1_ > 0 for t > 0; λ_1_ = 0 at other times). **d**, Analogous to **b**, but for the model in **c**. Here activity and noise are shown at two times in the trial: early, when steady-state is not yet reached (top) and late, at steady-state (bottom). At both times, residual dynamics is inflated, i.e. the regression slope between x(t) and x(t+1) (right) is larger than that obtained by applying only the recurrent dynamics (left), indicating inflation of the eigenvalues. Inflation occurs because the noise itself is correlated with activity in the recorded area (middle, slope > 0), an effect that results indirectly from the correlation between e(t) and e(t-1). At steady state, even the inflated residual dynamics is still stable (bottom-right, slope < 1; see also Extended Data 4b). However, immediately after the onset of the correlated input, residual dynamics erroneously reveals an instability (top-right, slope > 1). **e**, Parameters for the case of uncorrelated noise but time-varying noise variance. The variance of the noise injected into the input area is increased at time t = 0, from σ_latent_ = 10^−6^ to 10^−5^. **f**, A change in noise variance does not result in inflation of the residual dynamics, neither early nor late after the change (right, top and bottom; same slope as on the left; see also Extended Data Fig. 3a-c, squares).

**Extended Data Fig. 6:**
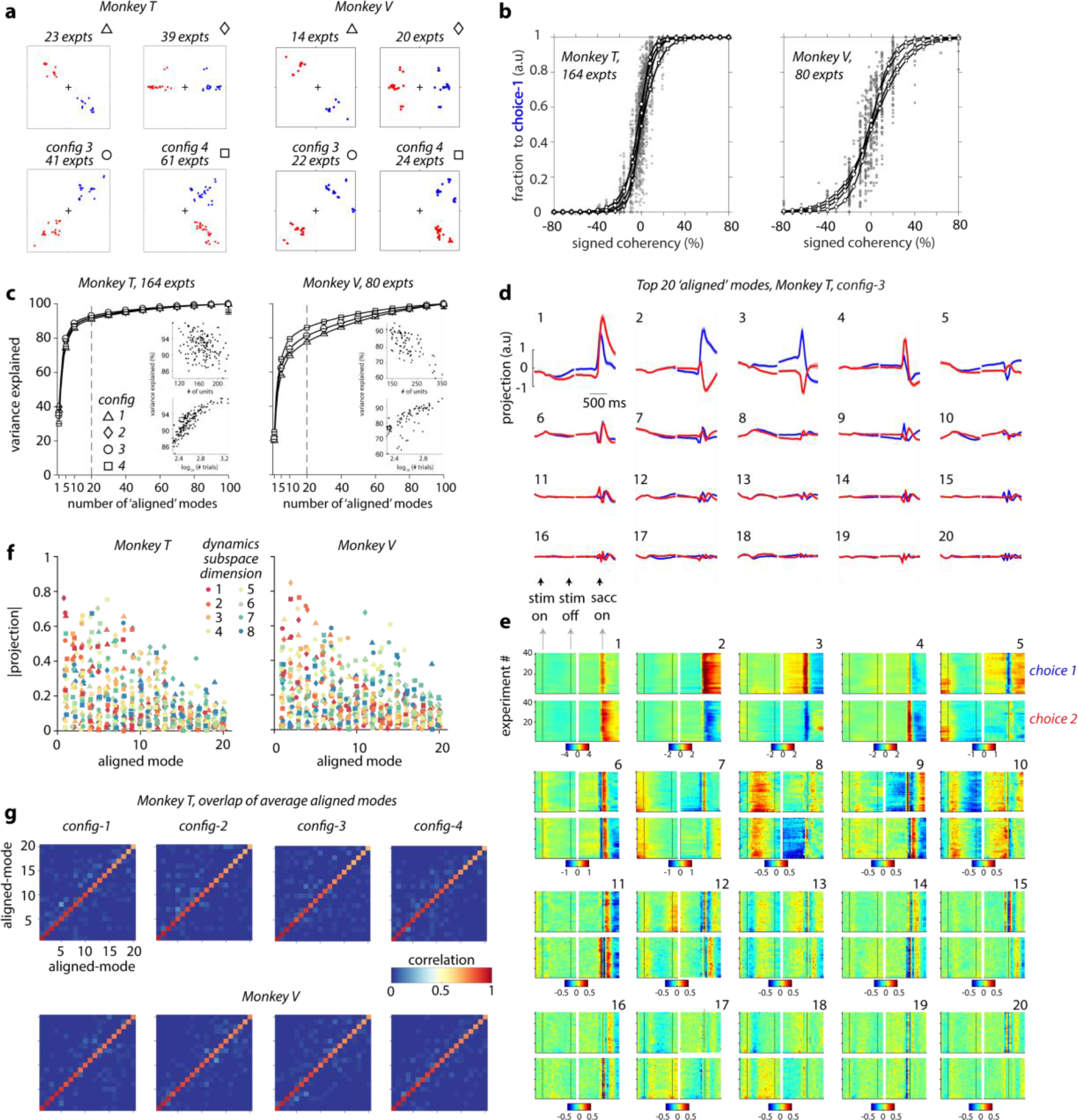
Alignment of neural population responses from different experiments. Validation of our session alignment procedure, Step 1 of the analysis pipeline (Extended Data Fig. 2). We aligned neural population responses of all experiments belonging to the same task configuration and then pooled the aligned single trial responses across experiments before computing the residuals used in estimating the dynamics. The outcome of the session alignment procedure is a set of 20 ‘aligned’ modes for each experiment, defined such that the activity of each mode has the same dependency on time and choice across experiments. **a**, Definition of task configurations. We assigned each experiment to one of four target configurations (distinguished by markers, indicated on top of each panel along with number of experiments) based on the angular position of the targets (blue: choice 1; red: choice 2). The position of the targets was similar, but not identical, across experiments within the same task configuration. (left: Monkey T, right: Monkey V). **b**, Psychometric curves for all experiments in both monkeys (left: Monkey T, right: Monkey V), showing the fraction of saccades to choice-1 as a function of the signed motion coherency. Each gray data point is computed from trials belonging to a single experiment. The employed values of signed coherency varied slightly across experiments, in an attempt to achieve a comparable overall performance in each experiment. Black curves show logistic functions fitted separately to data points from a given task configuration (different markers; see legends in c) and evaluated at logarithmically spaced levels of coherency (positions of the white markers along the x-axis). **c**, Cumulative variance explained in condition-averaged population responses as a function of the number of aligned modes in both monkeys (left: Monkey T, right: Monkey V). We show the mean across experiments of the cumulative variance explained for each task configuration (symbols as in **a**). Error-bars indicating twice the standard error of the mean are mostly smaller than the markers. The cumulative variance explained by the first 20 aligned modes for all 164 experiments in Monkey T and 80 experiments in Monkey V showed a strong positive trend with number of trials (inset, bottom) and a weak negative trend with the number of units (inset, top). **d**, Activity of the first 20 aligned modes (numbered from top-left to bottom-right) for config-3 in monkey T (15,524 trials across 41 experiments) ordered according to the amount of variance explained. Activity is defined as the projection of the population condition averages onto each mode. The projection was computed separately across experiments for choice 1 and choice 2 (blue and red) with responses aligned either to stimulus onset or saccade onset (black arrows). The resulting projections were then averaged across experiments (shading: twice the standard error of the mean across experiments). **e**, Same data as in **d**, but showing the time-course of each aligned mode (numbered from 1 to 20) for each individual experiment (y-axis) separately for the two choice conditions (choice 1 and choice 2, top and bottom sub-panels). Differences in the activation of a given mode across experiments (i.e. across rows in each sub-panel) are much smaller than the differences in the activations across modes (i.e. across sub-panels), demonstrating the success of the alignment procedure. **f**, Absolute value of the projection (y-axis) of the 8 basis vectors (dim-1 through dim-8; red to blue) that span the dynamics subspace (**U**_dyn_, estimated in Step 2 of the analysis pipeline; Extended Data Fig. 2) onto the 20 aligned modes, indicating the relative alignment of the aligned and dynamics subspace. The dynamics subspace is computed separately for each task configuration (symbols as **a**) in each monkey (left: Monkey T, right: Monkey V) and projects most strongly onto the first few aligned components (i.e large projection values for smaller aligned mode number). The dynamics subspace thus largely overlaps with the subspace of activity that capture most of the task-related variance in the responses. **g**, Evaluation of the alignment procedure for all task configurations (columns) in both animals (rows). Each element of the matrix is obtained from the correlation coefficient between the time-courses of two aligned modes (i.e. positions along horizontal and vertical axes). We show the median correlation coefficient across all pairs of dissimilar experiments. Values close to 1 along the diagonal and close to 0 off-diagonal indicate that the time-courses are much more similar across experiments than across modes, indicating successful alignment.

**Extended Data Fig. 7:**
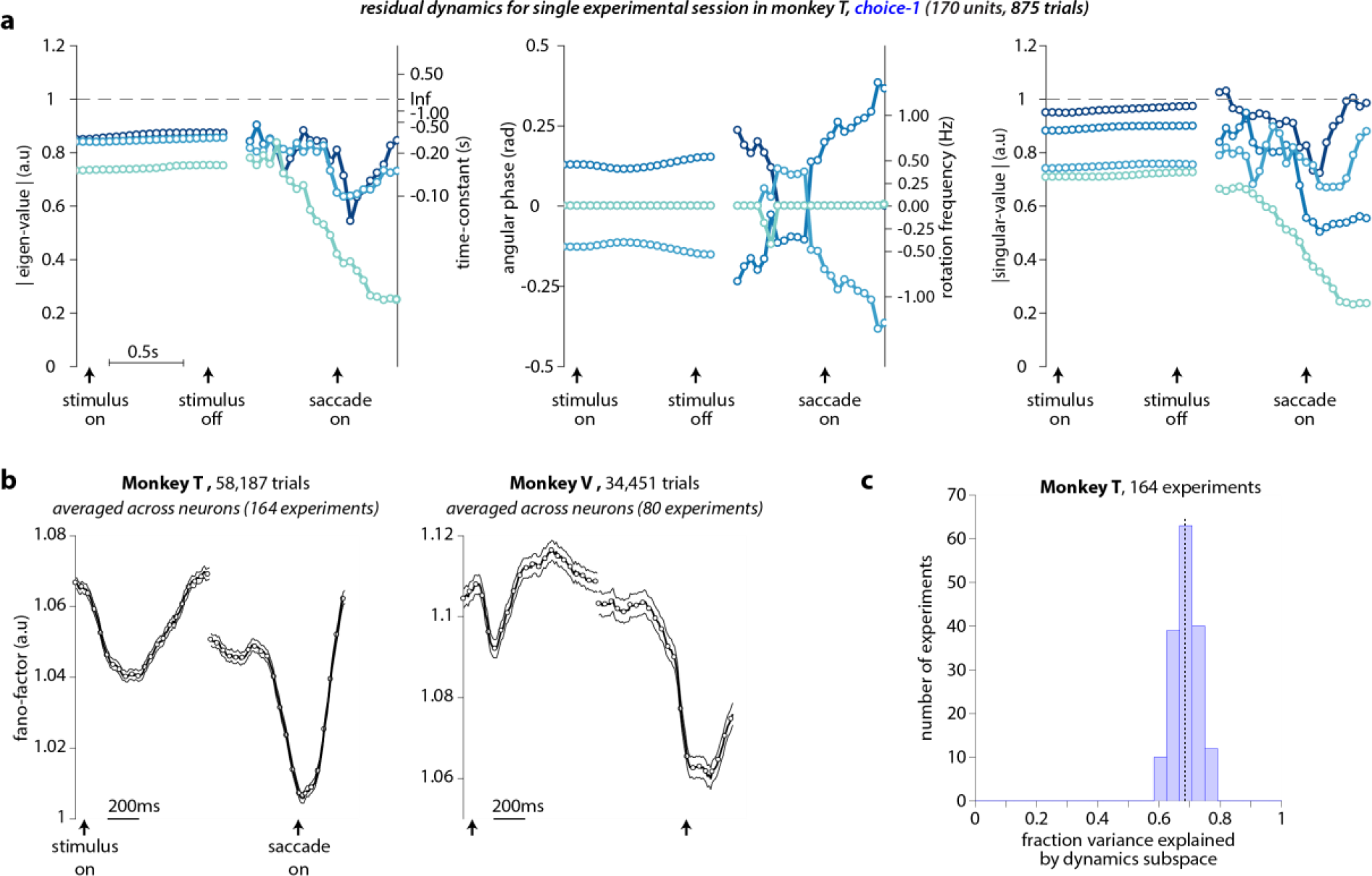
Single session and single unit results. **a**,Residual dynamics estimated using neural data for a single choice condition (choice-1, 875 trials) from a single experiment in monkey T. This experiment has the largest number of trials among all experiments in monkey T. Conventions as in Fig 4b-d. We estimated the residual dynamics directly from high-dimensional residual observations that corresponded to square-root transformed, binned spike-count vectors (dimensionality = number of units; 170 for this session), without performing the session alignment (step 1 in Extended Data Fig. 2). Overall, the properties of the residual dynamics estimated from this single session are similar to those obtained after pooling trials across sessions (Fig 4b-d, 8 dimensional), suggesting that the main features of the residual dynamics (Fig. 4) are not affected by the alignment procedure. The lower dimensionality of the estimated residual dynamics (4 dimensions, blue to cyan; compared to 8 dimensions in Fig. 4a-d) most likely is a consequence of the smaller number of available trials in the single session compared to the aligned sessions. The resulting smaller statistical power makes is harder to estimate, in particular, the faster decaying eigenmodes of the dynamics. **b**, Trial-by-trial variability in single neurons is transiently reduced at the onset of specific task-events. We quantified single neuron variability as the time-varying mean-matched Fano-factor computed by pooling each recorded unit (100ms long time bins), across all experiments in a monkey (empty circles; dashed curve: 95% normal confidence intervals; left: Monkey T, right: Monkey V). In both monkeys, the mean-matched Fano factor undergoes a transient reduction locked to the onset of the stimulus and the onset of the saccade. The reduction in variability around the time of saccade onset coincides with a contraction of the eigenvalues of the residual dynamics (Fig. 4b,e), suggesting that more quickly decaying dynamics may underlie variability quenching at that time. A contraction of eigenvalues, however, does not appear necessary to explain variability quenching, as an analogous contraction is not observed at the time of stimulus onset, despite the consistent reduction in variability at stimulus onset. **c**, Overall fraction of variance explained by the dynamics subspace. We quantified what fraction of the variance of the condition-averaged trajectories in the high-dimensional neural space (state space defined by the individual units) is contained in the dynamics subspace (**U**_dyn_, estimated in Step 2 of the analysis pipeline; Extended Data Fig. 2). Data from all 164 experiments in monkey T. On average in monkey T, the 8-dimensional dynamics subspace explains 68% of the variance (dashed vertical line) in the full trajectories in monkey T. We obtained similar values for monkey V (not shown).

**Extended Data Fig. 8:**
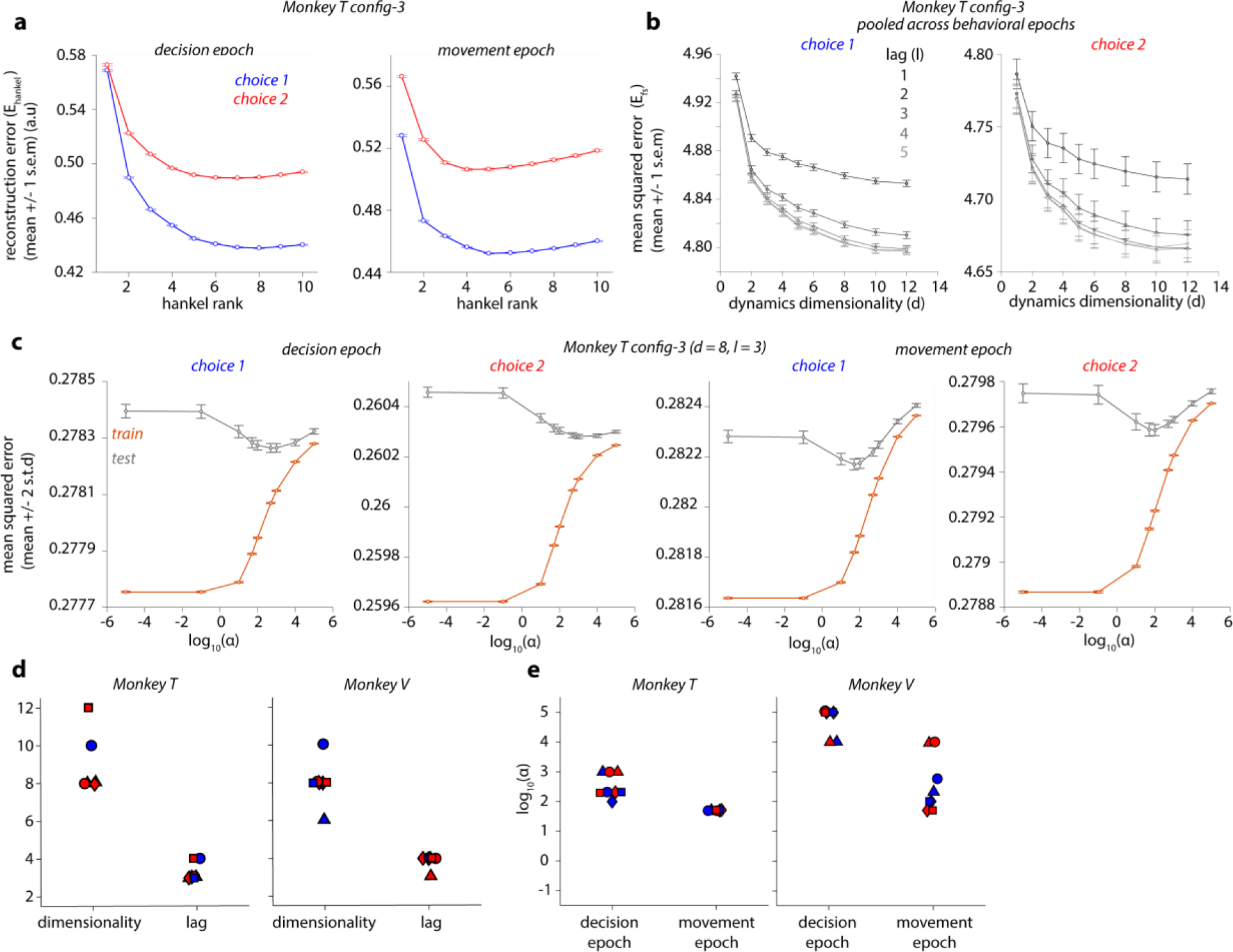
Cross-validation of hyper-parameters used for estimating residual dynamics. **a-c**, Representative results of the cross-validation procedure used to determine the various hyper-parameters of the analysis pipeline (see Extended Data Fig. 2) for neural data from a single task configuration in monkey T (config-3, see Extended Data Fig. 6a). **a**, Cross-validated hankel matrix reconstruction error (E_hankel_; circle: mean over 20 repeats of hold-out cross validation; error bars: 1 s.e.m) plotted as a function of the rank of the hankel matrix (r, step 2 in Extended Data Fig. 2) for residuals from the two epochs (left: decision; right: movement) and two choices (blue: choice 1; red: choice 2). The reconstruction error for each of the 20 repeats was computed by assigning a random 50% of the trials as a “training” set and the rest as a “test” set. **b**, 5-fold cross-validated mean squared error (circles: mean over 5 folds; error bars: 1 s.e.m) of the denoised residual predictions obtained from the first stage of the two-stage least squares regression (2SLS; step 3 in Extended Data Fig. 2), plotted as a function of the hyper-parameters: d (dimensionality of dynamics subspace); and l (number of past lags). For each cross-validation fold, a single mean squared error measure was computed by pooling the denoised predictions across time points in both epochs (left: choice 1; right: choice 2). **c**, 5-fold cross-validated mean squared error (circle: average across 5 ‘repeats’ of the 5-fold cross validation; error bars: 2 standard deviations across repeats) of the residual predictions obtained from the second stage of the 2SLS regression (step 4 in Extended Data Fig. 2), plotted as a function of the smoothness hyper-parameter α for different epochs (left: decision; right: movement) and choice (choice 1 and 2). Both the train (orange) and test (gray) error are shown. **d**, Summary showing the optimal value for the dimensionality d and lag l (step 3 in Extended Data Fig. 2) for all task configurations and monkeys (symbols as in Extended Data Fig. 6a). A dimensionality of 8 and a lag of 3 was deemed optimal for both monkeys and task configurations (used in Fig 4). **e**, Summary showing the optimal smoothness hyper-parameter α (step 4 in Extended Data Fig. 2) for all task configurations and monkeys. Final values of α were chosen to be the same across monkeys in Fig. 4 (decision epoch: α = 200; movement epoch: α = 50) despite a small degree of variability across the two monkeys. Same conventions as in **d**.

**Extended Data Fig. 9:**
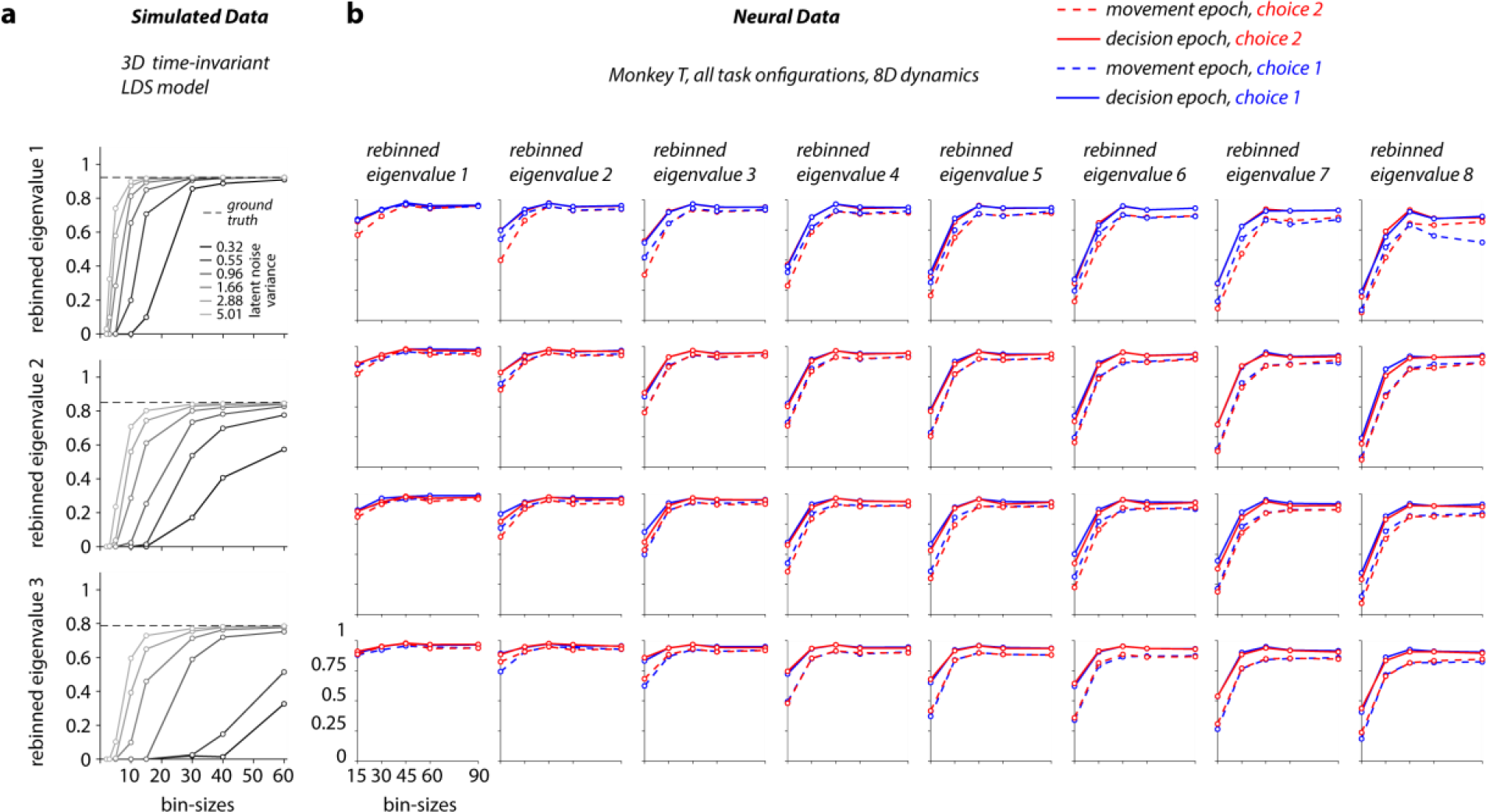
Assessing statistical bias of our eigenvalue estimates. We estimated the residual dynamics for different choices of bin size, to identify the smallest bin size resulting in unbiased estimates. In the discrete time formulation of a linear dynamical system, like the one we use here, re-binning of the responses trivially results in a scaling of the estimated eigenvalues of the residual dynamics. To compensate for this rescaling, here we “mapped” the estimated eigenvalues onto a common, reference bin size (see Methods, Effects of bin size). In the absence of statistical biases, the resulting “re-binned eigenvalue” would be independent of bin size. **a**, Re-binned eigenvalues for simulations of a time-invariant, latent-variable (3 latent dimensions), LDS model (reference bin size = 40ms) as a function of bin-size (dashed line: ground truth). Different gray lines correspond to models with different levels of latent noise (legend). When latent noise is large, estimates of the residual dynamics are biased for small bin sizes, but become unbiased when bin size is sufficiently large (light gray). When latent noise is too small, estimates are biased for any choice of bin size (black). **b**, Estimated, re-binned eigenvalues (reference bin size = 15ms) as a function of bin size for all configurations in monkey T. Columns correspond to the 8 distinct eigenmodes of the estimated 8-dimensional residual dynamics (left to right, largest to smallest EV), rows correspond to task configurations (top to bottom, config-1 to 4; see Extended Data Fig. 6a). Here the re-binned eigenvalues were computed separately for each choice (red vs blue) and averaged in small temporal windows specific to each epoch: 0.2-0.4s relative to stimulus onset (solid lines) and −0.15 to 0.25s relative to saccade onset (dashed lines). All main analyses of recorded neural responses are based on a bin size of 45ms, for which eigenvalue estimates have converged to an asymptote, suggesting that our estimates are not biased. Note that the re-binned eigenvalues for a bin size of 45ms are larger than the corresponding eigenvalues reported in other figures (e.g. Fig. 4b), because the former were mapped onto a reference bin size of 15ms.

**Extended Data Fig. 10:**
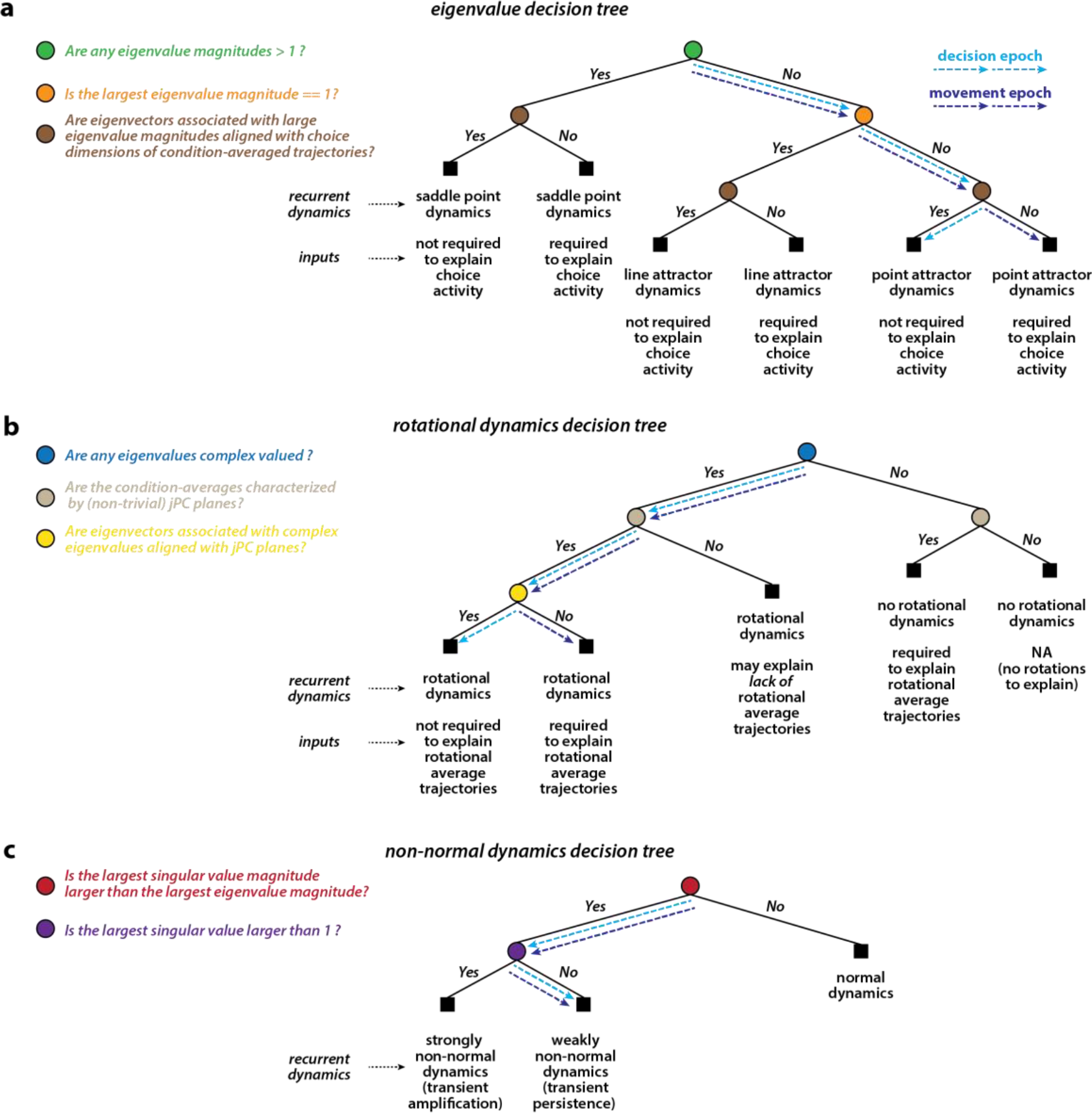
Decision trees for the analysis of residual dynamics. Simplified representation of the decision trees employed in the interpretation of residual dynamics estimated using PFC responses. Each decision tree incorporates questions (circles, legends on left) about the properties of the residual dynamics (see Fig 4) and their relation to the structure of condition-averaged responses (see Fig 5), and leads to specific conclusions about the properties of the underlying recurrent dynamics and inputs (text under squares). These trees primarily present a summary of the main arguments and conclusions about the inferred PFC dynamics. Notably, these trees do not represent the entire combinatorial space of scenarios that could potentially occur in the context of other neural datasets, and therefore should be applied with extreme caution. **a**, Eigenvalue decision tree: inferred properties of recurrent dynamics and inputs based on the magnitude of the eigenvalues of the residual dynamics, and the properties of the condition-averaged responses. Assumes that the condition-averaged responses are modulated by choice. **b**, Rotations decision tree: inferred properties of recurrent dynamics and inputs based on existence of complex eigenvalue and the properties of the condition-averaged trajectories. Assumes that the recurrent dynamics is at most weakly non-normal. **c**, Non-normality decision tree: inferred properties of the recurrent dynamics based on the eigenvalues and singular values.

