## Supplementary Material for "Residual dynamics resolves recurrent contributions to neural computation"

#### SUPPLEMENTARY INFORMATION

##### *Table of Contents*

**I. Methods**
1. Experimental Procedures........ 2
2. Behavioral Task........ 2-3
3. Neural Recordings........ 3
4. Data pre-processing........ 3-4
5. Overview of the analysis approach........ 4-7
6. Neural data analysis pipeline........
   6.1. Session alignment procedure........ 7-9
   6.2. Dynamics subspace estimation........ 9-13
   6.3. Two stage least squares (2SLS) state and dynamics estimation........ 13-16
7. Analysis of residual dynamics........ 17-18
8. Average task-activity subspaces and their relationship to residual dynamics........
   8.1. Computing average task-activity subspaces........ 18-20
   8.2. Comparison of residual eigenvectors to task-activity subspaces........ 20-21
9. Validation of robustness of analysis pipeline........
   9.1. Estimating bias in the estimates of residual dynamics........ 21-23
   9.2. Qualitative estimates of goodness of fit........ 24-26
10. Simulated Models........
   10.1. Models of decisions and movement........ 26-30
   10.2. Linear, time-varying state-space models with uncorrelated input noise........ 30-32
   10.3. Linear, time-varying state-space models with colored input noise........ 32-35
   10.4. Modular two-area recurrent neural network model........ 35-40

**II. Supplementary Math Notes**

A. Background on subspace identification theory……………………………………………… 41-44
B. Analytical derivations of residual dynamics for a linear dynamical system with…. colored input noise 44-47

### I. Methods

#### 1. Experimental Procedures

Neural activity was recorded from neural populations in the pre-arcuate gyrus of two adult male rhesus macaque monkeys: monkey T (14kg) and monkey V (11kg) performing a motion direction discrimination task. All surgical, behavioral, and animal-care procedures complied with National Institutes of Health guidelines and were approved by the Stanford University Institutional Animal Care and Use Committee.

#### 2. Behavioral Task

Monkeys were trained to discriminate the prevalent direction of motion of a random dot kinetogram and reported the perceived motion direction by performing saccadic eye movements to one of two targets on a screen[1] (Fig. 3a). Visual stimuli were presented on a cathode ray tube monitor controlled by a VSG graphics card (Cambridge Graphics, UK), at a frame rate of 120 Hz, and viewed from a distance of 57 cm. Each trial began with the appearance of a small spot that the monkey was required to fixate for a duration of 500ms (fixation period; $\pm$ 1.5° fixation window; degrees of visual angle). Eye position was measured with a scleral search coil (CNC Engineering, Seattle, WA). The fixation period was followed by the appearance of two saccade targets on the screen (target period). The eccentricity (6-18°) and angular location of the targets varied from session to session. After a 400ms delay, the random-dot stimulus was presented within a circular aperture of 7° (monkey T) or 6° (monkey V) in diameter, centered on the fixation point for a fixed duration of 800ms (decision epoch). The difficulty of the task was varied from trial to trial by changing the percentage of dots moving coherently in the same direction (motion strength). The motion strength was chosen randomly on each trial from a fixed set of values. The decision epoch was followed by a delay period of variable duration during which only the fixation point and the two saccade targets were visible to the monkey (300-1100ms, mean = 700ms). The fixation point disappeared at the end of the delay period providing a ‘Go’ cue for the monkey to initiate its choice saccade. The saccade was followed by an additional ‘hold’ period during which the monkey was required to fixate the target before the trial outcome (500-1200ms, mean 900ms; $\pm$ 2-4° fixation window, depending on eccentricity). At the end of the hold time, both targets disappeared and a liquid reward was dispensed for correct trials. For 0% motion coherence trials, the monkeys were rewarded randomly (50% probability).

#### 3. Neural Recordings

Single and multi-unit neural activity was recorded using multi-channel electrode arrays (Blackrock Microsystems, Salt Lake City, UT) with 96 electrodes (length = 1.5 mm; spacing = 0.4 mm). The array was surgically implanted using a previously published protocol[2] in the pre-arcuate gyrus (Brodmann's area 8Ar) a region of prefrontal cortex between the posterior end of the principal sulcus and the anterior bank of the arcuate sulcus (Fig. 3b). The arrays were implanted in the left cerebral hemisphere in both monkeys. Array signals were amplified with respect to a common subdural ground, filtered and digitized prior to spike-sorting. For each of the 96 electrodes, 'spikes' from the entire duration of a recording session were sorted and clustered offline (Plexon Inc., Dallas, Texas) based on a principal component analysis of voltage waveforms. This automated process returned a set of candidate action-potential classifications for each electrode that were subject to additional quality controls, including considerations of waveform shape, waveform reproducibility, inter-spike interval statistics, and the overall firing rate. Spike-sorting yielded approximately 100-200 single and multi-unit clusters distributed across the array in each recording session. Throughout this study, we use the term "units" to collectively refer to both isolated single-units and putative multi-units, as we do not attempt to make a distinction between them. We used neural data from a total of 81 and 76 recording sessions in monkey T and monkey V respectively, with a total of 58,187 and 34,451 trials. A subset of these data were analyzed in previously published studies[3,4].

#### 4. Data pre-processing

We considered neural data in two non-overlapping time epochs of the trial: -200 to +1000ms relative to random-dots onset, or -700 to +500ms relative to movement (saccade) onset. In each recording session, we first removed 'silent' units that had an average firing rate (computed across all trials and time-points) of $<$1Hz. The neural data in most sessions exhibited discontinuities, which manifested as an abrupt synchronous change in the overall firing rate of many units, locked to specific trial indices in the session that constituted 'change-points' in the overall firing rate. Consequently, by detecting these 'change points', we split each recording session into several shorter "experiments" that contained a smaller number of trials, but within which the overall firing rate was stationary. Any experiment with fewer than 200 trials was excluded from further analysis. Furthermore, within each experiment we removed those units that exhibited strong discontinuities in their temporally averaged firing rate across trials. The above procedure yielded a total of 164 experiments in monkey T and 80 experiments in monkey V. For the remainder of the analyses described in the paper, we used square-root transformed binned spike counts[5] computed for each unit in each experiment in non-overlapping time bins (typically of size 45ms, see Extended Data Fig. 9).

We assigned the behavioral and neural data belonging to each experiment to one of four different "task configurations" based on the coarse angular positions of the two choice targets (Extended Data Fig. 6a). To analyze the behavioral performance of the monkeys, each trial was categorized either as choice 1 or choice 2 trial depending on the selected target. In 3 out of 4 task configurations, choice 1 was associated with saccades into the visual hemifield contralateral to the recording site (blue targets, Extended Data Fig. 6a). To quantify the effect of motion coherency on the monkey's behavioral performance, for each experiment we computed the percentage of responses to the choice 1 target as a function of signed motion coherence and, fitted a logistic sigmoidal curve to all the resulting data points that came from the same task configuration (Extended Data Fig. 6b).

#### 5. Overview of the analysis approach

We assume that neural population responses are explained by a latent, low-dimensional state ($\mathbf{z}$), which evolves in time (on each trial) according to Equation 1 (same as main text).

$$\dot{\mathbf{z}}_{\mathrm{t}} = \mathbf{F}(\mathbf{z}_t) + \mathbf{u}_t + \boldsymbol{\epsilon}_t \tag{1}$$

We use a functional interpretation (Fig 1a-b) of the various factors composing Equation 1, with recurrent dynamics (**F**) and inputs ($\mathbf{u}_t$) contributing differently to patterns of variability in the recorded response. The latent noise ($\boldsymbol{\epsilon}_t$) is assumed to capture instantaneous variability (no autocorrelation; e.g: a white gaussian noise process). We explore two different functional regimes for the input – *simple input* (Fig 1b-d,2;Extended Data Fig. 3), in which, $\mathbf{u}_t$ is composed of a deterministic input drive (repeatable across trials of the same condition) that potentially varies across time and task conditions and instantaneous variability (analogous to $\boldsymbol{\epsilon}_t$); and *complex input* (Fig 1b; Extended Data Fig. 4,5) where the variability in the inputs is assumed to contain an additional "slow", temporally correlated component capturing upstream recurrent processing unobserved to the experimenter. Critically, for the analytical derivations below, we assume a simple input regime. The analytical derivations for the complex input regime are considered in **Supplementary Math Note B**.

For a dynamical system specified by Equation 1 and subject to assumptions of simple input (as described above), an analysis of response residuals can reveal the properties of recurrent dynamics $\mathbf{F}(.)$ even when input $\mathbf{u}_t$ is unknown. To illustrate why this is the case, we begin by considering the instantaneous change in the latent state on the $k^{th}$ trial at time 't', which is given by:

$$\dot{\mathbf{z}}_t^k = \mathbf{F}\left(\mathbf{z}_t^k\right) + \mathbf{u}_t + \boldsymbol{\epsilon}_t \tag{2}$$

Likewise, we assume that the instantaneous change in the average state across trials (denoted by $\langle\, . \rangle$) can be written as:

$$\langle \dot{\mathbf{z}_t} \rangle = \mathbf{F}(\langle \mathbf{z}_t \rangle) + \mathbf{u}_t \tag{3}$$

The equality in the above equation follows from Equation 2, in particular, if $\mathbf{F}$ is locally linear. Furthermore, the assumption that $\boldsymbol{\epsilon}_t$ is a zero-mean instantaneous noise process, eliminates it from Equation 3. Our approach is inherently local, as we seek to estimate the dynamics separately at each time and/or state-space location along a given condition-averaged neural trajectory (Extended Data Fig. 1a). The discrete-time state update equations for the average state and the $k^{\text{th}}$ single trial state can then be approximated as:

$$\langle \mathbf{z}_{t+1} \rangle \quad = \langle \mathbf{z}_t \rangle + \Delta t \cdot (\mathbf{F}(\langle \mathbf{z}_t \rangle) + \mathbf{u}_t) \tag{4}$$

$$\mathbf{z}_{t+1}^k \;=\; \mathbf{z}_t^k + \Delta t \cdot \left(\mathbf{F}\left(\mathbf{z}_t^k\right) + \mathbf{u}_t + \;\boldsymbol{\epsilon}_t\right) \tag{5}$$

The dynamics of the residual vector ($\tilde{\mathbf{z}}$) on the $k^{\text{th}}$ trial is then obtained by subtracting Equation 4 from Equation 5:

$$\underbrace{\mathbf{z}_{t+1}^k - \langle \mathbf{z}_{t+1} \rangle}_{=\tilde{\mathbf{z}}_{\mathrm{t+1}}^{\mathrm{k}}} \quad = \underbrace{\mathbf{z}_t^k - \langle \mathbf{z}_t \rangle}_{=\tilde{\mathbf{z}}_{\mathrm{t}}^{\mathrm{k}}} + \Delta t \cdot \left(\mathbf{F}\left(\mathbf{z}_t^k\right) - \mathbf{F}(\langle \mathbf{z}_t \rangle) + \;\boldsymbol{\epsilon}_t\right) \tag{6}$$

Thus, by rearranging the above equation, we see that the temporal evolution of the residuals is itself governed by a differential equation which can be expressed in terms of the single trial dynamics as follows:

$$\underbrace{\dot{\tilde{\mathbf{z}}}_t}_{\text{residual flow}} \quad = \underbrace{\left(\mathbf{F}\left(\mathbf{z}_t^k\right) + \mathbf{u}_t\right)}_{\text{"effective" flow at } \mathbf{z}_t^k} - \underbrace{\left(\mathbf{F}(\langle \mathbf{z}_t \rangle) + \mathbf{u}_t\right)}_{\text{"effective" flow at } \langle \mathbf{z_t} \rangle} + \;\boldsymbol{\epsilon}_t \tag{7}$$

Further, the second term on the right-hand side of Equation 6 can be re-expressed in terms of the residuals.

$$\mathbf{F}\left(\mathbf{z}_t^k\right) - \mathbf{F}(\langle \mathbf{z}_t \rangle) + \;\boldsymbol{\epsilon}_t \quad = \mathbf{F}\left(\langle \mathbf{z}_t \rangle + \tilde{\mathbf{z}}_t^k\right) - \mathbf{F}(\langle \mathbf{z}_t \rangle) + \;\boldsymbol{\epsilon}_t \tag{8}$$

Subjecting the first term on the right-hand side of Equation 8 to a first order Taylor expansion, we obtain:

$$\mathbf{F}\left(\langle \mathbf{z}_t \rangle + \tilde{\mathbf{z}}_t^k\right) = \mathbf{F}(\langle \mathbf{z}_t \rangle) + \underbrace{\nabla \mathbf{F}|_{\langle \mathbf{z}_t \rangle}}_{=\mathbf{J}_t} \, . \; \tilde{\mathbf{z}}_t^k + \text{higher order terms} \tag{9}$$

Ignoring the higher order terms and re-expressing Equation 6 using Equations 8 and 9 yields a discrete-time, time-varying, linear dynamical system at the level of the residuals:

$$\begin{aligned} \tilde{\mathbf{z}}_{t+1} &= \tilde{\mathbf{z}}_t + \Delta t . (\mathbf{J}_t \tilde{\mathbf{z}}_t + \boldsymbol{\epsilon}_t) \\ &= \Big(\underbrace{\mathbf{I} + \Delta t . \mathbf{J}_t}_{=\mathbf{A}_t}\Big) \tilde{\mathbf{z}}_t + \Delta t . \boldsymbol{\epsilon}_t \end{aligned} \tag{10}$$

The time-varying "dynamics matrix" ($\mathbf{A}_t$) that maps residuals from time 't' to 't+1' is directly related to the Jacobian ($\mathbf{J}_t$) of the dynamical system instantiated by the recurrently connected neural network, computed at each state along the average trajectory. Critically, $\mathbf{u}_t$ does not appear in Equation 10, meaning that, subject to the above assumptions on the nature of $\mathbf{u}_t$ (simple input; Fig 1b: top row) and $\boldsymbol{\epsilon}_t$, the residual dynamics $\mathbf{A}_t$ only reflects the recurrent dynamics. Ultimately, our goal is to reliably estimate $\mathbf{A}_t$ using neural population spike count residuals, and use them to make inferences about the types of neural computations specified by the dynamics implemented in recurrent neural circuits.

In practice, we assume that neural population spike count residuals can be modeled using a latent-variable, autonomous, linear, time-varying dynamical system as described below

$$\begin{aligned} \mathbf{x}_{t+1} &= \mathbf{A}_t \mathbf{x}_t + \boldsymbol{\epsilon}_t \\ \tilde{\mathbf{z}}_t &= \mathbf{C} \mathbf{x}_t + \boldsymbol{\eta}_t \end{aligned} \tag{11}$$

where, $\mathbf{x}_t$ is a low-dimensional 'latent' residual state whose temporal evolution is parameterized by the dynamics we seek to estimate ($\mathbf{A}_t$). The observations ($\tilde{\mathbf{z}}_t$) result from a linear combination (through an "observation matrix" $\mathbf{C}$) of the latent residual states, and $\boldsymbol{\epsilon}_t$ and $\boldsymbol{\eta}_t$ are "latent" and "observation" noise terms respectively, which are modelled as white gaussian noise disturbances. In this study, we refer to the subspace spanned by the columns of $\mathbf{C}$ as the "dynamics subspace" (later denoted by the matrix estimate $\mathbf{U}_{dyn}$). The "dynamics subspace" contains the *dynamically relevant* portion of response variability, meaning that trial-by-trial variability along any dimension within it covaries with variability along the same or other dimensions at later times. The observations ($\tilde{\mathbf{z}}_t$) could, in principle, directly correspond to the neural spike count residual vector. However, in our analysis pipeline, they correspond to a lower dimensional projection of the high-dimensional neural spike count residual vector, obtained after aligning neural data across multiple experiments.

A linear time-varying latent dynamical process as in Equation 11 makes exact probabilistic inference of the relevant parameters an intractable problem. Instead of using probabilistic techniques that rely on approximate inference[6,7], we estimated parameters $\mathbf{C}$ and $\mathbf{A}_t$ using an alternate set of approaches that combined subspace system identification (SSID) theory[8–10] and instrumental variable (IV) regression[11–13]. We provide a brief overview of the concepts underlying SSID of linear dynamical systems in **Supplementary Math Note A**. Going from neural spike counts to estimates of the dynamics subspace ($\mathbf{C}$) and the time-varying residual dynamics matrices ($\mathbf{A}_t$) involves a data analysis pipeline that can be broken down into four distinct steps (Extended Data Fig. 2), which are described in the following section.

### 6. Neural data analysis pipeline

The neural data analysis pipeline to estimate residual dynamics consists of four steps (Extended Data Fig. 2), that briefly involve (i) aligning neural responses across different experiments (**session alignment**), (ii) using 'aligned' residuals pooled across multiple experiments to estimate the dynamics subspace i.e. the column space of the observation matrix $\mathbf{C}$ in Equation 11 (**dynamics subspace estimation**), (iii) using 'aligned' residuals and the dynamics subspace to estimate the latent residual state $\mathbf{x}_{\mathrm{t}}$(**residual latent state estimation**) and, (iv) combining the outputs of the previous three steps to estimate the time-varying residual dynamics $\mathbf{A}_t$ (**time-varying dynamics estimation**). We used methods derived from SSID theory of linear systems in Step (ii) and two-stage least squares (2SLS) approach based on the concept of instrumental variables for Steps (iii) & (iv). We describe each of these four steps in further detail below.

#### 6.1. Session alignment procedure

To improve the statistical power of our analyses, we sought to pool trials across different experiments. To do so, we performed an alignment of condition-averaged, task-relevant subspaces of neural activity, associated with different experiments (Extended Data Fig. 2, Step 1). The goal of this alignment procedure was to identify a low-dimensional subspace capturing task-related variance that is common to all experiments. Such an alignment procedure was based on a key assumption that the neural population activity recorded across different experiments can be interpreted as different high-dimensional readouts of a fixed set of low-dimensional neural response patterns[5,14], an assumption that was valid given the stereotypy of the behavior and overall stability of the neural recordings. A full example of the results of the alignment procedure applied to neural data from a single monkey and a single task configuration is shown in Extended Data Fig. 6.

The alignment procedure operated on a block condition average matrix ($\overline{\mathbf{Y}}_{\mathrm{joint}}$) that was constructed (separately for each task configuration) by concatenating row-wise, the condition averaged, square-root transformed, neural population binned spike-counts ($\overline{\mathbf{Y}}_{\mathrm{i}}$) for each experiment:

$$\overline{\mathbf{Y}}_{\mathbf{joint}} = \begin{pmatrix} \overline{\mathbf{Y}}_1 \\ \overline{\mathbf{Y}}_2 \\ \vdots \\ \overline{\mathbf{Y}}_{\mathrm{P}} \end{pmatrix} = \mathbf{U}.\mathbf{S}.\mathbf{V}' \qquad (12)$$

where, $\overline{\mathbf{Y}}_{\mathrm{i}}$ is a $\mathrm{N_i} \times (\mathrm{T_{all}} \times \mathrm{C})$ data matrix (mean centered such that each row has zero mean; we denote the subtracted row means by $\boldsymbol{\mu}_i$ for experiment 'i'). Here, $\mathrm{N_i}$ is the number of units recorded in experiment 'i', $\mathrm{T_{all}}$ are the total number of time bins in both the decision and movement epochs, C is the total number of conditions, and P is the total number of experiments to be aligned. $\overline{\mathbf{Y}}_{\mathrm{i}}$ was computed by sorting single-trial trajectories (composed of binned spike counts) based on the saccadic choice of the monkey and averaging the single trial trajectories across all trials assigned to a particular choice (C = 2, choice-1 or choice-2).

Through the singular value decomposition (SVD) of $\overline{\mathbf{Y}}_{\mathrm{joint}}$ we obtained a matrix of left singular vectors ($\mathbf{U}$ in Equation 12) that was orthonormal and of dimensionality $\sum_{\mathrm{i}=1}^{\mathrm{P}} \mathrm{N_i} \times (\mathrm{T_{all}} \times \mathrm{C})$. The 'aligned' coordinate basis specific to an experiment, was read-out as block rows of the matrix $\mathbf{U}$ as illustrated below:

$$\mathbf{U} = \begin{bmatrix} \mathbf{u}_1^1 & \mathbf{u}_1^2 & \dots & \mathbf{u}_1^{\mathrm{T.C}} \\ \mathbf{u}_2^1 & \mathbf{u}_2^2 & \dots & \mathbf{u}_2^{\mathrm{T.C}} \\ \vdots & & \ddots & \vdots \\ \mathbf{u}_{\mathrm{P}}^1 & \mathbf{u}_{\mathrm{P}}^2 & \dots & \mathbf{u}_{\mathrm{P}}^{\mathrm{T.C}} \end{bmatrix} \qquad (13)$$

where, $\mathbf{u}_{\mathrm{i}}^{\mathrm{j}}$, a vector of dimensionality $\mathrm{N_i} \times 1$ is the left singular 'sub-vector' corresponding to $j^{\text{th}}$ mode in experiment *'i'*. We now define the matrix $\overline{\mathbf{U}}_{\mathrm{i}}$ as the $i^{\text{th}}$ block row of $\mathbf{U}$ and proceed to orthogonalize the columns of $\overline{\mathbf{U}}_{\mathbf{i}}$ using the QR decomposition as follows:

$$\overline{\mathbf{U}}_{\mathrm{i}} = \mathbf{Q}_{\mathrm{i}}.\mathbf{R}_{\mathrm{i}} \qquad (14)$$

We define the matrix $\mathbf{U}_{\mathrm{i,M}}^{\perp}$ (of dimensionality $\mathrm{N_i} \times \mathrm{M}$) as the first M columns of the orthogonal matrix $\mathbf{Q}_{\mathrm{i}}$ . The columns of $\mathbf{U}_{\mathrm{i,M}}^{\perp}$ constitute a set of orthogonal basis vectors that span a M-dimensional 'aligned' coordinate space for experiment 'i'. The 'aligned' single trial response $\mathbf{z}_t^i(k)$ for time-bin 't' and trial 'k' in the $i^{\text{th}}$ experiment (a vector of dimensionality $\mathrm{M} \times 1$), is then given by:

$$\mathbf{z}_t^i(k) = \mathbf{U}_{\mathrm{i,M}}^{\perp}{}'.\left(\mathbf{y}_t^i(k) - \boldsymbol{\mu}_i\right) \qquad (15)$$

where, $\mathbf{y}_t^i(k)$ is the corresponding original square-rooted binned spike count population vector (of dimensionality $\mathrm{N_i} \times 1$). Therefore, collecting all the $\mathbf{z}_t^i(k)$ across time points and trials for each experiment ‘$i$’ resulted in P aligned single trial data matrices $\mathbf{Z}^{\mathrm{i}}$ (i = 1, 2, …, P), each of size $\mathrm{M} \times \mathrm{T}_{\mathrm{all}} \times \mathrm{K}_{\mathrm{i}}$ , where $\mathrm{K}_{\mathrm{i}}$ is the number of trials in the $i^{\text{th}}$ experiment.

To choose M (the number of aligned modes), we inspected the cumulative amount of variance explained in the condition-averaged data matrix $\overline{\mathbf{Y}}_{\mathbf{i}}$ as a function of M, by progressively retaining a larger number of columns in the construction of $\mathbf{U}^{\perp}_{\mathrm{i,M}}$. The variance explained by M aligned modes in the $i^{\text{th}}$ experiment was then computed as - 100* $\left(1 - \frac{\mathrm{var}\left(\overline{\mathrm{Y}}_{\mathrm{i}} - \widetilde{\widetilde{\overline{\mathrm{Y}}}}_{\mathrm{i}}\right)}{\mathrm{var}(\overline{Y}_{\mathrm{i}})}\right)$, where $\widetilde{\widetilde{\overline{\boldsymbol{Y}}}}_{\mathrm{i}}$ is a denoised set of condition-averages obtained through the following projection:

$$\widetilde{\widetilde{\overline{\mathbf{Y}}}}_{\mathrm{i}} = \mathbf{U}^{\perp}_{\mathrm{i,M}} . \mathbf{U}^{\perp\,\prime}_{\mathrm{i,M}} . \overline{\mathbf{Y}}_{\mathrm{i}} \tag{16}$$

For our subsequent analyses, we chose M = 20, based on the cumulative variance explained plots (Extended Data Fig. 6c). To visualize each of the individual, aligned activity modes, we projected $\overline{\mathbf{Y}}_{\mathbf{i}}$ into the space spanned by the first 20 columns of $\mathbf{U}^{\perp}_{\mathbf{i,M}}$. The corresponding ‘aligned’ average activity modes (of size $20 \times \mathrm{T}_{\mathrm{all}} \times \mathrm{C}$, for experiment ‘i’) were either visualized individually for each experiment (Extended Data Fig. 6e), or by averaging across experiments (Extended Data Fig. 6d).

To evaluate the success of the alignment procedure, we computed a correlation coefficient Corr ($< \mathbf{z}^{i}_{(a)} >, < \mathbf{z}^{j}_{(b)} >$) for a given pair of aligned modes (indexed by $a$ and $b$, where $a, b \in$ {1,2,..,20}) across all possible pairs of experiments (indexed by $i$ and $j$), where $< \mathbf{z}^{i}_{(a)} >$ is the $a^{\text{th}}$ trial-averaged aligned activity time course vector (for both choices), of dimensionality $1 \times (\mathrm{T}_{\mathrm{all}} \times \mathrm{C})$. We then computed the median correlation coefficient across all pairs of dissimilar experiments ($i \neq j$) for each pair of modes and visualized the resulting mode x mode correlation matrix. The median correlation coefficient matrix (Extended Data Fig. 6g) displayed large values along the diagonal and almost zero values along the off-diagonals indicating that the aligned time-courses were much more similar across sessions than across modes.

#### 6.2. Dynamics Subspace Estimation

The second step of the data analysis pipeline involved using the aligned 20-dimensional neural response patterns to compute aligned residuals, which were then used to estimate the dynamics subspace (Extended Data Fig. 2, Step 2). The dynamics subspace is defined by a set of directions in the 20-dimensional aligned space, ordered according to their predictive power in explaining “future” variance in the aligned residuals based on the “past”, relative to any given time ‘t’ in the trial. This idea of finding “temporally predictive” directions has roots in subspace system identification (SSID) theory, which we review in Supplementary Math Note A. Specifically, SSID methods are well characterized for linear time-invariant systems[9]. In this study, we adapt these existing methods to the case of systems with linear time-varying dynamics as in Equation 11. This step of the pipeline consisted of a series of sub-steps that are described in detail below:

##### Computing aligned residuals

We computed single-trial residual trajectories by subtracting from each single trial trajectory its corresponding time-varying condition average trajectory (Extended Data Figs. 1a,2). The residuals were computed separately for each experiment using the corresponding aligned single trial response patterns $\mathbf{Z}^{\mathrm{i}}$ (where 'i' indexes experiments, see: Session alignment procedure). First, we *redefined* conditions separately for the two task epochs. For the decision epoch, we defined conditions based on the choice of the monkey *and* the motion strength of the random-dot stimulus (2 choices × 4 to 8 coherencies ≈ 8 to 16 conditions; since the number of distinct motion coherencies used varied slightly across different experiments for the two monkeys). For the movement epoch, we defined conditions based on the choice of the monkey and the length of the variable delay period preceding the 'go' cue by sorting trials recorded in each experiment into 5 different groups based on the length of the delay period (bin boundaries = [0 0.4 0.6 0.8 1.0 1.5] s) preceding the saccade. To ensure minimal overlap between the decision and movement epochs we excluded all trials that had an associated delay length < 400ms from the analysis. Thus, for the movement epoch, the above procedure yielded a total of 8 conditions in monkey T (2 choices × 4 delay length bins) and 6 conditions in monkey V (2 choices × 3 delay length bins; no trials in monkey V had a delay length larger than 1s) across all experiments.

Having defined conditions as above for the two task epochs, we computed the condition average trajectories specific to each epoch. For a given condition and epoch, we then removed the condition average trajectory from each of its corresponding single trial trajectories. This procedure resulted in 'aligned' residual data matrices $\tilde{\mathbf{Z}}^{\mathbf{i}}$ (i = 1, 2, …, P) for each experiment 'i', of dimensionality $20 \times \mathrm{T_{all}} \times \mathrm{K_i}$ (the residuals trajectories computed separately for the two epochs were concatenated along the time dimension), where $\mathrm{K_i}$ is the number of trials in the i$^{\text{th}}$ experiment. We then constructed 'pooled residual matrices' for each choice, $\tilde{\mathbf{Z}}_{\{\mathrm{choice}=1\}}$and $\tilde{\mathbf{Z}}_{\{\mathrm{choice}=2\}}$, by sorting the trials in each $\tilde{\mathbf{Z}}^{\mathbf{i}}$ based on the two choice conditions - choice 1 or choice 2, and pooling them across all 'P' experiments. All subsequent procedures were carried out *separately* on $\tilde{\mathbf{Z}}_{\{\mathrm{choice}=1\}}$ and $\tilde{\mathbf{Z}}_{\{\mathrm{choice}=2\}}$, unless otherwise indicated. Thus, for the sake of convenience, we drop the subscript and use $\tilde{\mathbf{Z}}$ to refer to the pooled, aligned matrix of residuals belonging to either choice.

##### Low-rank hankel matrix decomposition

SSID theory is centered around matrix decompositions of a special matrix known as the *hankel covariance* matrix[9,15,16]. Using $\tilde{\mathbf{Z}}$ we constructed a *time-varying* future-past hankel covariance matrix ($\mathbf{H_t}$) according to the mathematical definition presented in Equation 55 (see Supplementary Math Note A; Extended Data Fig. 2, Step 2). Specifically, we split the trials in $\tilde{\mathbf{Z}}$ (corresponding to a specific choice condition), into two random halves labelled the "training" set and "test" set, and used them to construct two distinct hankel matrices $\mathbf{H}_t^{train}$ and $\mathbf{H}_t^{test}$, for each time 't'. The order of the hankel matrix (given by *q* in Equation 55, see Supplementary Math Note A) that determines the number of "future" and "past" lags to use in the construction of $\mathbf{H}_\mathrm{t}$ was set to a value of 5. Increasing this value beyond 5 did not change the results of our analyses. We subjected $\mathbf{H}_t^{train}$ to a singular value decomposition as follows:

$$\mathbf{H}_t^{train} \quad = \mathbf{U}_t^{train}.\mathbf{S}_t^{train}.\mathbf{V}_t^{train'} \tag{17}$$

where, $\mathbf{U}_t^{train}$ and $\mathbf{V}_t^{train}$ are matrices whose columns define the left and right singular vectors of $\mathbf{H}_t^{train}$ respectively. $\mathbf{S}_t^{train}$ is a diagonal matrix whose non-zero elements are the corresponding singular values. The *r*-rank approximation of $\mathbf{H_t}^{train}$ was obtained by using a hard-thresholding of its singular values (diagonal entries of $\mathbf{S}_t^{train}$) as follows:

$$\mathbf{H}_{t,(r)}^{train} \quad = \mathbf{U}_{t,(r)}^{train}.\mathbf{S}_{t,(r)}^{train}.\mathbf{V}_{t,(r)}^{train'} \tag{18}$$

where, $\mathbf{U}_{t,(r)}^{train}$ and $\mathbf{V}_{t,(r)}^{train}$ are matrices containing the first *'r'* columns of $\mathbf{U}_t^{train}$ and $\mathbf{V}_t^{train}$ respectively. Similarly, $\mathbf{S}_{t,(r)}^{train}$ is a $\mathrm{r} \times \mathrm{r}$ diagonal matrix with the diagonal entries corresponding to the first 'r' diagonal entries of $\mathbf{S}_t^{train}$.

We used the r-rank approximation (Extended Data Fig. 2, Step 2) to compute a temporally averaged hankel matrix reconstruction error:

$$\mathrm{E_{hankel}} = \frac{1}{T - 2q + 1} \sum_{t=q+1}^{T-q+1} \| \mathbf{H}_t^{test} - \mathbf{H}_{t,(r)}^{train} \|_F^2 \tag{19}$$

where $\|.\|_F$ is the matrix frobenius norm and T is the total number of time bins in $\tilde{\mathbf{Z}}$ for a specific task epoch (either decision or movement). We computed $\mathrm{E_{hankel}}$ 20 times using different random splits of $\tilde{\mathbf{Z}}$ into "train" and "test" sets, for different values of the hankel rank (r). The average reconstruction error (averaged over 20 repeats) plotted as a function of r reached a minimum, before increasing for progressively larger values of r indicating that the hankel matrices were indeed approximately low-rank (Extended Data Fig. 8a). The optimal rank ($r_{opt}$) for a given choice condition (choice-1 or choice-2) and task epoch (decision or movement) combination was chosen to be the smallest value of 'r' for which the average reconstruction error was no more than one standard error above the minimum average reconstruction error (1 standard error rule). Thus, our procedure resulted in a single value of $r_{opt}$ for each task epoch and choice condition. While this single value may over/under estimate the approximate rank of the hankel matrix at a specific time ‘t’, we found that computing a different $r_{opt}$ for the hankel matrix at each ‘t’ also yielded similar results.

#### Estimating the dynamics subspace from observability matrices

Having obtained the optimal hankel rank ($r_{opt}$) in the previous step, we used it to estimate the first $r_{opt}$ left singular vectors of the time-varying hankel matrices constructed using $\tilde{\mathbf{Z}}$. These time-varying left singular vectors ($r_{opt}$ vectors for each ‘t’) were then used to compute the set of ordered directions that constituted the dynamics subspace. To do so, we adapted existing SSID methods that cater to time-invariant systems (see Supplementary Math Note A), to make them suitable for the time-varying dynamics model (Equation 11) that characterizes our definition of residual dynamics.

For this step, to compute the hankel matrices and the eventual dynamics subspace, we used a 5-fold cross validation approach that involved partitioning the trials in $\tilde{\mathbf{Z}}$ into 5 non-overlapping sets, with 4/5th of the trials constituting a “train” set ($\tilde{\mathbf{Z}}^{train}$) and the remaining 1/5th constituting the “test” set ($\tilde{\mathbf{Z}}^{test}$) for each fold.  Time-varying hankel matrices ($\mathbf{H_t}$) were computed using $\tilde{\mathbf{Z}}^{train}$ only, and subjected to an SVD resulting in matrices $\mathbf{U}_t^{train}$ , $\mathbf{S}_t^{train}$ as in Equation 17. We retained the first $r_{opt}$ (determined in the previous step) columns of $\mathbf{U}_t^{train}$ and the first $r_{opt}$ diagonal elements of the matrix $\mathbf{S}_t^{train}$ to define a time-dependent observability matrix $\widehat{\boldsymbol{\mathcal{O}}}_t$ (see Equation 59, Supplementary Math Note A) as shown below:

$$\widehat{\boldsymbol{\mathcal{O}}}_t \quad = \mathbf{U}_{t,(r_{opt})} . (\mathbf{S}_{t,(r_{opt})})^{\frac{1}{2}} \tag{20}$$

where, $\widehat{\boldsymbol{\mathcal{O}}}_t$ is a block matrix of size $(\mathrm{M} \times \mathrm{q}) \times \mathrm{r_{opt}}$, q = 5 is the order of the hankel matrix and M= 20 is the dimensionality of the aligned space. Similar to SSID for time-invariant systems, the first block row of $\widehat{\boldsymbol{\mathcal{O}}}_t$ specifies the *momentary* dynamics subspace at ‘t’, given by

$$\hat{\mathbf{C}}_t \quad = \widehat{\boldsymbol{\mathcal{O}}}_t(1{:}\mathrm{M}, :) \tag{21}$$

These momentary dynamics subspaces (defined at each ‘t’, separately for each task epoch and choice condition) were then combined across time, task epochs and choice conditions to define a single *time-invariant dynamics subspace* as specified by our model in Equation 11. To do so, we constructed a matrix $\widehat{\mathbf{C}_{\mathrm{all}}}$ by concatenating column-wise, the estimated momentary dynamics subspaces $\hat{\mathbf{C}}_t$ for all ‘t’ across both task epochs (decision or movement) and both choice conditions (choice *1* or *2)*

$$\widehat{\mathbf{C}_{\mathrm{all}}} = \left(\hat{\mathbf{C}}_{q+1}^{de,1} \dots \dots \hat{\mathbf{C}}_{T_{de}-q+1}^{de,1} \quad \hat{\mathbf{C}}_{q+1}^{mo,1} \dots \dots \hat{\mathbf{C}}_{T_{mo}-q+1}^{mo,1} \quad \hat{\mathbf{C}}_{q+1}^{de,2} \dots \dots \hat{\mathbf{C}}_{T_{de}-q+1}^{de,2} \quad \hat{\mathbf{C}}_{q+1}^{mo,2} \dots \dots \hat{\mathbf{C}}_{T_{mo}-q+1}^{mo,2}\right) \tag{22}$$

where, $\hat{\mathbf{C}}_{\mathrm{t}}^{\mathrm{de,\,j}}$ and $\hat{\mathbf{C}}_{\mathrm{t}}^{\mathrm{mo,\,j}}$are the momentary dynamics subspaces estimated for choice 'j' (j = 1 or 2), at time 't' in the trial during the decision (*de*) and movement (*mo*) epochs. In Equation 22, $T_{de}$ and $T_{mo}$ refers to the total number of time bins in the decision and movement epochs respectively and *q* denotes the order of the hankel matrix (Equation 55, see Supplementary Math Note A). The left singular vectors of $\widehat{\mathbf{C}_{\mathrm{all}}}$ , which by definition, span the *union* of the column spaces of all $\hat{\mathbf{C}}_t$ (across time, task epochs and choice conditions) specify a time-invariant dynamics subspace that is *shared* across time, task epochs and conditions. Henceforth, we denote the left-singular vectors of $\widehat{\mathbf{C}_{\mathrm{all}}}$ by $\mathbf{U}_{dyn}$, which is an orthonormal matrix of size $\mathrm{M} \times \mathrm{M}$. The $\mathrm{M}$ columns of $\mathbf{U}_{dyn}$ are ordered in terms of their relative importance in capturing temporally correlated variability in the residuals that result from the underlying dynamics (Fig. 4a, Extended Data Fig. 6f). In practice, only a subset of the columns of $\mathbf{U}_{dyn}$ may be sufficient to capture the latent residual dynamics across choice conditions and task epochs, whereby using more columns of $\mathbf{U}_{dyn}$ could lead to over-fitting. The retained columns of $\mathbf{U}_{dyn}$ (Fig. 4a, Extended Data Fig. 6f,8) in turn define the estimate of the observation matrix (**C**) in Equation 11. We determined the number of columns of $\mathbf{U}_{dyn}$ to retain (and thereby the dimensionality of the dynamics matrices $\mathbf{A}_{\mathrm{t}}$ in Equation 11) in the next step of the pipeline.

#### 6.3. Two-stage least squares (2SLS) state and dynamics estimation

We used the aligned residual trajectories ($\tilde{\mathbf{Z}}$) and the estimate of the dynamics subspace ($\mathbf{U}_{dyn}$) obtained in the previous step to (i) estimate the residual latent state ($\mathbf{x}_{\mathrm{t}}$ in Equation 11; Extended Data Fig. 2, Step 3) and, (ii) the time-varying dynamics matrices ($\mathbf{A}_{\mathrm{t}}$) that characterize the residual dynamics (Extended Data Fig. 2, Step 4), using a two-stage least squares(2SLS) approach based on instrumental variable regression, with steps (i) and (ii) above comprising the outputs of the first and second stage of the 2SLS regression respectively.

Before applying the 2SLS approach, we first obtained a d-dimensional 'noisy' estimate of the true underlying latent residual state, denoted by $\tilde{\mathbf{x}}_t$ as follows:

$$\tilde{\mathbf{x}}_{\mathrm{t}}^{train}(k) = (\mathbf{U}_{dyn}^{d})'\tilde{\mathbf{z}}_{\mathrm{t}}^{train}(k) \qquad (23)$$

where, $\tilde{\mathbf{z}}_{\mathrm{t}}^{train}(k)$ is a single residual vector indexed by time 't' and trial 'k' in $\tilde{\mathbf{Z}}^{train}$ and $\mathbf{U}_{\mathrm{dyn}}^{\mathrm{d}}$ are the first '*d*' columns of $\mathbf{U}_{\mathrm{dyn}}$ estimated using $\tilde{\mathbf{Z}}^{train}$. The hyper-parameter 'd' determines the dimensionality of the dynamics and is determined during the cross-validation procedure (Extended Data Fig. 2, Step 3).

We specifically make the distinction of referring to $\tilde{\mathbf{x}}_t$ in Equation 23 as a *'noisy'* estimate of the true latent residual state, because the projection defined in Equation 23 does not entirely eliminate aspects of "observation noise" in $\tilde{\mathbf{z}}_{\mathrm{t}}^{train}$ that lies within the column space of $\mathbf{U}_{\mathrm{dyn}}^{\mathrm{d}}$. This remaining observation noise in $\tilde{\mathbf{x}}_t$ makes the ordinary least squares (OLS) estimate of the dynamics matrix $\mathbf{A}_\mathrm{t}$ (derived directly using the $\tilde{\mathbf{x}}_t$, as suggested by Equation 11), biased and inconsistent. This is commonly referred to as the “error-in-variables” problem[17], in which the observation noise that corrupts $\tilde{\mathbf{x}}_t$ acts as a confounding variable, bringing about an *attenuation bias* in the estimate of $\mathbf{A}_\mathrm{t}$, as demonstrated in simulations of linear dynamical systems (Extended Data Fig. 3d). Such biases therefore make it difficult to interpret the properties of $\mathbf{A}_\mathrm{t}$, particularly its eigen/singular value spectrum, which are crucial for inferring the underlying neural computations.

Therefore, we instead used an approach based on instrumental variables (IV), commonly used in causal inference techniques to mitigate the deleterious effects of confounding variables[18]. The key underlying idea is to perform regression in two stages. In the first stage, we use suitable auxiliary variables (known as “instruments”) to obtain a de-noised estimate of the residual latent state, which is then used in the second stage to obtain unbiased and consistent estimates of $\mathbf{A}_\mathrm{t}$. Leveraging the Markovian nature of the dynamics, and making the critical assumption that the observation noise corrupting $\tilde{\mathbf{x}}_t$ is uncorrelated across time, allows us to use the past-lags of the noisy residual latent state $[\tilde{\mathbf{x}}_{t-1},\ \tilde{\mathbf{x}}_{t-2}\ \ldots \tilde{\mathbf{x}}_{t-l}]$ as valid "instruments" for the 2SLS procedure. Specifically, in the first stage we regress the explanatory variable ($\tilde{\mathbf{x}}_t$) against the past ’l’ lags (relative to ’t’) of the noisy latent residual state (instruments). In the second stage, we regress $\tilde{\mathbf{x}}_{t+1}$ against the ’denoised’ predictions of $\tilde{\mathbf{x}}_t$ obtained from the first stage.

The 2SLS can itself lead to biased estimates in the presence of ’weak’ instruments[19] i.e instrumental variables with low predictive power for the first stage of the 2SLS. Consequently, the choice of the hyper-parameters ’*d*’ (dimensionality of the residual latent state/dynamics) and ’*l*’ (the number of past lags in the first stage regression) are critical (Extended Data Fig. 2, Step 3). These hyper-parameters were determined using a grid search, as part of the 5-fold cross-validation approach (Extended Data Fig. 8b).

#### Residual latent state estimation

The first stage of the 2SLS procedure, involved estimating at each time-step, the parameter $\boldsymbol{\beta}_t^{l-}$ for the regression problem specified below:

$$\tilde{\mathbf{x}}_t^{train}(k) = \boldsymbol{\beta}_t^{l-}\tilde{\mathbf{x}}_{t,l-}^{train}(k) + \boldsymbol{\eta} \qquad (24)$$

Where, $\boldsymbol{\eta}$ is the noise term of the regression and $\tilde{\mathbf{x}}_{\mathrm{t,l-}}^{\mathrm{train}}(k)$ is a vector of size $(\mathrm{d} \times \mathrm{l}) \times 1$ constructed by vertically stacking the ’*l*’ past lags (relative to ’t’) of the noisy latent residual state for the k$^{th}$ trial in the training set as shown below:

$$\tilde{\mathbf{x}}_{t,l-}^{train}(k) = \begin{bmatrix} \tilde{\mathbf{x}}_{t-1}^{train}(k) \\ \tilde{\mathbf{x}}_{t-2}^{train}(k) \\ \vdots \\ \tilde{\mathbf{x}}_{t-l}^{train}(k) \end{bmatrix} \quad (25)$$

The OLS estimate of $\boldsymbol{\beta}_t^{l-}$ was then obtained as:

$$\widehat{\boldsymbol{\beta}}_t^{l-} \quad = (\widetilde{\mathbf{X}}_t^{train}\widetilde{\mathbf{X}}_{t,l-}^{train'})(\widetilde{\mathbf{X}}_{t,l-}^{train}\widetilde{\mathbf{X}}_{t,l-}^{train'})^{-1} \quad (26)$$

where, $\widetilde{\mathbf{X}}_t^{train}$ is a matrix of size $\mathrm{d} \times \mathrm{K_{train}}$ whose columns correspond to the noisy latent residual state $\tilde{\mathbf{x}}_t^{train}(k)$ for the different trials. Similarly, $\widetilde{\mathbf{X}}_{t,l-}^{train}$ is a matrix of size $(\mathrm{d} \times \mathrm{l}) \times \mathrm{K_{train}}$ constructed by collecting the stacked lag vectors $\tilde{\mathbf{x}}_{t,l-}^{train}(k)$ for all the corresponding trials. We performed the above regression *separately* for trials from the two task epochs and choice conditions.

To evaluate the specific choice of hyperparameters *d* and *l* (Extended Data Fig. 2, Step 3), we predicted each residual datapoint in the test set ($\tilde{\mathbf{Z}}^{test}$, remaining 1/5th of the data) using the parameters $\mathbf{U}_{dyn}^{d}$ and $\widehat{\boldsymbol{\beta}}_t^{l-}$ that were estimated using $\tilde{\mathbf{Z}}^{train}$. Similar to Equation 23, we first obtained a noisy latent residual state (denoted by $\tilde{\mathbf{x}}_{\mathrm{t}}^{test}(k)$) for each datapoint in $\tilde{\mathbf{Z}}^{test}$. We then evaluated predictions of the denoised/filtered residual state at time 't' for the kth trial in the test set, using $\widehat{\boldsymbol{\beta}}_t^{l-}$ as follows.

$$\hat{\tilde{\mathbf{x}}}_t^{test}(k) = \widehat{\boldsymbol{\beta}}_t^{l-}\tilde{\mathbf{x}}_{t,l-}^{test}(k) \quad (27)$$

where, $\tilde{\mathbf{x}}_{t,l-}^{test}(k)$ is constructed in a manner similar to Equation 25. We then back-projected the above *denoised* residual state *prediction* to obtain a corresponding prediction in the M-dimensional aligned space as follows:

$$\hat{\tilde{\mathbf{z}}}_t^{test}(k) = \mathbf{U}_{dyn}^{d}\hat{\tilde{\mathbf{x}}}_t^{test}(k) \quad (28)$$

These predictions were then used to compute a mean squared error metric defined as follows:

$$E_{fs} = \frac{1}{T_{pred}.K_{test}}\sum_{k}\sum_{t} \| \hat{\tilde{\mathbf{z}}}_t^{test}(k) - \tilde{\mathbf{z}}_t^{test}(k)\|_2^2 \quad (29)$$

The above error term was computed by pooling across both task epochs (i.e summation index 't' includes time bins from both, the decision and movement epochs) and, using only those time bins 't' that had an index greater than the maximum lag tested as part of the grid-search cross-validation. $T_{pred}$ corresponds to the total number of time-bins across both epochs used to compute the error measure and $K_{test}$ is the total number of trials in $\tilde{\mathbf{Z}}^{test}$.

We sampled different values of d and l on a two-dimensional grid of points and inspected the resulting average (across CV folds) and standard error of $E_{fs}$, for the different combinations of d and l. This average cross-validated mean-squared error measure revealed a tendency to over-fit for large values of 'd' and 'l' (Extended Data Fig.8b) as expected. We found that the average cross-validated mean squared error on the test set did not appreciably decrease for values of *d* beyond 5, and for values of *l* beyond 3. Across most datasets (combinations of different choices and task configurations), we found that values $d_{opt}$ = 8 and $l_{opt}$ = 3 or 4 were optimal choices of *d* and *l* respectively (Extended Data Fig. 8d). These optimal values were determined as the combination of *d* and *l* resulting in the smallest number of parameters in $\widehat{\boldsymbol{\beta}}_t^{l-}$, which had associated with them an average $E_{fs}$ value that was no more than one standard error above the minimum average $E_{fs}$ (1 standard error rule). Ultimately, we used $d_{opt}$ = 8 and $l_{opt}$ = 3 for estimating the dynamics matrices across all the datasets in both monkeys, despite there being some variability in these hyper-parameter values across different datasets. Such a choice was motivated by the need for consistency, which allowed us to compare and pool results across different datasets. The 8-dimensional dynamics subspace $\mathbf{U}_{dyn}^{d_{opt}}$ explained 68% (Monkey T, median across 164 experiments; Extended Data Fig.7c) and 55% (Monkey V, median across 80 experiments) of the variance in the trial-averaged trajectories in the high-dimensional neural space (computed prior to alignment of sessions). In comparison, the first 20 aligned dimensions ($\mathbf{U}_{\mathrm{i,M=20}}^{\perp}$), which are optimized to capture variance in the condition-averaged trajectories across all experiments, captured 87% of overall variance in monkey T (median across 164 experiments) and 73% (median across 80 experiments) in Monkey V (Extended Data Fig 4).

#### Time-varying dynamics estimation

Having estimated the optimal value of the hyper-parameters $d_{opt}$ and $l_{opt}$ of the first stage of the 2SLS regression, we recomputed the dynamics subspace ($\mathbf{U}_{dyn}^{d}$) and, the denoised residual latent state ($\hat{\tilde{\mathbf{x}}}_t(k)$ from Equation 27) using all datapoints in $\tilde{\mathbf{Z}}$. The corresponding matrix of denoised residual latent states is denoted by $\hat{\tilde{\mathbf{X}}}$ (of dimensionality $d_{opt} \times \mathrm{T_{all}} \times \mathrm{K}$, where K is the total number of trials in $\tilde{\mathbf{Z}}$). The second stage of the 2SLS procedure involved regressing the denoised residual latent states at time 't' (denoted by $\hat{\tilde{\mathbf{X}}}_t$) against the 'noisy' residual latent states at 't+1' (denoted by $\tilde{\mathbf{X}}_{t+1}$), in order to obtain the residual dynamics matrices $\mathbf{A}_t$. To solve such a regression problem, we optimized the following penalized least squares objective that imparted smoothness in the temporal sequence of $\mathbf{A}_t$[20]

$$\mathcal{L} = \sum_t \| \tilde{\mathbf{X}}_{t+1} - \mathbf{A}_t \hat{\tilde{\mathbf{X}}}_t \|_F^2 + \alpha \| \mathbf{A}_{t+1} - \mathbf{A}_t \|_F^2 \qquad (30)$$

where, $\alpha$ was a regularization parameter (Extended Data Fig. 2, Step 4) and 't' indexed the time-bins within a single task epoch (either decision or movement epoch). The above regularized least squares optimization problem was solved in closed-form to obtain $\mathbf{A}_t$. The optimal value of $\alpha$ was determined in a separate 5-fold cross-validation procedure. Very small values of $\alpha$ exhibited over-fitting while very large values led to under-fitting (Extended Data Fig. 8c, shows an example configuration in monkey T). We found considerable differences in the optimal $\alpha$ value between the two monkeys and across the two task epochs (Extended Data Fig. 8e), with larger values (implying higher smoothing) in the decision epoch compared to the movement epoch, and in monkey V compared to monkey T.

Across all datasets in both monkeys, we chose $\alpha_{opt}$ = 200 for the fits in the decision epoch and $\alpha_{opt}$ = 50 for the final fits of the dynamics matrices. Although the cross-validation results suggested larger values of $\alpha_{opt}$ for monkey V, we decided to use values comparable to those used in monkey T to simplify comparisons between monkeys.

#### 7. Analysis of residual dynamics

Having determined the relevant hyper-parameters ($d_{opt}$, $l_{opt}$ and $\alpha_{opt}$ ) of the 2SLS procedure, we used them in a final step to estimate the time-varying dynamics matrices $\mathbf{A}_t$ (Equation 30) using all the datapoints in $\tilde{\mathbf{Z}}$, separately for each choice condition and task epoch. We analyzed the eigenvalue and singular value spectra of the estimated $\mathbf{A}_{\mathrm{t}}$ to make inferences about the underlying dynamics. In order to plot the time-variation in the eigenvalue spectra, we sorted the eigenvalues and the associated eigenvectors at each time step such that they were maximally consistent with those at the preceding time-step. The eigenvalues/eigenvectors at the very first time-step were sorted in descending order of their absolute values. This was done using a modified version of the 'eigenshuffle' MATLAB script freely available online (*https://www.mathworks.com/matlabcentral/fileexchange/22885-eigenshuffle*). A similar procedure was used to sort the time-varying singular values and its associated right and left singular vectors.

We computed the time constant of the dynamics (Fig. 4b,e) directly from the absolute value of the eigenvalues of $\mathbf{A}_{\mathrm{t}}$ as follows:

$$\tau_t^j = \frac{\Delta t}{\log(|\lambda_t^j|)} \tag{31}$$

where, $\lambda_{\mathrm{t}}^{\mathrm{j}}$ is the eigenvalue at time 't' associated with the $\mathrm{j}^{th}$ eigenmode and $\Delta$t is the time step (= 45 ms - size of time bin).

We analyzed the imaginary components of the eigenvalues of $\mathbf{A}_\mathrm{t}$ to obtain evidence for rotational dynamics (Fig. 4d,g). The natural oscillation frequency associated with the $j^{th}$ eigenmode is related to the eigenvalues of the dynamics matrix as follows:

$$f_t^j = \frac{\angle\lambda_t^j}{2\pi\Delta t} \tag{32}$$

where $\angle\lambda_t^j$ is the angular phase of the $j^{th}$ complex eigenvalue (computed using the atan2 function in MATLAB).

We computed the largest eigenvalue magnitude ($|\lambda_t^{max}|$) and singular value ($|\sigma_t^{max}|$) at time 't' as:

$$\begin{aligned} |\lambda_t^{max}| &= \max_j |\lambda_t^j| \\ |\sigma_t^{max}| &= \max_j |\sigma_t^j| \end{aligned} \tag{33}$$

where 'j' indexes the eigen/singular modes (Data Fig. 4b-c,e-f).

Finally, we quantified the magnitude of non-normality of the dynamics (Fig. 4h) based on a previously proposed measure[21] comparing the singular values and absolute eigenvalues of $\mathbf{A}_\mathrm{t}$:

$$d_F(\mathbf{A}_t) = \frac{\sqrt{\sum_j (\sigma_t^j)^2 - \sum_j (|\lambda_t^j|)^2}}{\sqrt{\sum_j (|\lambda_t^j|)^2}} \tag{34}$$

where, $\sigma_t^j$ and $\lambda_t^j$ are the $j^{th}$ singular and eigenvalue respectively.

### 8. Average task activity subspaces and their relationship to residual dynamics

#### 8.1. Computing average task activity subspaces

We used the 20-dimensional neural response patterns, obtained after session alignment ($\mathbf{Z}$, output of Step 1 in Extended Data Fig. 2), to compute four distinct task activity subspaces optimized to capture variance in the aligned, trial-averaged trajectories due to choice (condition-dependent), time (condition-independent) and rotations (Fig. 3c-d). These task activity subspaces, denoted as $\mathbf{U}_{task}^j$ ($j \in \{\mathrm{choice}, \mathrm{time}, \mathrm{jPC}_{12}, \mathrm{jPC}_{34}\}$, were computed separately for the decision (aligned to dots onset) and movement epochs (aligned to movement onset).

To compute the "choice" and "time" subspaces, we pooled session-aligned single trials from all experiments of a task configuration, and sorted them into one of two conditions depending on the choice of the monkey (choice 1 or choice 2), resulting in two condition trial-averaged response matrices $\langle\mathbf{Z}\rangle_{\text{choice}=1}$ and $\langle\mathbf{Z}\rangle_{\text{choice}=2}$ of dimensionality $\mathrm{M} \times \mathrm{T}_{epoch}$, with $\mathrm{T}_{epoch}$ indicating the number of time bins in the task epoch under consideration and M = 20, the number of aligned dimensions. We used $\langle\mathbf{Z}\rangle_{\text{choice}=1}$ and $\langle\mathbf{Z}\rangle_{\text{choice}=2}$ to compute a normalized "difference response matrix" ($\boldsymbol{D}$) and a "sum response matrix" ($\boldsymbol{S}$) through elementwise subtraction and addition respectively:

$$\begin{aligned} \boldsymbol{D} &= 0.5 * (\langle\mathbf{Z}\rangle_{\text{choice}=1} - \langle\mathbf{Z}\rangle_{\text{choice}=2}) \\ \boldsymbol{S} &= 0.5 * (\langle\mathbf{Z}\rangle_{\text{choice}=1} + \langle\mathbf{Z}\rangle_{\text{choice}=2}) \end{aligned} \tag{35}$$

The first two principal components of the difference response matrix ($\boldsymbol{D}$) together constituted the "choice" subspace, since by definition, they captured most of the variance in neural response patterns due to *differences* between choices. Similarly, the first two principal components of the sum response matrix ($\boldsymbol{S}$) together constituted the "time" subspace, since, they captured maximal variance due to choice-independent components of neural activity.

To compute the jPC subspaces, we temporally smoothed the session-aligned single trials (box filter, width = 180ms) prior to computing the trial-averaged response matrices $\langle\mathbf{Z}\rangle_{\text{choice}=1}$ and $\langle\mathbf{Z}\rangle_{\text{choice}=2}$. The jPC vectors in the decision and movement epochs were estimated using trial-averaged responses restricted to a narrow time window - [500 1000]ms aligned to dots onset, and [-250 250]ms aligned to movement onset respectively. The above choice of time windows was motivated by the observation of rotational dynamics in the residuals late in the decision epoch in one monkey (monkey T, Fig. 4d) and, previous reports of rotational dynamics during movement[22]. As in the original jPCA study[22], we performed a pre-processing PCA step which retained the top 4 principal component vectors (computed jointly on $\langle\mathbf{Z}\rangle_{\text{choice}=1}$ and $\langle\mathbf{Z}\rangle_{\text{choice}=2}$, restricted to the above time windows). However, unlike the original study[22], we did not remove the condition-independent components of neural activity prior to applying the jPCA algorithm. This procedure resulted in two distinct, orthogonal jPC planes ($\text{jPC}_{12}$ and $\text{jPC}_{34}$). Each jPC plane was defined by a pair of complex-conjugate jPC vectors ($\mathbf{v}_1$ and $\mathbf{v}_2$). To determine the projection of the responses into the jPC subspace, we computed a pair of suitably normalized real-valued projection vectors $\mathbf{u}_1$ and $\mathbf{u}_2$ as: $\mathbf{u}_1 = \mathbf{v}_1 + \mathbf{v}_2$ and $\mathbf{u}_2 = \mathrm{j} * (\mathbf{v}_1 - \mathbf{v}_2)$, which spanned the same subspace as $\mathbf{v}_1$ and $\mathbf{v}_2$. The imaginary components of the eigenvalues associated with $\mathbf{v}_1$ and $\mathbf{v}_2$ directly specified the natural frequency of rotation associated with that jPC plane. The jPC planes were ordered in descending order of their associated rotation frequency.

Only the two jPC planes ($\text{jPC}_{12}$ and $\text{jPC}_{34}$) were constrained to be mutually orthogonal. The range (computed across task configurations) of the largest subspace angle (in degrees) between any pair of task activity subspaces ( $\mathbf{U}_{task}^{i}$ and $\mathbf{U}_{task}^{j}$) in a given task epoch is summarized in the table below:

| | $\angle jPC_{12}, choice$ | $\angle jPC_{12}, time$ | $\angle jPC_{34}, choice$ | $\angle jPC_{34}, time$ | $\angle choice, time$ |
|---|---|---|---|---|---|
| Monkey 'T' decision epoch | [76.8, 85.0] | [57.3, 64.3] | [34.2, 50.7] | [73.9 88.2] | [76.3 89.2] |
| Monkey 'T' movement epoch | [58.5, 83.8] | [21.1, 39.1] | [31.1, 41.9] | [62.6, 79.6] | [79.3, 88.9] |
| Monkey 'V' decision epoch | [79.8, 87.5] | [78.6, 85.3] | [18.2, 49.7] | [74.7, 87.2] | [79.2, 88.2] |
| Monkey 'V' movement epoch | [65.1, 83.9] | [16.4, 36.1] | [42.6, 61.2] | [61.9, 89.0] | [76.2, 85.4] |

Importantly, the task-activity subspaces $\mathbf{U}_{task}^{j}$ capture variance in the aligned, trial-averaged trajectories, but, need not perfectly align with the 8-dimensional dynamics subspace $\mathbf{U}_{dyn}^{d_{opt}}$ computed using the residuals. To assess the extent to which the activity in the 2-dimensional task-activity subspaces (Fig. 3) unfold within the dynamics subspace, we computed the fraction of total variance in a given task activity subspace that can be attributed to activity unfolding within the dynamics subspace as follows:

$$\frac{\mathrm{Tr}\left(\mathrm{Cov}\left(\mathbf{U}_{dyn}^{d_{opt}}\left(\mathbf{U}_{dyn}^{d_{opt}\,\prime}\,\mathbf{U}_{task}^{j}\right)\mathbf{U}_{task}^{j\,\prime}\langle\mathbf{Z}\rangle\right)\right)}{\mathrm{Tr}\left(\mathrm{Cov}\left(\mathbf{U}_{task}^{j}\mathbf{U}_{task}^{j\,\prime}\langle\mathbf{Z}\rangle\right)\right)}$$

where $\mathrm{Tr}(.)$ is the matrix trace operator, $\mathrm{Cov}(.)$ corresponds to the covariance matrix of the argument, and $\langle\mathbf{Z}\rangle$ is the matrix of aligned, condition-averaged trajectories (dimensionality: $\mathrm{M} \times (\mathrm{T}_{epoch} \times\ \mathrm{C})$, where M = 20 and C = 2)

We computed a null distribution corresponding to the above fraction by replacing the numerator of the above expression by the quantity: $\mathrm{Tr}\left(\mathrm{Cov}\left(\mathbf{U}_{dyn}^{d_{opt}}\left(\mathbf{U}_{dyn}^{d_{opt}\,\prime}\,\mathbf{U}_{rand}\right)\mathbf{U}_{task}^{j\,\prime}\langle\mathbf{Z}\rangle\right)\right)$, where $\mathbf{U}_{rand}$ is a pair of a random, orthogonal directions that define a 2D subspace embedded within the 20-dimensional aligned space. The null distribution (obtained by randomly sampling $\mathbf{U}_{rand}$ 5000 times) provides the range of possible values for the above fraction due to chance alignment of the 8-dimensional dynamics subspace with an arbitrary 2D subspace in the 20-dimensional aligned space. The fraction of variance explained was in the range 0.66-0.94 (median = 0.85) for Monkey T, with 31/32 (32/32) datapoints lying beyond the 99$^{th}$ (95$^{th}$) percentile of the null distribution). The range was 0.41-0.95 (median = 0.72) for Monkey V, with 25/32 (28/32) datapoints beyond the 99th (95th) percentile of the null distribution). These findings imply that the components of the dynamics revealed by projections onto the task activity subspaces largely unfold within the dynamics subspace.

#### 8.2. Comparison of residual eigenvectors to task activity subspaces

In order to compare subspaces that capture the residual dynamics to those that capture the trial-averaged trajectories, we computed the alignment between each task activity subspace (i.e $\mathbf{U}_{task}^{j}$ , $j \in \{\mathrm{choice}, \mathrm{time}, \mathrm{jPC}_{12}, \mathrm{jPC}_{34}\}$) and the eigenvectors of the estimated residual dynamics, separately within each task epoch. Further, we used a general linear model to quantify the relationship between the alignment and the corresponding eigenvalues (Fig. 5). Specifically, for every estimated real-valued eigenvalue (pooled across dimensions, times in trial, and saccade targets/choices), we computed the subspace angle between a chosen task activity subspace (a 2D subspace) and the associated real-valued eigenvector. For every estimated complex-conjugate eigenvalue pair, we computed a pair of subspace angles between a 2-dimensional eigenplane spanned by a pair of real-valued projection vectors $\mathbf{u}_1$ and $\mathbf{u}_2$ (computed as described previously, i.e $\mathbf{u}_1 = \mathbf{v}_1 + \mathbf{v}_2$ and $\mathbf{u}_2 = \mathrm{j} * (\mathbf{v}_1 - \mathbf{v}_2)$, where $\mathbf{v}_1$ and $\mathbf{v}_2$ are the complex-conjugate eigenvector pair) and the average task activity. Prior to computation of the subspace angles, we projected each eigenvector/eigenplane through the columns of $\mathbf{U}_{\mathrm{dyn}}$ (estimate of the dynamics subspace), which ensured that eigenvectors shared the same dimensionality as the average task activity subspaces, which were originally defined in the 20-dimensional space of aligned response patterns.

We quantified the relationship between the eigenvalues and the alignment of the corresponding eigenvector with the task activity subspace using a linear model (separately for each task epoch). Specifically, we regressed each subspace angle indexed by a given task activity subspace (y-axis, Fig 5e) against the eigenvalue magnitude (|ev|, x-axis in Fig. 5e, top) and the associated rotation frequency (freq, x-axis in Fig 5e, bottom) as shown below:

$$alignment(EV, j) = \beta_1^j |ev| + \beta_2^j freq$$

where ‘j’ indexes the individual task -activity subspaces ( $j \in \{\mathrm{choice}, \mathrm{time}, \mathrm{jPC}_{12}, \mathrm{jPC}_{34}\}$ ). The regression coefficients were estimated using least squares and we also reported the 95% normal confidence intervals for each regression coefficient (errorbars in Fig. 5f).

### 9. Validation of robustness of analysis pipeline

#### 9.1. Estimating bias in the estimates of residual dynamics

The choice of bin-size for binning spike-counts was critical in order to avoid biases in the estimates of residual dynamics (Extended Data Fig. 9). To illustrate the effect of bin-size on the quality of the estimated residual dynamics, we simulated data from a continuous-time, *time-invariant* linear dynamical system with a gaussian observation model:

$$\begin{aligned} \dot{\mathbf{x}}_t &= \mathbf{A}\mathbf{x}_t + \mathbf{b} + \boldsymbol{\epsilon}_t \\ \mathbf{y}_t &= \mathbf{C}\mathbf{x}_t + \mathbf{d} + \boldsymbol{\eta}_t \\ \mathbf{x}_1 &\sim \mathcal{N}(0, \mathbf{Q}_0) \\ \boldsymbol{\epsilon}_t &\sim \mathcal{N}(0, \mathbf{Q}) \\ \boldsymbol{\eta}_t &\sim \mathcal{N}(0, \mathbf{R}) \end{aligned} \tag{36}$$

where, $\mathcal{N}(.,.)$ denotes a normal distribution. We simulated 5000 single-trial trajectories from a system with 3 latent dimensions (Dim($\mathbf{x}$) = 3) and 20 observed dimensions (Dim($\mathbf{y}$) = 20) in 1ms time steps for a total of 1500 time steps. The elements of the eigenvector matrix of $\mathbf{A}$ were sampled randomly from a standard normal distribution and the three eigenvectors were orthogonalized (normal dynamics) to have unit norm. The eigenvalues of $\mathbf{A}$ were set to (-2, -4, -6) indicating stable and strongly decaying dynamics. The latent input vector $\mathbf{b}$ was set to a $[2\ 2\ 2]^{\mathrm{T}}$. The covariance of the latent noise ($\boldsymbol{\epsilon}_t$) was set to a scaled identity matrix $\sigma^2\mathbf{I}$, where values of $\sigma^2$ (grey lines, Extended Data Fig. 9a) were swept across two orders of magnitude to assess how latent variance affects the estimates of the residual dynamics. The elements of the observation matrix $\mathbf{C}$ were sampled randomly from a standard normal distribution followed by an orthogonalization step using the QR decomposition. Thus, observation matrix $\mathbf{C}$ was a random orthogonal matrix**.** The elements of the baseline input $\mathbf{d}$ were sampled from a uniform distribution between $[0,8]$. The observation noise matrix $\mathbf{R}$ was set to be diagonal with the diagonal elements sampled from a uniform distribution between $[0,0.05]$. The initial noise covariance was obtained by solving the continuous time Lyapunov equation in steady-state.

We estimated the dynamics matrix ($\mathbf{A}_{\mathrm{t}}$) using our estimation procedure for a range of different bin-sizes. First, we computed residuals by removing the average trajectory from the single trial trajectories and then "re-binned" the residuals in non-overlapping bins of different sizes, chosen from the set $[2\ 3\ 5\ 10\ 15\ 30\ 40\ 60]$ ms. We chose a hankel order (q in Equation 55) of 5 and did not optimize the hyper-parameters $l$ and $\alpha$ during the estimation procedure. Instead, we chose these hyper-parameters sensibly while ensuring that they were consistent across the different bin size categories. For instance, $\alpha$ was chosen to be large (= $10^6$ ) in order to facilitate smoothness in the time-varying dynamics matrices, thus ensuring time-invariant fits. Additionally, we chose $l$ such that it roughly translated into equal units of time (in seconds) for the different bin sizes.

We assessed the effect of bin-size on the estimated dynamics matrices in terms of their eigenvalues. Importantly, eigenvalues for different bin-sizes cannot be compared directly, as discretizing a continuous dynamical system trivially results in eigenvalues that depend on the duration of the underlying time-step (here the bin size). The same dynamical system, when expressed at step-sizes $\Delta t_j$ and $\Delta t_{ref}$, would result in eigenvalues $\hat{\lambda}_{\Delta t_j}$ and $\hat{\lambda}^{(j)}_{\Delta t_{ref}}$ related by the following scaling relation:

$$\hat{\lambda}^{(j)}_{\Delta t_{ref}} = (\hat{\lambda}_{\Delta t_j})^{\frac{\Delta t_{ref}}{\Delta t_j}} \tag{37}$$

To discount these trivial differences, we thus transformed each eigenvalue $\hat{\lambda}_{\Delta t_j}$ obtained for bin size $\Delta t_j$ into an 're-binned' eigenvalue $\hat{\lambda}^{(j)}_{\Delta t_{ref}}$ expected for a reference bin size $\Delta t_{ref}$ = 40ms.

We compared the estimated, 're-binned' eigenvalues to the 'ground truth' eigenvalue expected for a bin size of $\Delta t_{ref}$, obtained by discretizing the continuous time dynamics update equation of the model (Extended Data Fig. 9a). Importantly, the absolute value of $\hat{\lambda}^{(j)}_{\Delta t_{ref}}$ asymptotically converged to the ground truth for increasing bin sizes, meaning that for large bin sizes the estimates of the dynamics matrix converged to an unbiased estimate. This convergence was independent of the specific choice of $\Delta t_{ref}$.

We observed a similar asymptotic convergence of the re-binned eigenvalue for increasing bin sizes in the measured neural data (Extended Data Fig. 9b), and used it to determine an optimal bin size for which estimates of the dynamics can be expected to be unbiased. We re-binned the recorded spiking data for monkey T in bin sizes of $[15\ 30\ 45\ 60\ 90]$ ms. After obtaining binned spike counts for these different bin sizes, we aligned responses from different experiments. Specifically, we did not re-compute the session alignment (Step 1 of Extended Data Fig. 2), but rather, we projected the single trial, binned spike counts corresponding to each bin size into the precomputed aligned basis $\mathbf{U}^{\perp}_{\mathbf{i,M}}$ (M = 20, see Equations 14 & 15) determined for a bin size of 45ms. We then recomputed the residuals using the pooled (across experiments) trials in the 20-dimensional 'aligned' space and re-estimated the dynamics matrix $\mathbf{A}_t$.

As in the simulated models, we did not seek to optimize any of the hyper-parameters (d, l and $\alpha$, Steps 3 and 4, Extended Data Fig. 2) involved in the estimation procedure, as the goal of this analysis was to understand how bin size alone effects the estimated eigenvalues. We fixed the dimensionality of the dynamics subspace (d) to a value of 8 and the smoothness regularization parameter ($\alpha$) to a value of 200 and 50 for the decision and movement epochs respectively. These values were the optimal ones as determined by cross-validation for residuals binned in 45 ms bins (Extended Data Fig. 8 and Fig. 4). Likewise, we chose the lag parameter $l$ separately for each bin-size such that it roughly translated into equal units of time for the different bin sizes, for example, $l$ = 3 for bin size 45ms (=135ms long window in the past); $l$ = 2 for bin size 60ms (=120ms long window in the past). We then compared rebinned eigenvalues of $\mathbf{A}_\mathrm{t}$ expected under a reference bin size ($\Delta\mathrm{t}_\mathrm{ref}$) of 15ms (Equation 37).

We focused on the rebinned eigenvalues in the two temporal windows showing the most pronounced time-dependencies (Fig 4): $\mathrm{t} \in [200\,400]$ms aligned to dots onset and $\mathrm{t} \in [-150\,250]$ms aligned to movement onset. We temporally averaged the absolute value of the rebinned eigenvalues in these two windows, and found that they asymptotically converged for bin sizes greater than 30ms (Extended Data Fig. 9b). This asymptotic convergence was observed across all 8 eigen-modes in both task conditions (choice 1 or choice2) and task epochs (decision or movement), and was consistent across the different task configurations in monkey T (Extended Data Fig. 9b) and V (not shown). Based on these findings, a bin size of 45ms was well motivated for our analyses.

#### 9.2. Qualitative estimates of goodness of fit

The cross-validation procedure (Extended Data Fig. 8) used to determine the optimal values of the hyperparameters (*d, l,* $\alpha$) of the pipeline provided the fit performance on held-out trials in terms of the mean squared error (m.s.e). Since residual dynamics is estimated using a two-stage least squares (2SLS) approach, we obtained two distinct cross-validated m.s.e values corresponding to each stage (first stage: see Extended Data Fig 8b; second stage: see Extended Data Fig 8c – shown only for a single configuration in Monkey T). These cross-validated m.s.e values translated to a coefficient of determination ($R^2$) of 0.0367 (mean across all task configurations and choices, std = 0.0064) and 0.0390 (std = 0.0029) for the first stage regression, and 0.0577 (mean across all task configurations, choices and task epochs, std = 0.014) and 0.065 (std = 0.013) for the second stage regression in Monkey T and Monkey V respectively.

It was difficult to gauge the 'goodness of fit' based on the above $R^2$ values, as their relatively small magnitude could have been due to unstructured observation noise (which cannot be predicted by any model) dominating the variability in the residuals. To determine if this was indeed true, we simulated residuals from a time-varying linear dynamical system (Equation 11) where the dynamics matrix ($\mathbf{A}_\mathrm{t}$) and the observation matrix ($\mathbf{C} \equiv \mathbf{U}_{dyn}^{d_{opt}}$) were matched to the estimates of residual dynamics and dynamics subspace obtained using the neural data.

To simulate residuals from the model (Equation 11), we also required estimates of the latent noise covariance ($\mathrm{Cov}(\boldsymbol{\epsilon}_t) = \mathbf{Q}$), the variance in the initial latent state ($\mathrm{Cov}(\tilde{\mathbf{x}}_0) = \mathbf{Q}_0$), and the observation noise covariance ($\mathrm{Cov}(\boldsymbol{\eta}_t) = \mathbf{R}$). These parameters were obtained from the aligned neural residuals ($\tilde{\mathbf{z}}_t$) by maximizing the likelihood under the model as follows:

$$\widehat{\mathbf{R}} \quad = \frac{1}{(T-l).K}\,\mathrm{diag}(\sum_{t=l+1}^{T}\sum_{k=1}^{K}[\,\tilde{\mathbf{z}}_t(k) - \mathbf{U}_{dyn}^{d_{opt}}\hat{\hat{\mathbf{x}}}_t(k)]\left[\tilde{\mathbf{z}}_t(k) - \mathbf{U}_{dyn}^{d}\hat{\hat{\mathbf{x}}}_t(k)\right]')$$

$$\widehat{\mathbf{Q}}_0 \quad = \frac{1}{\mathrm{K}}\sum_{k=1}^{K}\hat{\hat{\mathbf{x}}}_{l+1}\,(k)\hat{\hat{\mathbf{x}}}_{l+1}(k)'$$

$$\widehat{\mathbf{Q}} \quad = \frac{1}{d.(T-l-1).K}\,Tr\left(\sum_{t=l+1}^{T-1}\sum_{k=1}^{K}[\,\hat{\hat{\mathbf{x}}}_{t+1}(k) - \mathbf{A}_t\hat{\hat{\mathbf{x}}}_t(k)]\left[\hat{\hat{\mathbf{x}}}_{t+1}(k) - \mathbf{A}_t\hat{\hat{\mathbf{x}}}_t(k)\right]'\right)$$

where, $\hat{\hat{\mathbf{x}}}_t(k)$ is the denoised residual latent state at time 't' on trial 'k' obtained from the first stage of the 2SLS procedure (Equation 27, see also Equation 30).

Using the estimates of the dynamics matrix ($\mathbf{A}_{\mathrm{t}}$), dynamics subspace ($\mathbf{U}_{dyn}^{d_{opt}}$) and, the noise covariances ($\widehat{\mathbf{R}}, \widehat{\mathbf{Q}}, \widehat{\mathbf{Q}}_{\mathbf{0}}$ ) derived above, we simulated residual responses (using Equation 11) with a matched number of trials and overall level of variability. We then computed the coefficients of determination for these simulated responses, under the idealized assumption that our analysis pipeline works perfectly, i.e. is able to perfectly retrieve the dynamics (second stage regression) and the de-noised residual latent states at each time (first stage regression). We reasoned that the resulting coefficients of determination (denoted by $\mathbf{R}^2_{\text{sim-fs}}$ and $\mathbf{R}^2_{\text{sim-ss}}$ in the equations below) provide a realistic benchmark, if not a strict upper limit, to the fit quality that one can hope to obtain in the context of large observation noise.

To compute $\mathbf{R}^2_{\text{sim-fs}}$ and $\mathbf{R}^2_{\text{sim-ss}}$, we projected the simulated 20-dimensional observations ($\tilde{\mathbf{z}}_t{}^{sim}(k)$) into the estimated dynamics subspace (using Equation 23) and computed the amount of variance explained in the resulting projection by (i) the true latent state of the simulated model (denoted by $\tilde{\mathbf{x}}_t{}^{sim}(k)$), and by (ii) a 'noise-free' analogue of the latent state obtained by propagating the true latent state through the corresponding estimated dynamics matrix. The former ($\mathbf{R}^2_{\text{sim-fs}}$) provides a benchmark for comparing the coefficient of determination obtained for the first stage of the 2SLS procedure, whereas the latter ($\mathbf{R}^2_{\text{sim-ss}}$) provides the same for the second stage of the 2SLS procedure. Mathematically, these quantities were defined as follows:

$$\mathrm{R}^2_{sim-fs} \; = \; 1.0 - \frac{\sum_t\sum_k\left(\mathbf{U}_{dyn}^{d_{opt}'}.\,\tilde{\mathbf{z}}_t{}^{sim}(k) - \;\tilde{\mathbf{x}}_t{}^{sim}(k)\right)^2}{\sum_t\sum_k\left(\mathbf{U}_{dyn}^{d_{opt}'}\,\tilde{\mathbf{z}}_t{}^{sim}(k)\right)^2}$$

$$\mathrm{R}^2_{sim-ss} = 1.0 - \frac{\sum_t \sum_k \left(\mathbf{U}_{dyn}^{d_{opt}'} . \tilde{\mathbf{z}}_t{}^{sim}(k) - \mathbf{A}_{\mathrm{t}-1} \tilde{\mathbf{x}}_{t-1}{}^{sim}(k)\right)^2}{\sum_t \sum_k \left(\mathbf{U}_{dyn}^{d_{opt}'} \tilde{\mathbf{z}}_t{}^{sim}(k)\right)^2}$$

where, $\tilde{\mathbf{x}}_t{}^{sim}(k) = \mathbf{A}_{\mathrm{t}-1} \tilde{\mathbf{x}}_{t-1}{}^{sim}(k) + \hat{\boldsymbol{\epsilon}}_{t-1}$, and $\hat{\boldsymbol{\epsilon}}_{t-1}$ is a sample from a multi-variate gaussian with covariance $\widehat{\mathbf{Q}}$.

We found that the range of values of $\mathrm{R}^2_{sim-fs}$ and $\mathrm{R}^2_{sim-ss}$ qualitatively matched the range of values of the cross-validated coefficient of determination obtained for the data reported above. For the first stage regression, we obtained $\mathrm{R}^2_{sim-fs}$ = 0.0738±0.011 for Monkey T and 0.0958±0.33 for Monkey V (mean ±std across task configurations and choices). For the second stage regression, we obtained $\mathrm{R}^2_{sim-ss}$ = 0.0512±0.0171 for Monkey T and 0.0674±0.0157 for Monkey V (mean ±std across task configurations, task epochs and choices). The similarity of these simulated $R^2$ values to those observed in the real data suggest that the low coefficient of determination is likely a consequence of residuals being dominated by uncorrelated observation noise.

### 10. Simulated Models

We validated our approach on simulated data from a diverse set of models, each of which at its essence was a linear or non-linear dynamical system, potentially driven by time-varying inputs (Fig. 1c-d; Extended Data Fig. 1). We grouped the different simulated models into four distinct categories, which we describe in detail below.

#### 10.1 Models of decisions and movement

We simulated single trial responses from 3 different "models of decisions" and 3 different "models of movement" that had distinct intrinsic dynamics and time-varying external inputs, but that resulted in similar condition average trajectories (Fig. 1c-d). The purpose of these simulations was to demonstrate the overarching motivation behind analyzing neural residuals. The nature of the intrinsic dynamics and the temporal structure of the inputs were distinct across the six models, with these choices being informed by previously proposed models of sensory evidence integration and movement generation (see main text). All six models were described by a 2-dimensional latent ($\mathbf{x}$) *and* observed state ($\mathbf{y}$; observation matrix was set to the identity matrix i.e $\mathbf{C} = I$) as defined below:

$$\dot{\mathbf{x}} = \mathbf{F}(\mathbf{x}) + \mathbf{u}_t + \boldsymbol{\epsilon}_{\mathbf{t}} \qquad (38)$$

$$\mathbf{y}_t = \mathbf{x}_t + \boldsymbol{\eta}_t$$

where, the latent dynamics was governed by a process similar to Equation 1 while, the observation process was linear-gaussian characterized by an observation noise term $\boldsymbol{\eta}_t$. For each of the six model types, we simulated a total of 4000 trials that belonged to one of two conditions ("choice-1" or "choice-2") for a duration of 1s with an integration time-step of 1ms. We estimated the time-varying residual dynamics matrices $\mathbf{A}_t$ by deploying the 2SLS regression (Steps 3 and 4 of analysis pipeline Extended Data Fig. 2) directly on the 2D residuals (i.e without Steps 1 and 2 of the analysis pipeline, see Extended Data Fig. 2) that were computed by removing the condition average trajectories for the two choices from their corresponding 2D single trial trajectories. Additionally, we did not optimize any of the hyper-parameters involved in the dynamics estimation. We used dimensionality (d) = 2, lag (l) = 5 and a regularization parameter ($\alpha$) = 100 for fitting the dynamics matrices for all models. We describe the specific details of all the six models below:

*Saddle node (Unstable Integration)*

This model implemented binary sensory decisions through an instability (saddle node), described by a non-linear dynamical process. The latent state ($\mathbf{x}$) of the model was governed by the following equation:

$$\mathbf{F}(\mathbf{x}) = \begin{bmatrix} -6x_{(1)}^3 + 20x_{(1)} \\ -10x_{(2)} \end{bmatrix} \tag{39}$$

where, $x_{(i)}$ is the $i^{th}$ dimension of the 2-dimensional latent state. $\mathbf{F}(.)$ was characterized by 3 distinct fixed points - an unstable fixed point (saddle node) at the origin: (0,0) where the dynamics are transiently stable and, two stable fixed points located symmetrically at (1.825,0) and (-1.825,0), where the dynamics are globally stable. We sampled initial conditions from a zero-mean bivariate Gaussian distribution with covariance equal to a scaled identity matrix ($= \sigma_i^2.\mathbf{I}$), with $\sigma_i^2$ = 0.005. On each trial 'k' and at each time 't', we chose the inputs driving the model as shown below:

$$\mathbf{u}_t(k) = c(k).\begin{bmatrix} 8 \\ 8 \end{bmatrix} \tag{40}$$

where, $c(k) \in \{-1,1\}$ was a sample from a binomial distribution that determined the sign of the input on each trial. Hence, the deterministic component of the input for this simulation was condition dependent but *time-invariant* i.e. its magnitude remained fixed across trials but its sign varied randomly from trial to trial. Changing the sign of the input from trial to trial thus had the desired effect of the simulated single trial responses belonging to one of two different conditions corresponding to the two different choices ("choice-1" trials: $\mathrm{c(k)}$ = +1, "choice-2" trials: $\mathrm{c(k)}$ = -1). The covariance of the random zero-mean gaussian disturbance ($\boldsymbol{\epsilon}_\mathbf{t}$) was also set to a scaled identity matrix ($= \sigma_e^2.\mathbf{I}$), with $\sigma_e^2$ = 0.0001. The variance of the observation noise ($\boldsymbol{\eta}_t$) was also set to a scaled identity matrix ($= \sigma_n^2.\mathbf{I}$), with $\sigma_n^2$ = 0.000001.

*Line attractor (Perfect Integration)*

We also simulated a model of decision in which evidence integration took place along a line attractor. The line attractor model was implemented using a linear dynamical process, which additionally was characterized by non-normal dynamics:

$$\mathbf{F}(\mathbf{x}) \quad = \mathbf{A}.\mathbf{x} = \begin{bmatrix} 0 & -5 \\ 0 & -5 \end{bmatrix} \mathbf{x} \tag{41}$$

The dynamics matrix ($\mathbf{A}$) had two distinct eigenvalues equal to 0 and -5 and their associated eigenvectors were $[1\ 0]^\mathrm{T}$ and $[1/\sqrt{2}\ \ 1/\sqrt{2}]^\mathrm{T}$ respectively, with the former specifying the local direction of the line attractor. The non-normal nature of the dynamics resulted from the non-orthogonality between the two eigenvectors. The deterministic component of the input across trials at each time instant ($\mathbf{u}_\mathrm{t}$) was chosen independently for each condition ("choice-1" or "choice-2") such that the condition average at the next time step had an exact match to the corresponding condition averaged response simulated from the saddle node model. The covariance matrix of the random zero-mean Gaussian disturbance ($\boldsymbol{\epsilon}_\mathbf{t}$) was again set to a scaled identity matrix ($= \sigma_e^2.\mathbf{I}$), with $\sigma_e^2$ = 0.00003. The other two noise parameters ($\sigma_i^2$, $\sigma_n^2$) were the same as in the saddle node model.

In order to illustrate the various hypothesized relationships between the residual dynamics and the condition-averaged trajectories (Fig. 5), we also simulated an "augmented" line attractor model (Fig 5a-b) characterized by two additional latent dimensions (4 latent dimensions in total). These two additional latent dimensions were orthogonal to the first two latent dimensions (as defined above) and were associated with quick, decaying dynamics (eigenvalues: -7.5 and -10) and sinusoidal inputs. The temporal evolution of this 4-dimensional dynamical system was governed by the following dynamics equation:

$$\dot{\mathbf{x}} = \mathbf{F}(\mathbf{x}) + \mathbf{u}_t + \boldsymbol{\epsilon}_\mathbf{t} = \begin{bmatrix} 0 & -5 & 0 & 0 \\ 0 & -5 & 0 & 0 \\ 0 & 0 & -7.5 & 0 \\ 0 & 0 & 0 & -10 \end{bmatrix} \mathbf{x} + \begin{bmatrix} \mathrm{u_1(t)} \\ \mathrm{u_2(t)} \\ \mathrm{u_3(t)} \\ \mathrm{u_4(t)} \end{bmatrix} + \boldsymbol{\epsilon}_\mathbf{t}$$

where, the input components $\mathrm{u_1(t)}$ and $\mathrm{u_2(t)}$ were the same as defined previously for the line attractor model, and components $\mathrm{u_3(t)}$ and $\mathrm{u_4(t)}$ were defined as sinusoidal functions,

$20 \times cos(1.6\pi f t)$ and $20 \times sin(1.6\pi f t)$ respectively. The various noise parameters of the model ($\sigma_i^2$, $\sigma_e^2, \sigma_n^2$) were the same as in the line attractor model.

*Point Attractor with time-varying inputs (Leaky Integration)*

A 'leaky' model of evidence integration was implemented with the following linear dynamics function:

$$\mathbf{F}(\mathbf{x}) \quad = \mathbf{A}.\mathbf{x} = \begin{bmatrix} -20 & 0 \\ 0 & -40 \end{bmatrix} \mathbf{x} \tag{42}$$

The matrix $\mathbf{A}$ in the above equation is normal (orthogonal eigenvectors) and has two distinct strongly stable eigenvalues equal to -20 and -40. The deterministic component of the input ($\mathbf{u}_\mathrm{t}$) for this model was again chosen to ensure an exact match between the resulting condition averages and those corresponding to the saddle node model. The various noise parameters of the model ($\sigma_i^2$, $\sigma_e^2, \sigma_n^2$) were the same as in the line attractor model.

*Rotational Dynamics*

We implemented a model of movement that was characterized by rotational dynamics. We parameterized the dynamics in terms of polar coordinates $[\mathrm{r}\ \phi]$:

$$\begin{aligned} r &= \sqrt{x_{(1)}^2 + x_{(2)}^2} \\ \phi &= tan^{-1}\left(\frac{x_{(2)}}{x_{(1)}}\right) \\ \dot{r} &= 0 \\ \dot{\phi} &= 1.2r \end{aligned} \tag{43}$$

The latent state ($\mathbf{x}_\mathrm{t}$) on each 'trial' was a result of purely autonomous dynamics set by an initial condition that differed across trials. The initial condition on the $\mathrm{k}^{\mathrm{th}}$ trial was sampled from a Gaussian distribution as follows:

$$\mathbf{x}_0^k \sim \mathcal{N}\left(\begin{bmatrix} c(k) \\ 0 \end{bmatrix}, \begin{bmatrix} 0.01 & 0 \\ 0 & 0.0001 \end{bmatrix}\right) \tag{44}$$

where, $\mathrm{c(k)}$ determined the sign of the mean of the random initial condition vector on each trial (similar to $\mathrm{c(k)}$ in the saddle node dynamics model). Thus, differences in initial conditions across trials determined the two types of conditions ("choice-1" trials: = $\mathrm{c(k)}$ = +1, "choice-2" trials: = $\mathrm{c(k)}$ = -1). The latent noise ($\boldsymbol{\epsilon}_\mathrm{t}$) was described by a zero-mean gaussian with covariance set to a scaled identity matrix ($= \sigma_e^2.\mathbf{I}$), with $\sigma_e^2$ = 0.00003. The variance of the observation noise ($\boldsymbol{\eta}_t$) was also set to a scaled identity matrix ($= \sigma_n^2.\mathbf{I}$), with $\sigma_n^2$ = 0.000001.

Similar to the augmented line attractor model, we also simulated an augmented rotational dynamics model with two additional latent dimensions. The recurrent dynamics and the inputs associated with these two additional dimensions were the same as defined previously for the augmented line attractor model.

##### *Dynamic Attractor*

The apparent rotational structure in the condition averages could also result from a dynamical system characterized by entire trajectories that are locally attracting instead of having purely rotational dynamics. This is in contrast to the classical 'steady state' and static notion of an attractor state. We constructed a dynamical system with a stable attracting circular trajectory. We parameterized the dynamics in terms of polar coordinates as follows:

$$\begin{aligned} \rho &= 1 - r \\ \dot{\rho} &= -20\rho \\ \dot{\phi} &= 1.2(1 - \rho) \end{aligned} \tag{45}$$

The initial conditions for all trials were chosen to be the same as the initial conditions generated for the rotational dynamics model. The model was completely autonomous without any external inputs. The latent noise ($\boldsymbol{\epsilon}_{\mathbf{t}}$) was described by a zero-mean gaussian with covariance set to a scaled identity matrix ($= \sigma_e^2.\mathbf{I}$), with $\sigma_e^2$ = 0.0001. The variance of the observation noise ($\boldsymbol{\eta}_t$) was also set to a scaled identity matrix ($= \sigma_n^2.\mathbf{I}$), with $\sigma_n^2$ = 0.000001.

##### *Point Attractor with time-varying inputs (input-driven movement dynamics)*

The dynamics of this model was the same as the dynamics of the leaky integrator model proposed for evidence integration. This model however was non-autonomous (unlike the rotational dynamics and dynamic attractor models). The deterministic component of the driving input for each trial type was chosen independently, such that the resulting condition averages matched to those simulated from the rotational dynamics model. The initial conditions were the same as those generated for the rotational dynamics simulation. The covariance of the noise term ($\boldsymbol{\epsilon}_{\mathbf{t}}$) was set to a diagonal matrix with each entry equal to 0.0003.

#### 10.2: Linear, time-varying state-space models with uncorrelated input noise

We also validated our approach on simulated data from six distinct latent-variable state space models (Extended Data Fig. 3), which were characterized by (i) three distinct linear but time-varying latent dynamics processes, and (ii) two distinct observation processes, either linear-gaussian or poisson. In addition to these six models, we also simulated three more models with linear-gaussian observations but, which were subject to both time-varying dynamics *and* time-varying latent noise. These simulations demonstrated the robustness of our estimation procedure to different latent dynamics and observation model types. For all nine models, the residual dynamics were estimated using neural residuals binned in 45ms time bins that were subjected to the full analysis pipeline (Extended Data Fig. 2, without session alignment) with cross-validation of the hyper-parameters.

##### *Linear-Gaussian Models*

Models with a linear-gaussian observation process followed the same functional form as Equation 36, with the exception that the dynamics matrix varied as a function of time ($\mathbf{A}_\mathrm{t}$ instead of $\mathbf{A}$ in Equation 36). The models were defined by 3 latent dimensions and 20 observed dimensions.

For linear-gaussian models characterized by *time-invariant latent noise*, the latent noise covariance was chosen to be a scaled identity matrix ($\mathbf{Q} = 0.5\mathbf{I}$). The observation noise matrix $\mathbf{R}$ was set to be diagonal with the diagonal elements sampled from a uniform distribution between $[0{,}0.01]$. The steady-state initial noise covariance ($\mathbf{Q}_0$) was obtained by solving the continuous time Lyapunov equation. The above choice of $\mathbf{Q}$, $\mathbf{R}$ and $\mathbf{Q}_0$ resulted in simulated data with variability that matched the level of variability in the neural data (Extended Data Fig. 3e).

The other simulation parameters were the same as those used for the simulations of the time-invariant LDS model used to determine the optimal bin-size (see Sec. 9.1-Estimating bias in the estimates of residual dynamics). The time-varying dynamics matrices underlying the three, distinct latent dynamical processes are described below:

1. *Switching eigenvalues (non-rotational)*

   The time-varying dynamics matrices of this model were normal (orthogonal eigenvectors) but showed an abrupt switch in the eigenvalue spectrum (Extended Data. Fig. 3a-c, top row). The eigenvectors of the ($\mathbf{A}_\mathrm{t}$) were invariant in time. The elements of the eigenvector matrix were sampled from a standard normal distribution and then orthogonalized with respect to one another. The eigenvalue spectrum was always real and switched from $[-1, -3, -5]$ to $[-2, -4, -6]$ midway through the trial.

2. *Switching eigenvectors*

   The dynamics matrices of this model had eigenvalues $[-1, -3, -5]$ which did not vary in time. Instead the associated eigenvector basis exhibited a switch from an orthogonal ('normal' dynamics) to a non-orthogonal eigenvector basis ('non-normal' dynamics) mid-way through the trial (Extended Data. Fig. 3a-c, middle row).

3. *Switching eigenvalues (rotational)*

The time-varying dynamics matrices of this model (Extended Data. Fig. 3a-c, bottom row) were normal (orthogonal eigenvectors) but showed an abrupt switch in the eigenvalue spectrum from real - $[-1, -3, -5]$ to complex-valued eigenvalues - $[-2 + \mathrm{i}2\pi, -2 - \mathrm{i}2\pi, -5]$ mid-way through the trial. These complex eigenvalues were associated with 'slow' rotations of 1Hz frequency.

Linear-gaussian models with *time-varying* latent noise differed only in a single aspect. They were characterized by latent noise covariance that was itself time-dependent ($\mathbf{Q}_t$ instead of $\mathbf{Q}$). The latent noise covariance switched from $0.5\mathbf{I}$ to $0.3\mathbf{I}$ midway through the trial, locked to the corresponding change in the latent dynamics ($\mathbf{A}_\mathrm{t}$).

##### *Poisson Models*

Models described by a poisson observation process shared the same time-varying, latent dynamics processes as the models with the gaussian observations, but were associated with 100 observed dimensions. The poisson observations ($\mathbf{s}_t$), which corresponded to spike counts in 1ms bins were described as follows:

$$\begin{aligned} \mathbf{y}_t &= \mathbf{C}\mathbf{x}_t + \mathbf{d} \\ \mathbf{s}_t &\sim Poisson(exp(\mathbf{y}_t)) \end{aligned} \tag{46}$$

where, $\mathbf{x}_t$ is the latent state defined by the latent dynamical process. The observation matrix $\mathbf{C}$ was the same as in the linear-gaussian model simulations. The elements of the baseline input vector $\mathbf{d}$ were chosen uniformly at random in the range [-4.8 -4.1], while the latent noise covariance was time-invariant and set to a scaled identity matrix - $\mathbf{Q} = 12500\mathbf{I}$. The above choice of $\mathbf{Q}$ and $\mathbf{d}$ ensured realistic, over-dispersed fano factors between 1 and 1.5, and single-trial variability that matched the level of variability in the neural data (Extended Data Fig. 3e). The steady-state initial noise covariance ($\mathbf{Q}_0$) was once again obtained by solving the continuous time lyapunov equation. For the poisson models, the neural residuals were computed after performing a square-root transformation of spike counts, counted in 45ms long time bins.

To match the overall level of variability in the simulated residuals to those measured in the data, we computed three different measures that specified (i) the total magnitude of instantaneous variability ($\boldsymbol{l}_0 = \langle \|\tilde{\mathbf{Z}}_\mathrm{t}\tilde{\mathbf{Z}}_\mathrm{t}{}'\|_\mathrm{F}^2 \rangle_\mathrm{t}$), (ii) the total magnitude of lagged co-variability ($\boldsymbol{l}_1 = \langle \|\tilde{\mathbf{Z}}_{\mathrm{t}+1}\tilde{\mathbf{Z}}_\mathrm{t}{}'\|_\mathrm{F}^2 \rangle_\mathrm{t}$) and, (iii) the strength of the most dominant mode of lagged co-variability ($\mathbf{pvar} = \langle \sigma_{max}(\tilde{\mathbf{Z}}_{\mathrm{t}+1}\tilde{\mathbf{Z}}_\mathrm{t}{}') \rangle_\mathrm{t}$) in the simulated and measured residuals, where, $\tilde{\mathbf{Z}}_\mathrm{t}$ is the aligned residual data matrix at time ‘t’ of dimensionality $20 \times \mathrm{K}$ (=number of trials), $\| . \|_{\mathrm{F}}$ is the frobenius norm, and $\sigma_{max}(.)$ corresponds to the largest singular value of its matrix-valued argument. The latent noise parameter ($\mathbf{Q}$) of the linear gaussian/poisson models, the observation noise parameter ($\mathbf{R}$) of the linear gaussian models, and the mean offset ($\mathbf{d}$) of the poisson model simulations were chosen such that the measures of variability (as defined above) qualitatively matched to those computed using PFC neural residuals.

#### 10.3: Linear time-varying state-space models with colored input noise

To study the inflationary effects of correlated input variability on estimates of residual dynamics in the complex input regime (Fig 1b), we simulated state-space models with linear time-invariant latent dynamical processes characterized by latent noise with decaying temporal autocorrelations (colored noise). These models were a precursor to more complex modular RNN models characterized by distributed recurrence, and allowed us to understand how neural activity that is a consequence of recurrence in unobserved/unrecorded areas, influences residual dynamics measured within recorded/observed areas. We assumed a time-invariant, linear state-space model governed by the following equations.

$$\begin{aligned} \mathbf{x}(\mathrm{t}+1) &= \mathbf{A}\mathbf{x}(\mathrm{t}) + \boldsymbol{\varepsilon}(\mathrm{t}) \\ \boldsymbol{\varepsilon}(\mathrm{t}+1) &= \boldsymbol{\phi}\, \boldsymbol{\varepsilon}(\mathrm{t}) + \boldsymbol{\zeta}(\mathrm{t}) \\ \mathbf{z}(\mathrm{t}) &= \mathbf{C}\mathbf{x}(\mathrm{t}) + \boldsymbol{\eta}(\mathrm{t}) \end{aligned} \tag{47}$$

where, $\boldsymbol{\zeta}(t)$ and $\boldsymbol{\eta}(t)$ are assumed to be zero-mean white gaussian noise processes with covariance matrices $\mathbf{Q}$ and $\mathbf{R}$ respectively. A convenient interpretation of the above model assumes that $\mathbf{x}(\mathrm{t})$ is the latent population state in the recorded/observed area, yielding observations $\mathbf{z}(\mathrm{t})$. On the other hand, $\boldsymbol{\varepsilon}(\mathrm{t})$, which is equivalent to an auto-correlated input noise process, can be considered to be the latent population state of unobserved or unrecorded areas, that directly influence $\mathbf{x}(\mathrm{t})$. Therefore, $\boldsymbol{\phi}$ encapsulated the dynamics of the unobserved/unrecorded areas, and its eigenvalues determined the time-scale of the autocorrelation of $\boldsymbol{\varepsilon}(\mathrm{t})$. For a given $\mathbf{A}$, $\boldsymbol{\phi}$ and $\mathbf{Q}$, we were able to analytically derive the two-stage least squares (2SLS) estimate of the residual dynamics (see Supplementary Math Note B), assuming that (i) the model operated in *steady-state* and, (ii) that we only had access to $\mathbf{x}(\mathrm{t})$ through observations $\mathbf{z}(\mathrm{t})$.

##### Steady-state analyses

We systematically varied $\mathbf{A}$, $\boldsymbol{\phi}$ in the model of Equation 47 to quantify the effect of correlated input noise on analytically derived estimates of residual dynamics, under steady-state conditions (Extended Data. Fig. 4,5). To simplify our analyses, we only considered a system with 2-

dimensional dynamics (i.e. $\mathrm{Dim}(\mathbf{x}) = \mathrm{Dim}(\boldsymbol{\varepsilon}) = 2$), although our analytical derivations were applicable for arbitrary dimensonalities. As discussed in the points below, we explored three distinct categories of models, all specified by Equation 47, but which differed in the nature of $\mathbf{A}$ and $\boldsymbol{\phi}$ (i.e. either stable or rotational). Within each category, we generated multiple instantiations of the model by systematically sweeping the magnitude and phase of the eigenvalues of $\mathbf{A}$ and $\boldsymbol{\phi}$. For all model instantiations across all three categories, the covariance of the latent noise process $\boldsymbol{\zeta}(\mathrm{t})$ was set to a scaled identity matrix ($\mathbf{Q} = \sigma_q^2 . \mathbf{I}$), with $\sigma_q^2$ sampled uniformly at random in the range [$10^{-12}$, $10^{-10}$], separately for each model instantiation. Similarly, the covariance of the observation noise process $\boldsymbol{\eta}(\mathrm{t})$ was set to a scaled identity matrix ($\mathbf{R} = \sigma_r^2 . \mathbf{I}$), with $\sigma_r^2$ sampled uniformly at random in the range [$10^{-4}$, $10^{-2}$], separately for each model.

1. *Inflation of eigenvalues (Extended Data Fig. 4a-b)*

   We instantiated a total of 5000 distinct models within this category by varying the eigenvalue magnitudes of $\mathbf{A}$ and $\boldsymbol{\phi}$. For all 5000 models, we set one of the eigenvalues of $\mathbf{A}$ to a fixed constant such that it was associated with a decay time constant of 400ms. We swept the other eigenvalue ($\lambda_2$, Extended Data. Fig. 4a-b) on a uniform grid in the range [0.4 0.999], resulting in 1000 distinct $\mathbf{A}$ matrices. Both eigenvalues of $\boldsymbol{\phi}$ were constrained to be equal to one another ($\lambda_1$, Extended Data. Fig. 4a-b) and were sampled in 5 discrete steps ($\lambda_1 = [0.40\ 0.75\ 0.91\ 0.97\ 0.99]$), which resulted in 5 distinct $\boldsymbol{\phi}$ matrices. The eigenvectors of each $\mathbf{A}$ and $\boldsymbol{\phi}$ were the cardinal unit vectors $\mathbf{e_1}$, $\mathbf{e_2}$ of the 2D space. We also simulated 5 additional models, in which, $\lambda_2$ (eigenvalue specifying $\mathbf{A}$) was set to zero specifying a scenario, in which, recurrent dynamics only unfolds upstream within the input area (Extended Data Fig. 4b).

2. *Inflation of rotation frequency (Extended Data Fig. 4c-d)*

   We instantiated a total of 5000 distinct models within this category by varying the eigenvalue phase of $\boldsymbol{\phi}$ and the eigenvalue magnitude of $\mathbf{A}$ (Extended Data. Fig. 4c-d). Since $\boldsymbol{\phi}$ was associated with rotational dynamics, it was associated with a pair of complex-conjugate eigenvalues, which could be completely specified using its magnitude and phase. For all 5000 models, we set the magnitude of the complex-conjugate eigenvalue pair to a fixed constant that translated into a decay time constant of 1s. We instead systematically varied the phase of the complex-conjugate eigenvalue pair in 5 discrete steps, resulting in 5 distinct $\boldsymbol{\phi}$ matrices that had associated rotation frequencies ($\omega_1$ , Extended Data. Fig. 4d) equal to [0.5 0.75 1.00 1.25 1.5] Hz. Both eigenvalues of $\mathbf{A}$ were constrained to be equal to one another ($\lambda_2$, Extended Data. Fig. 4c-d) and sampled from a uniform grid in the range [0.4 0.999], resulting in 1000 distinct $\mathbf{A}$ matrices. The eigenvectors of $\mathbf{A}$ were the cardinal unit vectors

$\mathbf{e_1}, \mathbf{e_2}$ of the 2D space. The eigenvectors of $\boldsymbol{\phi}$ were complex-conjugate and were chosen such that $\boldsymbol{\phi}$ was characterized by a circularly symmetric rotational flow-field.

3. *Equivalence of upstream and local recurrent dynamics (Extended Data Fig. 4e-f)*

   We instantiated a total of 5000 distinct models within this category by varying the eigenvalue phase of $\mathbf{A}$ and the eigenvalue magnitude of $\boldsymbol{\phi}$, in a manner that was mirror symmetric to the way the parameters were specified above.

In addition to the above steady-state analyses, we also simulated data from a model that was once again specified by Equation 47, but with a 1-dimensional latent state (i.e. Dim$(\mathbf{x})$ = Dim$(\boldsymbol{\varepsilon})$ = 1), in order to provide a graphical explanation for the inflation of the estimates of residual dynamics. We simulated 1000 trials in steps of 45ms starting from a base scenario (Extended Data Fig. 5a-b) where the temporal latent process of $\mathbf{x}$ was driven by input noise that was not temporally autocorrelated (i.e $\boldsymbol{\phi}$ = 0) and with a fixed variance (Var ($\boldsymbol{\zeta}(\mathrm{t})$) = 1e-6). We then either switched (t=0 is denoted as time of switch in Extended Data Fig 5) to a condition (i) where the input noise ($\boldsymbol{\varepsilon}$) was autocorrelated with $\boldsymbol{\phi}$ = 0.798 (Extended Data Fig. 5c-d, or (ii) where the variance was increase by a factor of 10 to 1e-5 (Extended Data Fig. 5e-f).

#### 10.4 Modular two-area recurrent neural network model

##### Simulation details

We simulated a previously proposed modular RNN model for perceptual decision making, which emulated the distributed fronto-parietal decision network[23]. The model consisted of two individual recurrent modules or "areas" named PPC and PFC. Each area was characterized by a 2-dimensional state variable that encapsulated the activations of two neural populations that were selective for either choice (choice 1 or choice 2). The four neural populations across both areas were connected to one another using excitatory/inhibitory (E/I) intra-areal (within area) recurrent connections, and inter-areal (between areas) E/I feedforward and feedback connections. The intra-areal recurrent connectivity was determined by weight matrices $\mathbf{J}^{ppc}$ and $\mathbf{J}^{pfc}$, whereas the inter-areal connectivity was determined by feed-forward weight matrix $\mathbf{J}^{ppc\rightarrow pfc}$ and feedback weight matrix $\mathbf{J}^{ppc\leftarrow pfc}$. Each of the above four weight matrices were directly determined by 4 scalar parameters $\mathrm{J}_s{}^{ppc}$, $\mathrm{J}_s{}^{pfc}$, $\mathrm{J}_s{}^{ppc\rightarrow pfc}$ and $\mathrm{J}_s{}^{ppc\leftarrow pfc}$, which modulated the coupling strength of the connectivity matrices[23]. We constrained all our analyses to a setting where the within area recurrent weights of each area were the same (i.e $\mathbf{J}^{ppc} = \mathbf{J}^{pfc}$, by virtue of setting $\mathrm{J}_s{}^{ppc}$ = $\mathrm{J}_s{}^{pfc}$ = $\mathrm{J}_{self}$). We simulated two different classes of networks, one in which feedback was absent (i.e $\mathbf{J}^{ppc\leftarrow pfc} = \mathbf{0}$, by virtue of setting $\mathrm{J}_s{}^{ppc\leftarrow pfc}$ = 0) and, another in which feedback was present and equaled the feed-forward connection strength (i.e $\mathbf{J}^{ppc\rightarrow pfc} = \mathbf{J}^{ppc\leftarrow pfc}$, by virtue of setting $\mathrm{J}_s{}^{ppc\rightarrow pfc}$ = $\mathrm{J}_s{}^{ppc\leftarrow pfc}$ = $\mathrm{J}_{across}$). We refer the reader to the original study for specific details about the network architecture, dynamics, and the relationship between the various scalar parameters (defined above) and the corresponding weight matrices[23].

Thirty different network configurations with distinct intra-areal and inter-areal connectivity were simulated by sweeping the two scalar parameter values $\mathrm{J}_{self}$ (5 distinct values, colored markers, Fig. 6c,f) and $\mathrm{J}_{across}$ (6 distinct values, x-axis Fig. 6c,f) for each of the two network classes (i.e with and without feedback connections). We simulated 6000 trials from each network, each trial spanning a total of 1.2s, using a simple Euler integration scheme in steps of 0.1ms. Only PPC was driven using external current input that was defined as follows:

$$I^k(t) = \begin{cases} 0\,, \quad 0 < t \le T_{on} \\ I_e\left(1 \pm \dfrac{c(k)}{100\%}\right), \qquad t > T_{on} \end{cases} \tag{48}$$

where, $I_e$ = 0.0130nA, $T_{on}$ (= 400ms) is the time of stimulus onset and, *c(k)* corresponds to the coherency on the $k^{\text{th}}$ trial. We simulated only trials with zero coherency (c(k) = 0) and assigned each trial as either "choice-1" or "choice-2", depending on the population choice readout from "PFC" at the last time step of the trial. Each population within an area also received noisy current inputs determined by an Ornstein-Uhlenbeck (OU) process whose variance $\sigma_{noise}$ was set to 0.07, so as to match the level of signal to noise in the recorded neural data.

We denote the simulated activations of the network at time 't' from area '*a*' as $\mathbf{x}_t^a$, which constitutes a 2-dimensional latent state (one dimension per choice selective population). We projected this 2-dimensional latent state linearly into a 10-dimensional observation space to obtain area-specific observations $\mathbf{y}_t^a$ as described below:

$$\mathbf{y}_t = \begin{pmatrix} \mathbf{y}_t^{ppc} \\ \mathbf{y}_t^{pfc} \end{pmatrix} = \mathbf{C}_{\text{model}}\mathbf{x}_{\text{t}} + \boldsymbol{\eta}_{\text{t}} = \begin{bmatrix} \mathbf{C}_{\text{ppc}} & \mathbf{0} \\ \mathbf{0} & \mathbf{C}_{\text{pfc}} \end{bmatrix} \begin{pmatrix} \mathbf{x}_t^{ppc} \\ \mathbf{x}_t^{pfc} \end{pmatrix} + \boldsymbol{\eta}_{\text{t}} \tag{49}$$

where, $\mathbf{C}_{\text{ppc}}, \mathbf{C}_{\text{pfc}} \in \mathcal{R}^{10\times 2}$ are random orthogonal matrices that form the diagonal entries of a block diagonal matrix $\mathbf{C}_{\text{model}} \in \mathcal{R}^{20\times 4}$ and, $\boldsymbol{\eta}_{\text{t}} \in \mathcal{R}^{20\times 1}$ was an observation noise vector drawn from a multi-variate gaussian with isotropic variance equal to 0.0006. Therefore, the global observation state $\mathbf{y}_t$, used to define residuals across both areas of the model consisted of observations from a total of 20 observation dimensions, 10 each assigned to PPC and PFC. For the network configuration shown in Fig. 6a ($\mathrm{J}_{self}$ = 0.36, $\mathrm{J}_{across} = 0.08$, no feedback), we simulated an identical network (with frozen noise) for a 'shuffled' condition, in which, only the feed-forward current inputs at each time from PPC to PFC were randomly shuffled across trials, in order to remove any slow temporal autocorrelations.

#### Residual dynamics estimation

We estimated residual dynamics either 'locally' (see Fig. 6b), using observations of PPC alone ($\mathbf{y}_t^{ppc}$) or PFC alone ($\mathbf{y}_t^{pfc}$); or 'globally' (Fig. 7a), using observations from both areas ($\mathbf{y}_t$). Observations were temporally binned in 45ms long bins (as in the neural data) and residual dynamics was computed separately for trials belonging to each of the two choice conditions by employing the full analysis pipeline, but excluding the session alignment step (Step 1 in Extended Data Fig. 2).

For local estimates of residual dynamics, the cross-validation of the hyper-parameters (see Step 3 of Extended Data Fig. 2) yielded dynamics of dimensionality 'd' = 2 for all network configurations, which was consistent with the ground truth. For the global estimates, depending on the underlying network configuration, we recovered residual dynamics with dimensionalities d in the range 2-4. As in the neural data (Extended Data Fig. 8), we consistently recovered an optimal lag (l) of either 3 or 4 across most model configurations. Therefore, for the final model fits (Figs. 6,7), we used a fixed lag 'l' = 3 for all model configurations. The smoothness hyper-parameter ($\alpha$ in Equation 30) of the second stage of the two-stage least squares procedure (see Step 4 of Extended Data Fig. 2) was uniquely determined for each model using the cross-validation procedure described previously.

Additionally, we also computed the *local choice* residual dynamics, which was defined as the dynamics obtained by fitting the 1-dimensional projection of the local residuals of a given area along the choice dimension specific to that area (see Equation 50 for definition of choice dimension). Furthermore, we also examined the relationship between the 1D *local choice* residual dynamics and the network connectivity parameters $\mathrm{J}_{self}$ and $\mathrm{J}_{across}$. We measured the largest eigenvalue magnitude (across time in the trial) along the choice dimension local to each area and plotted it as a function of $\mathrm{J}_{self}$(colors in Fig. 6c,f) and $\mathrm{J}_{across}$ (x-axis in Fig. 6c,f). We derived confidence intervals for the largest eigenvalue magnitude along the choice dimension (error bars in Fig. 6c,f) by bootstrapping the 1D fits of the residual dynamics by repeated resampling (with replacement) of the simulated trials.

#### Task-relevant subspaces and its overlap with residual dynamics

For each network configuration, we computed two distinct "task-relevant" dimensions, similar in spirit to the choice and time subspaces computed using trial-averaged neural responses in the real neural data (Fig. 3c-d). These task-relevant dimensions were *area-specific*, as in, they were computed separately for each area of the two-area RNN. Given that each area of the model is characterized by two populations (selective for either choice 1 or choice 2), implying a two-dimensional latent state-space per area, the direction $[1\ -1]^T$ of the latent space is naturally defined as the "choice" axis and the direction $[1\ 1]^T$ as the "time" axis. Therefore, in total we obtained 4 latent "task-relevant" dimensions within a 4-dimensional latent-space of the model (2 latent dimensions per area) as shown below:

$$\begin{aligned} \boldsymbol{u}_{choice}^{ppc} &= [1\ \ -1\ \ 0\ \ 0]^T \\ \boldsymbol{u}_{time}^{ppc} &= [1\ \ 1\ \ 0\ \ 0]^T \\ \boldsymbol{u}_{choice}^{pfc} &= [0\ \ 0\ \ 1\ -1]^T \\ \boldsymbol{u}_{choice}^{pfc} &= [0\ \ 0\ \ 1\ -1]^T \end{aligned} \qquad (50)$$

where, the first 2 dimensions of the 4D latent space correspond to PPC and the last two dimensions correspond to PFC. Therefore, activity modulations along the choice and time model axes as defined above gave rise to condition-dependent and condition-independent variance respectively as seen in the simulated neural activity within each area (Fig. 6a,d).

To compute the overlap of these task-relevant dimensions with the local and global estimates of residual dynamics, for each model configuration (pair of $\mathrm{J}_{self}$ and $\mathrm{J}_{across}$), we extracted the corresponding estimated residual eigenvectors at a specific time instant $t_{max}^{pfc}$, which was defined as the time instant at which the maximum eigenvalue magnitude was attained 'locally' within PFC ($t_{max}^{pfc} = \operatorname{argmax}_t \max_j |\lambda_j^{pfc}(t)|$ , where *'j'* indexes the *j*th eigen-mode). The overlap between residual eigenvectors and the task-relevant model dimensions was quantified as the subspace angle between pairs of vectors. In order to use vectors of a consistent dimensionality, we represented all the relevant vectors/dimensions in the 20-dimensional space of observations. We achieved this by projecting the 4-dimensional task-relevant dimensions (defined in Equation 50) through the ground-truth model observation matrix $\mathbf{C}_{\mathrm{model}}$, and the estimated 'local' and 'global' residual eigenvectors through the corresponding estimate of $\mathbf{C}_{\mathrm{model}}$ (i.e dynamics subspace) derived through our fitting procedure.

#### Simulated perturbation experiments

The modular RNN model also provided an ideal framework to demonstrate how residual dynamics can be combined with causal perturbations to reveal distributed computations in in neural circuits. We performed a set of targeted causal perturbation experiments (Fig. 8) in the two example network configurations (Fig. 6 & 7) that differed in particular in terms of the presence of long-range feedback connections from PFC to PPC. We first obtained a set of "ground truths" that summarized how activity patterns associated with each area change in response to an injected perturbation. We then attempted to predict the perturbed activity patterns using either the ‘local’ or the ‘global’ estimates of residual dynamics.

Perturbations corresponded to small pulse-like current injections within a specific area along one of the two distinct task-relevant, model directions as defined previously, i.e. along $\boldsymbol{u}_{choice}^{ppc}$, $\boldsymbol{u}_{time}^{ppc}$, $\boldsymbol{u}_{choice}^{pfc}$, or $\boldsymbol{u}_{time}^{pfc}$. The magnitude of each perturbation was set to a fixed constant of 0.001nA, which was chosen to approximately match the extent of single-trial neural variability in the absence of a perturbative input. As a consequence, the perturbations were restricted to the “intrinsic manifold” of naturally occurring activity patterns of the model. Furthermore, the pulse-like perturbations were very short (0.1ms length), and restricted to six different times in the trial ([270 405 540 675 810 945]ms after trial onset). For each network configuration, we simulated 6000 unperturbed trials, in which the model was simulated using the same control conditions as described previously (see Simulation details). Additionally, we also simulated 6000 perturbed trials for each of the 24 distinct perturbation conditions (4 directions across 2 nodes *x* 6 times). Each perturbed trial was paired with a corresponding unperturbed trial, with which it shared the time course of the latent noise and input noise.

To compute the ground truth impulse response for a specific perturbation condition, we sorted the “unperturbed” trials into two choice conditions (as before, assignment of choice was based on thresholding the single-trial readout along $\boldsymbol{u}_{choice}^{pfc}$ at the last time instant), and computed trial-averaged trajectories specific to each area, for each choice condition (choice-1 and choice-2). Likewise, we sorted the perturbed trials in each area based on the assignment of choice determined for the corresponding unperturbed trials, and computed “perturbed residuals” by subtracting from each perturbed single trial the corresponding unperturbed, choice-specific trial-averaged trajectory. The ground truth impulse response in a given area was then defined as the mean across trials of the ‘perturbed residuals’ specific to that area.

To construct the impulse response $\mathbf{v}_{prop}(t')$ predicted by residual dynamics, the estimated time-varying dynamics matrices ($\mathbf{A}_{\mathrm{t}}$) were combined as follows:

$$\mathbf{v}_{prop}(t') = \mathbf{U}_{dyn}^{d}.(\mathbf{A}_{t'}.\mathbf{A}_{t'-\Delta t}. \dots \mathbf{A}_{t_0+\Delta t}).\mathbf{U}_{dyn}^{d}{}'\mathbf{v} \qquad (51)$$

where, $\Delta t$ corresponds to the temporal bin size, $t_0$ corresponds to the time at which the perturbation is applied, $\mathbf{U}_{dyn}^{d}$ is the estimate of the dynamics subspace and, $\mathbf{v}$ corresponds to the initial condition. We estimated residual dynamics using only the ‘unperturbed’ trials, separately for each choice condition. We constructed two types of predicted impulse responses – (i) using local residual dynamics estimates constructed out of partial observations only from a single area (i.e either $\mathbf{y}_t^{ppc}$ or $\mathbf{y}_t^{pfc}$) and, (ii) using global residual dynamics estimates that were derived using observations from both areas ($\mathbf{y}_t$). The initial condition $\mathbf{v}$ was set to the ground truth impulse response computed at the exact time instant of that perturbation. In other words, here we did not attempt to predict $\mathbf{v}$ itself (i.e. activity during the current injection), but rather how activity in the simulated networks evolved from $\mathbf{v}$ after the end of the current injection.

To summarize the ground-truth and predicted impulse responses, we computed the norm of the impulse response and plotted it as a function of time in the trial (Fig. 8, colored points: ground truth, black/gray curves: predictions). We only reported the two-step look-ahead impulse responses, i.e we only plotted the impulse response at the time-instant of a given perturbation (which corresponds to $\mathbf{v}$ for both the prediction and the ground-truth) and the two time instants that immediately followed the perturbation ($t' = t_0 + 2\Delta t$, in Equation 51). The impulse response predicted using global dynamics (recordings from entire circuit) explained more variance in the ground-truth impulse response ($R^2 = 0.968$) than the impulse response predictions based on local dynamics (partial recordings from single area, $R^2 = 0.928$).

To better understand the properties of the simulated and predicted impulse responses, we also simulated impulse responses in simpler, two-dimensional linear time-invariant dynamical systems meant to reproduce key features of the global dynamics in the two example networks (Fig. 7d). The two cardinal directions of the linear dynamical systems corresponded to the choice modes in PPC and PFC, and perturbations were applied along one or the other cardinal direction, mimicking perturbations local to the corresponding areas. The two models only differed in the arrangement of the two eigenvectors, but not in the magnitudes of the two associated EV. One EV was unstable, the other stable (magnitudes of 1.1 and 0.5, respectively). For the feedforward model, the unstable eigenvector ($EV_1$, see Fig. 7a, top) projected mostly onto the PPC choice mode (angle with $PPC_c$ = 15 deg, Fig. 7d, top), while the stable eigenvector ($EV_3$, see Fig. 7a, top) was perfectly aligned with the PFC choice mode (angle with $PFC_c$ = 0, Fig 7d, top). For the "feedback" model, both the unstable ($EV_1$, see Fig. 7a, bottom) and stable ($EV_4$, see Fig. 7a, bottom eigenvectors were equally shared across PPC and PFC (angle of 45 deg with $PPC_c$ and $PFC_c$; Fig 7d, bottom).

### II. Supplementary Math Notes

#### A. Background on subspace identification theory

In this section, we provide a brief overview of subspace system identification (SSID) theory as applied to parameter identification of linear time-invariant (LTI) state-space models. We adapt linear, time-invariant SSID methods described below to models characterized by a linear time-varying latent dynamical process, such as the one used in our study to investigate residual dynamics (Equation 11, see Methods). A standard linear, time-invariant state-space model can be described as:

$$\begin{aligned} \mathbf{x}(\mathrm{t}+1) &= \mathbf{A}\mathbf{x}(\mathrm{t}) + \boldsymbol{\varepsilon}(\mathrm{t}) \\ \tilde{\mathbf{z}}(\mathrm{t}) &= \mathbf{C}\mathbf{x}(\mathrm{t}) + \boldsymbol{\eta}(\mathrm{t}) \end{aligned} \tag{52}$$

where, $\mathbf{A}$ and $\mathbf{C}$ are the dynamics and observation matrices respectively, $\mathbf{x}(\mathrm{t})$ is a $p \times 1$ latent state at time ‘t’, $\tilde{\mathbf{z}}(\mathrm{t})$is a corresponding $n \times 1$ observation vector, and $\boldsymbol{\varepsilon}(\mathrm{t})$ and $\boldsymbol{\eta}(\mathrm{t})$ are zero-mean white gaussian noise processes. In our study, “observations” $\tilde{\mathbf{z}}(\mathrm{t})$ correspond to aligned neural residuals, computed by subtracting the condition-averaged trajectory from each corresponding aligned single-trial trajectory (see Methods).

A common approach for parameter estimation of models such as those described by Equation 52 involves probabilistic inference techniques. The expectation-maximization (EM) algorithm provides closed-form updates for the parameter estimates of an LTI model through the Kalman filtering and smoothing recursions[24]. However, estimating system parameters for time-varying dynamics, as in our model (Equation 11, see Methods), is intractable using the exact EM. Such models typically require approximate inference techniques[6,7] (e.g. variational inference) that rely on gradient optimization, which are susceptible to local minima in parameter space.

We therefore opted for an alternative approach based on subspace identification techniques[8,9]. These non-probabilistic methods rely on a series of matrix decompositions and linear regressions to estimate the system parameters in closed form. SSID methods are ‘non-optimal’ in a probabilistic sense, in that, they do not explicitly model the uncertainty in the underlying latent state. However, these methods do not suffer from problems of local minima and can yield unbiased and consistent parameter estimates for large sample sizes.

The key idea underlying SSID methods is to build a linear predictor of the underlying latent state directly from the observed time series $\tilde{\mathbf{z}}$, by matching the empirical first and second order moments to those predicted by the model (Equation 52). This predictor, serves as a “bottleneck”, summarizing all the information in the "past" of the observed time-series that is relevant to predicting the "future" of the time-series. Mathematically, for a given time 't' and trial 'k', we can define the finite "future" and "past" (relative to 't') of the observed time series as follows:

$$\begin{aligned} \tilde{\mathbf{z}}_{\mathbf{t}}^{+}(k) &= [\tilde{\mathbf{z}}_{\mathrm{t}}(k)' \quad \tilde{\mathbf{z}}_{\mathrm{t+1}}(k)' \dots \tilde{\mathbf{z}}_{\mathrm{t+q-1}}(k)']' \\ \tilde{\mathbf{z}}_{\mathbf{t}}^{-}(k) &= [\tilde{\mathbf{z}}_{\mathrm{t-1}}(k)' \quad \tilde{\mathbf{z}}_{\mathrm{t-2}}(k)' \dots \tilde{\mathbf{z}}_{\mathrm{t-q}}(k)']' \end{aligned} \tag{53}$$

where, $\tilde{\mathbf{z}}_{\mathrm{t}}(k)$ is the observation vector of dimensionality $\mathrm{n} \times 1$ in time-bin 't' on trial 'k' that populates the observation data matrix $\tilde{\mathbf{Z}}$ (of dimensionality $\mathrm{n} \times \mathrm{T} \times \mathrm{K}$, where T and K are the total number of time-bins and trials respectively). $\tilde{\mathbf{z}}_{\mathrm{t}}^{+}$ and $\tilde{\mathbf{z}}_{\mathrm{t}}^{-}$ respectively are the 'future' and 'past' observation vectors (each of dimensionality $(\mathrm{n}.\mathrm{q}) \times 1$) constructed by stacking together *q* 'future' and, *q* 'past' lags of the observed time series relative to 't'.

An ordinary least squares linear predictor of the "future" ($\tilde{\mathbf{z}}_{\mathbf{t}}^{+}$) based on the past ($\tilde{\mathbf{z}}_{\mathbf{t}}^{-}$) is defined as:

$$\hat{\tilde{\mathbf{z}}}_{\mathbf{t}}^{+} = \left(\mathrm{Cov}(\tilde{\mathbf{z}}_{\mathrm{t}}^{+}, \tilde{\mathbf{z}}_{\mathrm{t}}^{-})\mathrm{Cov}^{-1}(\tilde{\mathbf{z}}_{\mathrm{t}}^{-}, \tilde{\mathbf{z}}_{\mathrm{t}}^{-})\right)\tilde{\mathbf{z}}_{\mathrm{t}}^{-} = \mathbf{H}_{\mathrm{t}}\mathbf{G}_{\mathrm{t}}{}^{-1}\tilde{\mathbf{z}}_{\mathrm{t}}^{-} \tag{54}$$

The "future-past" covariance matrix $\mathbf{H}_t$ specified by $\mathrm{Cov}(\tilde{\mathbf{z}}_{\mathrm{t}}^{+}, \tilde{\mathbf{z}}_{\mathrm{t}}^{-})$ in Equation 54 is referred to as a "*hankel matrix*", and is a key element of SSID methods. The hankel matrix is in fact a block matrix of size $(\mathrm{n} \times \mathrm{q}) \times (\mathrm{n} \times \mathrm{q})$, with the individual blocks representing temporally lagged covariance matrices computed using observation vectors $\tilde{\mathbf{z}}$ at different time-lags (relative to 't')

$$\mathbf{H}_{\mathrm{t}} = \mathrm{Cov}(\tilde{\mathbf{z}}_{\mathrm{t}}^{+}, \tilde{\mathbf{z}}_{\mathrm{t}}^{-}) = \begin{bmatrix} \tilde{\mathbf{Z}}_{\mathrm{t}}.\tilde{\mathbf{Z}}_{\mathrm{t-1}}' & \tilde{\mathbf{Z}}_{\mathrm{t}}.\tilde{\mathbf{Z}}_{\mathrm{t-2}}' & \dots & \tilde{\mathbf{Z}}_{\mathrm{t}}.\tilde{\mathbf{Z}}_{\mathrm{t-q}}' \\ \tilde{\mathbf{Z}}_{\mathrm{t+1}}.\tilde{\mathbf{Z}}_{\mathrm{t-1}}' & \tilde{\mathbf{Z}}_{\mathbf{t+1}}.\tilde{\mathbf{Z}}_{\mathrm{t-2}}' & \dots & \tilde{\mathbf{Z}}_{\mathrm{t+1}}.\tilde{\mathbf{Z}}_{\mathrm{t-q}}' \\ \vdots & & \ddots & \vdots \\ \tilde{\mathbf{Z}}_{\mathrm{t+q-1}}.\tilde{\mathbf{Z}}_{\mathrm{t-1}}' & \tilde{\mathbf{Z}}_{\mathrm{t+q-1}}.\tilde{\mathbf{Z}}_{\mathrm{t-2}}' & \dots & \tilde{\mathbf{Z}}_{\mathrm{t+q-1}}.\tilde{\mathbf{Z}}_{\mathrm{t-q}}' \end{bmatrix} \tag{55}$$

where, $\tilde{\mathbf{Z}}_{\mathrm{t}}$ is an individual slice of the observation data matrix $\tilde{\mathbf{Z}}$ indexed by time 't'. For LTI systems in steady state, stationarity implies that the "future-past" hankel covariance matrix in Equation 55 is *time independent*, since the individual blocks of the hankel matrix are covariance matrices that only depend on the relative lag between temporal observations (see Equations 55 and 56) under stationary conditions.

The parameter '*q*' (referred to as the order of the hankel matrix) is a critical parameter required for accurate model identification of an LTI model. SSID theory specifies that *q* has to be chosen to be at least as large as the dimensionality of the true underlying latent state. An alternative interpretation of the above statement is that a given choice of $q$ sets the upper bound for the dimensionality of the latent state and dynamics that one can recover based on the hankel matrix constructed out of the observations.

For an LTI system in steady-state, the time independent hankel matrix ($\mathbf{H}$) can be written as:

$$\mathbf{H} \quad = \mathrm{Cov}(\tilde{\mathbf{z}}_t^+, \tilde{\mathbf{z}}_t^-) = \begin{bmatrix} \mathbf{\Lambda}_1 & \mathbf{\Lambda}_2 & \dots & \mathbf{\Lambda}_q \\ \mathbf{\Lambda}_2 & \mathbf{\Lambda}_3 & \dots & \mathbf{\Lambda}_{q+1} \\ \vdots & & \ddots & \vdots \\ \mathbf{\Lambda}_q & \mathbf{\Lambda}_{q+1} & \dots & \mathbf{\Lambda}_{2q-1} \end{bmatrix} \tag{56}$$

where, $\mathbf{\Lambda}_l$ = Cov $(\tilde{\mathbf{Z}}_{t+l}, \tilde{\mathbf{Z}}_t)$. These lagged covariance matrices can then be written as a function of the underlying model parameters. Specifically, $\mathbf{\Lambda}_l$ has the following functional form:

$$\mathbf{\Lambda}_l = \mathrm{Cov}\left(\tilde{\mathbf{Z}}_{t+l}, \tilde{\mathbf{Z}}_t\right) = \mathbf{C}\mathbf{A}^l\mathbf{P}_t\mathbf{C}' \tag{57}$$

where, $\mathbf{P}_t = \mathrm{Cov}(\mathbf{x}_t, \mathbf{x}_t)$ is the zero-lag covariance of the latent state. For an LTI system in steady state, $\mathbf{P}_t$ is typically independent of 't' (stationarity). Given the form of the lagged covariance $\mathbf{\Lambda}_l$ in Equation 57 and assuming steady-state, we can factorize $\mathbf{H}$ as shown below:

$$\mathbf{H} \quad = \underbrace{\begin{bmatrix} \mathbf{C} \\ \mathbf{CA} \\ \mathbf{CA}^2 \\ \vdots \\ \mathbf{CA}^{q-1} \end{bmatrix}}_{\boldsymbol{\mathcal{O}}} \underbrace{\left[\mathbf{APC}' \quad \mathbf{A}^2\mathbf{PC}' \quad \cdots \quad \mathbf{A}^q\mathbf{PC}'\right]}_{\boldsymbol{\Psi}} \tag{58}$$

The matrix $\boldsymbol{\mathcal{O}}$ in Equation 58 is referred to as the *observability* matrix, and is a critical concept in linear systems theory. When $\boldsymbol{\mathcal{O}}$ is full rank, latent states $\mathbf{x}_t$ are uniquely determinable based on observations $\tilde{\mathbf{z}}_t$ alone. By construction, the column space of $\boldsymbol{\mathcal{O}}$ spans the same subspace as the column space of $\mathbf{C}$, which we have referred to alternately as the dynamics subspace (see Methods). Therefore, estimating the column space of $\boldsymbol{\mathcal{O}}$ allows us to obtain an estimate of the dynamics subspace. Based on the relationship between $\boldsymbol{\mathcal{O}}$ and $\mathbf{H}$ in Equation 58, if $\boldsymbol{\mathcal{O}}$ is full rank (i.e latent states are uniquely determinable), then $\boldsymbol{\mathcal{O}}$ and $\mathbf{H}$ span the same column space. Therefore, $\boldsymbol{\mathcal{O}}$ can be obtained empirically through a singular value decomposition of $\mathbf{H}$ because the left singular vectors of $\mathbf{H}$ specify its column space.

$$\mathbf{H} = \mathbf{U}.\mathbf{S}.\mathbf{V}' \tag{59}$$
$$\widehat{\boldsymbol{\mathcal{O}}} = \mathbf{U}_{(\mathrm{r})}.\mathbf{S}_{(\mathrm{r})}{}^{1/2}$$

The particular definition of $\widehat{\boldsymbol{\mathcal{O}}}$ in Equation 58 i.e the first ‘r’ left singular vectors ($\mathbf{U}_{(\mathrm{r})}$) weighted by the square root of their respective singular values ($\mathbf{S}_{(\mathrm{r})}$) is often referred to as the “balanced realization”. This choice of weighting has no bearing on the directions spanned by the dynamics subspace ($\hat{\mathbf{C}}$).

Obtaining an accurate estimate of $\widehat{\boldsymbol{\mathcal{O}}}$ requires determining the rank of $\mathbf{H}$. Typically, for appropriate choices of $q$ ($q \geq p$), the Cayley-Hamilton theorem requires that $\mathbf{H}$ is low rank and that the rank of $\mathbf{H}$ matches the dimensionality of the latent state ($p$). Therefore, the parameter $r$ in Equation 59 specifies the estimated rank of $\mathbf{H}$.

Based on the definition of $\boldsymbol{\mathcal{O}}$ in Equation 58, the estimate of the observation matrix/dynamics subspace ($\hat{\mathbf{C}}$) can then be read off as the first block-row of $\widehat{\boldsymbol{\mathcal{O}}}$.

$$\hat{\mathbf{C}} = \widehat{\boldsymbol{\mathcal{O}}}(1\!:\mathrm{n}, :) \tag{60}$$

We adapted and extended the above framework described for LTI systems to a model characterized with a linear time-varying (LTV) latent dynamical process, by explicitly considering the dependence of the hankel and observability matrices on time. Therefore, in our data analysis pipeline, we build time-dependent hankel matrices $\mathbf{H}_{\mathrm{t}}$, by considering a window of observations (residuals) centred around time ‘t’. These time-dependent hankel matrices then specify time-dependent observability matrices $\boldsymbol{\mathcal{O}}_{\mathrm{t}}$ in a manner similar to Equation 59. However, the explicit dependence of observability matrices on time adds a complication in the estimation of a time-invariant observation matrix, as in the model used to estimate residual dynamics (Equation 11). Therefore, we develop a set of alternate procedures, wherein, we use $\boldsymbol{\mathcal{O}}_{\mathrm{t}}$ to first estimate a ‘momentary’ dynamics subspace using the same definitions as in Equations 58-60 ($\hat{\mathbf{C}}_t$, see Equations 20 and 21) that is time-dependent. We then use the sequence of time-dependent momentary dynamics subspaces estimated across all time to construct a single time-invariant dynamics subspace (Equation 22).

### B. Analytical derivations of residual dynamics for a linear dynamical system with colored input noise

In this section, we derive the effect of temporally correlated (colored) input noise (Fig 1b, complex input) on estimates of residual dynamics. We assumed the linear time-invariant state-space model specified by Equation 47 (see Methods). We assume that we only have access to latent residual states $\mathbf{x}(\mathrm{t})$ through observed residuals $\mathbf{z}(\mathrm{t})$, and that the model operates under steady-state conditions. By defining an augmented latent state, $\mathbf{p}(\mathrm{t})$, we can rewrite Equation 47 as follows:

$$\underbrace{\begin{bmatrix}\mathbf{x}(\mathrm{t}+1)\\ \boldsymbol{\varepsilon}(\mathrm{t}+1)\end{bmatrix}}_{\mathbf{p}(\mathrm{t}+1)} = \underbrace{\begin{bmatrix}\mathbf{A} & \mathbf{I}\\ \mathbf{0} & \boldsymbol{\phi}\end{bmatrix}}_{\widetilde{\mathbf{A}}} \underbrace{\begin{bmatrix}\mathbf{x}(\mathrm{t})\\ \boldsymbol{\varepsilon}(\mathrm{t})\end{bmatrix}}_{\mathbf{p}(\mathrm{t})} + \underbrace{\begin{bmatrix}\mathbf{0}\\ \boldsymbol{\zeta}(t)\end{bmatrix}}_{\boldsymbol{\vartheta}(\mathrm{t})}$$

$$\begin{bmatrix}\mathbf{z}(\mathbf{t})\\ \mathbf{0}\end{bmatrix} = \begin{bmatrix}\mathbf{C} & \mathbf{0}\\ \mathbf{0} & \mathbf{0}\end{bmatrix}\begin{bmatrix}\mathbf{x}(\mathbf{t})\\ \boldsymbol{\varepsilon}(\mathrm{t})\end{bmatrix} + \begin{bmatrix}\boldsymbol{\eta}(\mathrm{t})\\ \mathbf{0}\end{bmatrix} \tag{61}$$

where,

$$\boldsymbol{\vartheta}(\mathrm{t}) \sim \mathcal{N}\left(\begin{bmatrix}\mathbf{0}\\ \mathbf{0}\end{bmatrix}, \begin{bmatrix}\mathbf{0} & \mathbf{0}\\ \mathbf{0} & \mathbf{Q}\end{bmatrix}\right) \tag{62}$$

We define the *l*-lag covariance between random variables **u** and **v** as follows:

$$\mathbf{C}_{\mathrm{ss}}^{(\mathrm{l})}(\mathbf{u},\mathbf{v}) = \mathrm{Cov}(\mathbf{u}(\mathrm{t}+\mathrm{l}), \mathbf{v}(\mathrm{t})) \tag{63}$$

The steady-state zero-lag covariance of the augmented latent state $\mathbf{p}(\mathrm{t})$ can be computed using the discrete-time lyapunov equation for a specified $\widetilde{\mathbf{A}}$ and $\mathbf{Q}$. It corresponds to a block matrix $\mathbf{C}_{\mathrm{ss}}^{(0)}$ which can be written as follows:

$$\begin{aligned}\mathbf{C}_{\mathrm{ss}}^{(0)}(\mathbf{p},\mathbf{p}) = \mathrm{Cov}\big(\mathbf{p}(\mathrm{t}), \mathbf{p}(\mathrm{t})\big) &= \begin{bmatrix}\mathrm{Cov}\big(\mathbf{x}(\mathrm{t}), \mathbf{x}(\mathrm{t})\big) & \mathrm{Cov}\big(\mathbf{x}(\mathrm{t}), \boldsymbol{\varepsilon}(\mathrm{t})\big)\\ \mathrm{Cov}\big(\boldsymbol{\varepsilon}(\mathrm{t}), \mathbf{x}(\mathrm{t})\big) & \mathrm{Cov}\big(\boldsymbol{\varepsilon}(\mathrm{t}), \boldsymbol{\varepsilon}(\mathrm{t})\big)\end{bmatrix}\\ &= \begin{bmatrix}\mathbf{C}_{\mathrm{ss}}^{(0)}(\mathbf{x},\mathbf{x}) & \mathbf{C}_{\mathrm{ss}}^{(0)}(\mathbf{x},\boldsymbol{\varepsilon})\\ \mathbf{C}_{\mathrm{ss}}^{(0)}(\boldsymbol{\varepsilon},\mathbf{x}) & \mathbf{C}_{\mathrm{ss}}^{(0)}(\boldsymbol{\varepsilon},\boldsymbol{\varepsilon})\end{bmatrix}\end{aligned} \tag{64}$$

The steady-state *lag-l* covariance of the augmented state $\mathbf{p}(\mathrm{t})$ is therefore defined as:

$$\mathbf{C}_{\mathrm{ss}}^{(\mathrm{l})}(\mathbf{p},\mathbf{p}) = \mathrm{Cov}(\mathbf{p}(\mathrm{t}+\mathrm{l}), \mathbf{p}(\mathrm{t})) = \widetilde{\mathbf{A}}^{\boldsymbol{l}}\mathbf{C}_{\mathrm{ss}}^{(0)}(\mathbf{p},\mathbf{p}) \tag{65}$$

Now, the exponentiated dynamics matrix $\widetilde{\mathbf{A}}^{l}$ in the above equation has the following general form:

$$\widetilde{\mathbf{A}}^{l} = \begin{bmatrix}\mathbf{A}^{l} & \mathbf{U}(l)\\ \mathbf{0} & \boldsymbol{\phi}^{l}\end{bmatrix} \tag{66}$$

where,

$$\mathbf{U}(l) = \sum_{k=0}^{l-1} \mathbf{A}^k \boldsymbol{\phi}^{\,l-1-k} \tag{67}$$

Using Equations 64, 66 and 67, Equation 65 can therefore be expanded as follows:

$$\begin{aligned} \mathbf{C}_{\mathrm{ss}}^{(l)}(\mathbf{p},\mathbf{p}) &= \widetilde{\mathbf{A}}^{l}\mathbf{C}_{\mathrm{ss}}^{(0)}(\mathbf{p},\mathbf{p}) = \begin{bmatrix} \mathbf{A}^l & \mathbf{U}(l) \\ \mathbf{0} & \boldsymbol{\phi}^l \end{bmatrix} \begin{bmatrix} \mathbf{C}_{\mathrm{ss}}^{(0)}(\mathbf{x},\mathbf{x}) & \mathbf{C}_{\mathrm{ss}}^{(0)}(\mathbf{x},\boldsymbol{\varepsilon}) \\ \mathbf{C}_{\mathrm{ss}}^{(0)}(\boldsymbol{\varepsilon},\mathbf{x}) & \mathbf{C}_{\mathrm{ss}}^{(0)}(\boldsymbol{\varepsilon},\boldsymbol{\varepsilon}) \end{bmatrix} \\ &= \begin{bmatrix} \mathbf{A}^l\mathbf{C}_{\mathrm{ss}}^{(0)}(\mathbf{x},\mathbf{x}) + \mathbf{U}(l)\mathbf{C}_{\mathrm{ss}}^{(0)}(\mathbf{x},\boldsymbol{\varepsilon}) & \mathbf{A}^l\mathbf{C}_{\mathrm{ss}}^{(0)}(\mathbf{x},\boldsymbol{\varepsilon}) + \mathbf{U}(l)\mathbf{C}_{\mathrm{ss}}^{(0)}(\boldsymbol{\varepsilon},\boldsymbol{\varepsilon}) \\ \boldsymbol{\phi}^l\mathbf{C}_{\mathrm{ss}}^{(0)}(\boldsymbol{\varepsilon},\mathbf{x}) & \boldsymbol{\phi}^l\mathbf{C}_{\mathrm{ss}}^{(0)}(\boldsymbol{\varepsilon},\boldsymbol{\varepsilon}) \end{bmatrix} \end{aligned} \tag{68}$$

Therefore, based on Equation 68, the lag-*l* steady state covariance of the latent state $\mathbf{x}$ of Equation 47 can be read off as the upper left block of $\mathbf{C}_{\mathrm{ss}}^{(l)}(\mathbf{p},\mathbf{p})$ as below:

$$\mathbf{C}_{\mathrm{ss}}^{(l)}(\mathbf{x},\mathbf{x}) = \mathbf{A}^l\mathbf{C}_{\mathrm{ss}}^{(0)}(\mathbf{x},\mathbf{x}) + \mathbf{U}(l)\mathbf{C}_{\mathrm{ss}}^{(0)}(\mathbf{x},\boldsymbol{\varepsilon}) \tag{69}$$

The expression in Equation 69 can be used to build a two stage least squares (2SLS) estimator. As outlined in the Methods, the first stage of the 2SLS regression, involves regressing the latent state $\mathbf{x}$(t) against its past in order to build a predictor $\hat{\mathbf{x}}$(t). In the second stage, the predictor $\hat{\mathbf{x}}$(t) is regressed against $\mathbf{x}$(t+1) in order to estimate the dynamics matrix. Assuming, that we use *m* past lags for the first-stage of the 2SLS, we constructed the following three 'stacked' multi-lag covariance matrices using $\mathbf{C}_{\mathrm{ss}}^{(\mathrm{l})}(\mathbf{x},\mathbf{x})$

$$\begin{aligned} \mathbf{P}_0 &= \begin{bmatrix} \mathbf{C}_{\mathrm{ss}}^{(0)}(\mathbf{x},\mathbf{x}) & \mathbf{C}_{\mathrm{ss}}^{(1)}(\mathbf{x},\mathbf{x}) & \dots & \mathbf{C}_{\mathrm{ss}}^{(\mathrm{m}-1)}(\mathbf{x},\mathbf{x}) \\ \mathbf{C}_{\mathrm{ss}}^{(-1)}(\mathbf{x},\mathbf{x}) & \mathbf{C}_{\mathrm{ss}}^{(0)}(\mathbf{x},\mathbf{x}) & \dots & \mathbf{C}_{\mathrm{ss}}^{(\mathrm{m}-2)}(\mathbf{x},\mathbf{x}) \\ \vdots & & \ddots & \vdots \\ \mathbf{C}_{\mathrm{ss}}^{(-\mathrm{m}-1)}(\mathbf{x},\mathbf{x}) & \mathbf{C}_{\mathrm{ss}}^{(-\mathrm{m}-2)}(\mathbf{x},\mathbf{x}) & \dots & \mathbf{C}_{\mathrm{ss}}^{(0)}(\mathbf{x},\mathbf{x}) \end{bmatrix} \\ \mathbf{P}_1 &= \begin{bmatrix} \mathbf{C}_{\mathrm{ss}}^{(1)}(\mathbf{x},\mathbf{x}) & \mathbf{C}_{\mathrm{ss}}^{(2)}(\mathbf{x},\mathbf{x}) & \dots & \mathbf{C}_{\mathrm{ss}}^{(\mathrm{m})}(\mathbf{x},\mathbf{x}) \end{bmatrix} \\ \mathbf{P}_2 &= \begin{bmatrix} \mathbf{C}_{\mathrm{ss}}^{(2)}(\mathbf{x},\mathbf{x}) & \mathbf{C}_{\mathrm{ss}}^{(3)}(\mathbf{x},\mathbf{x}) & \dots & \mathbf{C}_{\mathrm{ss}}^{(\mathrm{m}+1)}(\mathbf{x},\mathbf{x}) \end{bmatrix} \end{aligned} \tag{70}$$

where, $\mathbf{C}_{\mathrm{ss}}^{(-\mathrm{k})}(\mathbf{x},\mathbf{x}) = \mathbf{C}_{\mathrm{ss}}^{(\mathrm{k})}(\mathbf{x},\mathbf{x})^{\mathrm{T}}$. The two-stage least squares estimator based on *m* past lags is then given by:

$$\widehat{\mathbf{A}}_{2\mathrm{sls}}^{(\mathrm{m})} = \left(\mathbf{P}_2.(\mathbf{P}_0{}^{-1})^{\mathrm{T}}\mathbf{P}_1{}^{\mathrm{T}}\right)\left(\mathbf{P}_1.(\mathbf{P}_0{}^{-1})^{\mathrm{T}}\mathbf{P}_1{}^{\mathrm{T}}\right)^{-1} \tag{71}$$

As in the data, we use a lag of m = 3, in order to construct the 2SLS analytical estimator of the residual dynamics in Extended Data Fig. 4. The pseudocode for constructing the estimator is given below:

**Algorithm 1:** Compute analytical 2SLS estimator for linear-SSM with correlated input noise

*INPUT:* $\mathbf{A}, \boldsymbol{\phi}, \mathbf{Q}, m$

*OUTPUT:* $\widehat{\mathbf{A}}_{2\mathrm{sls}}^{(\mathrm{m})}$

1. Construct $\widetilde{\mathbf{A}} = \begin{bmatrix} \mathbf{A} & \mathbf{I} \\ \mathbf{0} & \boldsymbol{\phi} \end{bmatrix}$ and $\widetilde{\mathbf{Q}} = \begin{bmatrix} \mathbf{0} & \mathbf{0} \\ \mathbf{0} & \mathbf{Q} \end{bmatrix}$
2. Solve for $\mathbf{C}_{\mathrm{ss}}^{(0)}(\mathbf{p},\mathbf{p})$ : $\widetilde{\mathbf{A}}\,\mathbf{C}_{\mathrm{ss}}^{(0)}\widetilde{\mathbf{A}}^{\boldsymbol{T}} - \mathbf{C}_{\mathrm{ss}}^{(0)} + \widetilde{\mathbf{Q}} = \mathbf{0}$ (discrete-time lyapunov equation)
3. Use individual block entries of $\mathbf{C}_{\mathrm{ss}}^{(0)}(\mathbf{p},\mathbf{p})$, $\mathbf{A}$ and $\boldsymbol{\phi}$ to define $\mathbf{C}_{\mathrm{ss}}^{(l)}(\mathbf{x},\mathbf{x})$ for $l$ = 0,1,2 … m+1 (Equation 69)
4. Define $\mathbf{P}_0$, $\mathbf{P}_1$ and $\mathbf{P}_2$ using $\mathbf{C}_{\mathrm{ss}}^{(l)}(\mathbf{x},\mathbf{x})$ as in Equation 70
5. Compute and return $\widehat{\mathbf{A}}_{2\mathrm{sls}}^{(\mathrm{m})}$ using output of step 4 in Equation 71
